# Evolutionary Origin of Vertebrate OCT4/POU5 Functions in Supporting Pluripotency

**DOI:** 10.1101/2022.04.13.488202

**Authors:** Woranop Sukparangsi, Elena Morganti, Molly Lowndes, Hélène Mayeur, Melanie Weisser, Fella Hammachi, Hanna Peradziryi, Fabian Roske, Jurriaan Hölzenspies, Alessandra Livigni, Benoit Gilbert Godard, Fumiaki Sugahara, Shigeru Kuratani, Guillermo Montoya, Stephen R. Frankenberg, Sylvie Mazan, Joshua M Brickman

**Author notes:** **Corresponding authors :** Joshua M Brickman and Sylvie Mazan. These authors contributed equally to this work (Woranop Sukparangsi, Elena Morganti).

## Abstract

The support of pluripotent cells over time is an essential feature of development. In eutherian embryos, pluripotency is maintained from naïve states in peri-implantation to primed pluripotency at gastrulation. To understand how these states emerged, we reconstruct the evolutionary trajectory of the *Pou5* gene family, which contains the central pluripotency factor OCT4. By coupling evolutionary sequence analysis with functional studies in mouse Embryonic Stem Cells (ESCs), we found that the ability of POU5 proteins to support pluripotency originated in the gnathostome lineage, prior to the generation of two paralogues, *Pou5f1* and *Pou5f3* via gene duplication. In osteichthyans, retaining both genes, the paralogues differ in their support of naïve and primed pluripotency. This specialization of these duplicates enables the diversification of function in self-renewal and differentiation. By integrating sequence evolution, ESC phenotypes, developmental contexts and structural modelling, we pinpoint OCT4 regions sufficient for naïve pluripotency and describe their adaptation over evolutionary time.

## Introduction

Pluripotency refers to the capacity of a cell to give rise to all lineages of the adult body, including the germ line. This functional property was historically defined based on the advent of mouse Embryonic Stem Cells (ESCs), which made the mouse the reference model to define and explore the molecular basis for pluripotency. As the number of culture models expanded, it became clear that pluripotent cells exist across a range of cell states and developmental windows. In mammals, pluripotent cells can be found throughout distinct developmental stages *in vivo*, transitioning from an initial naïve state to a lineage primed one as development progresses from pre-implantation stages to gastrulation (reviewed in ref^1^). In mouse, these two states can be captured and cells can be expanded *ex vivo* in well-defined culture conditions. Mouse ESCs represent a naïve pluripotent state and their gene expression pattern approximates that of the Inner Cell Mass (ICM) of pre-implantation embryos. Mouse Epiblast Stem Cells (EpiSCs) represent a primed pluripotent state, which is more reminiscent of later pre-gastrulation stages of development^2,3^. The regulation of these pluripotent states has been extensively investigated and involves the input of extrinsic signals into a complex network of transcription factors. While naïve and primed cells share expression of a number of transcription factors, including OCT4 (POU5F1), SOX2 and NANOG, the transition from a naïve to primed state involves major changes in embryonic environment, transcriptomic profile (with the down-regulation and up-regulation of state or stage specific pluripotency regulators such as *Esrrb*, *Prdm15* or *FoxD3*) and enhancer or chromatin landscapes^4–12^. These molecular changes parallel a remodelling of embryo architecture, including epithelialisation and generation of the amniotic cavity^13,14^.

While the functional definition of pluripotency is unique to mammals, the concept of pluripotent populations is central to all developmental biology. Even with a plethora of mechanistic information characterising pluripotent states in the mouse, there is a scarcity of data on their evolutionary origin and conservation across vertebrates. ESCs exhibiting either naïve or primed pluripotency have been obtained in humans and other primates^15–20^, but a clear set of distinct cell types has yet to be defined in marsupials and monotremes^21,22^. Similarly, the existence of a naïve pluripotent state in the zebrafish embryo, based on early expression dynamics of a selection of pluripotency markers, remains hypothetical in the absence of established cell lines exhibiting ESC properties^23^. Altogether, the existence of naïve and primed pluripotent states, as extensively described in the mouse, remains unclear outside eutherians. An alternative approach to investigate their origin is to deconstruct their evolutionary trajectory, analysing when the capacity of key members of the pluripotency network to support these states emerged during evolution. We have used this approach, focusing on class V POU domain (POU5) transcription factors (OCT4 in the mouse) at key nodes of the vertebrate tree. This small multigene family comprises two orthology classes, *Pou5f1* and *Pou5f3*, in jawed vertebrates (gnathostomes)^24,25^. While a number of key nodes in gnathostome evolution retain both genes, why a single paralogue is retained in many vertebrate species remains a mystery. In eutherians, *Oct4*, which belongs to the *Pou5f1* class, is the only representative of the gene family and is a central regulator of pluripotency both *in vivo* and *in vitro*. It is absolutely required to establish and maintain pluripotency in all contexts, but depending on expression levels, it can also mediate differentiation into distinct embryonic lineages^26–29^. This functional complexity is confirmed by *in vivo* analyses, with distinct roles for this factor depending on both stage and cellular context. Prior to implantation, from early to late blastocyst stages, OCT4 is first required to maintain the ICM and inhibit trophoblast differentiation, then for specification of both primitive endoderm (PrE) and epiblast^30–32^. At later stages, the loss of OCT4 from post-implantation epiblast results in multiple abnormalities, including a general disorganisation of germ layers, impaired expansion of the primitive streak and apoptosis of primordial germ cells (PGCs)^26,33–34^. In primed pluripotent cells, *in vitro*, the immediate phenotype in response to inducible removal of OCT4 is a loss of E-cadherin (CDH1) and impaired adhesion^35^. Thus, mouse OCT4 is required to regulate pluripotency by both supporting self-renewal and establishing competence for differentiation. It is also at the heart of both primed and naïve pluripotency networks, but acting to regulate different sets of enhancers in these distinct pluripotent states^5^. While naïve pluripotency concerns pre-implantation and appears specific to eutherian mammals, there is support for a conserved POU5 dependent network regulating aspects of pluripotency in other species. Evidence for a conserved role of POU5s in the control of pluripotency has been obtained in frog (*Xenopus*), chick, axolotl and teleosts^36–41^. Similarly, the knock-down of OCT4 homologues in *Xenopus* and zebrafish lead to gastrulation phenotypes reminiscent of those observed in the mouse and similarly related to impaired cell adhesiveness^35,42^. In these species, which have lost the *Pou5f1* class, all pluripotency related functions are fulfilled by POU5F3 rather than POU5F1.

A phenotypic complementation, or rescue assay, for OCT4 has been developed in mouse ESCs, providing a means to evaluate the ability of heterologous POU5 proteins to substitute for OCT4 in the support of naïve pluripotency and in the control of the balance between self-renewal and differentiation^43^. POU5 proteins from different species exhibit varying abilities to rescue in this assay, irrespective of the orthology class. For instance, human, platypus and axolotl POU5F1s, as well as *Xenopus* XlPOU91 (XlPOU5F3.1), one of the three POU5F3 forms identified in this species, are endowed with a similar rescue capacity, indicating that they harbour essential structural determinants required to support naïve pluripotency in mouse ESCs. In contrast, moderate or undetectable rescue ability was observed for chick and zebrafish POU5F3, respectively^36,38^. The existence of homologues with varying OCT4-like activity suggests that the role of this factor in pluripotency has undergone functional diversification across vertebrates. Here, we take advantage of the OCT4 complementation system to explore when POU5 proteins acquired the capacity to fulfil mouse OCT4 functions and how they evolved in the context of the duplication that gave rise to the POU5F1 and POU5F3 forms. Our results indicate that the capacity of POU5 proteins to support naïve pluripotency is a gnathostome characteristic, which emerged prior to the duplication giving rise to the *Pou5f1* and *Pou5f3* orthology classes and was elaborated in the sarcopterygian lineage. This was a result of a stepwise process, involving specialisations of the two paralogues impacting the structural orientation of two regions of the protein that allowed neo-functionalisation and reversion. Altogether, these data unveil an ancient evolutionary history for pluripotency that suggests that the states, extensively analysed in eutherians, existed long before the advent of placental development.

## Results

### Evolutionary dynamics of the *Pou5* gene family in vertebrates

Previous characterisation of *Pou5f1* and *Pou5f3* has highlighted multiple losses of either one of the two paralogues in many osteichthyan (including tetrapod) lineages^24^. To explore the evolutionary dynamics of the *Pou5* gene family across vertebrates, we performed a comprehensive survey of these genes in a broad sampling of vertebrates, taking advantage of available genomic databases (Supplementary Table 1). Deduced amino acid sequences were submitted to sequence comparisons, phylogenetic and synteny analyses (Fig. 1a-c, Supplementary Information 1). All vertebrate full-length coding sequences predicted from our genomic searches exhibited a very similar organisation into five coding exons, with conserved locations of intron-exon boundaries, albeit with a reverse order between exons 4-5 and 2-3 in *E. burgeri*, possibly related to a genome assembly error (Supplementary Table 1). Their assignment to the *Pou5* gene family is supported by the high level of conservation of the POU specific domain and POU homeodomain with residues identified as POU5 synapomorphies (L146^(POU16)^, K149^(POU19)^, C245^(POU115)^, Supplementary Information 1; ref^44^) and the presence of a N-terminal motif shared by all POU5 proteins (Supplementary information 1). While sequence comparisons highlight a few signature residues of osteichthyan POU5F1 and POU5F3 in the POU specific domain and homeodomain (residue D/E at D205^(POU75)^ and residue -/R between K226^(POU96)^ – R227^(POU97)^; Fig. 1a; ref^24^), these candidate class hallmarks are not maintained in orthologous chondrichthyan sequences, suggesting the fixation of novel selective constraints in the osteichthyan lineage (Fig. 1a). Lamprey and hagfish POU5 share a number of residues not found in their gnathostome counterparts, supporting the monophyly of cyclostome POU5 (Supplementary Information 1).

**Fig. 1.**
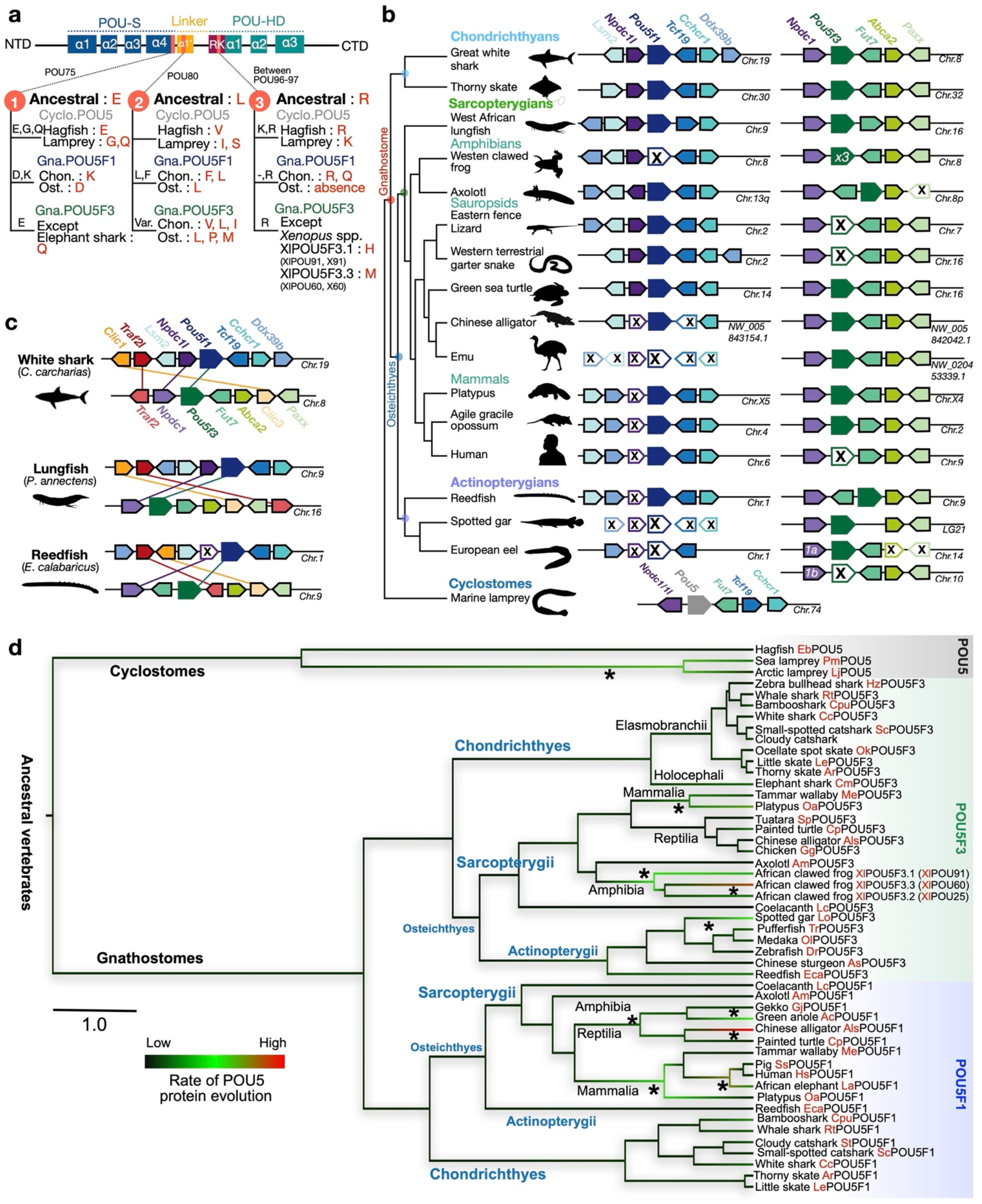
Evolution of the POU5 family in vertebrates. **a**, Differences between gnathostome POU5F1 and POU5F3 proteins in the POU-S-Linker-POU-HD. Ancestral residues prior to the duplication generating the gnathostome paralogues are shown in bold characters for positions POU75, POU80 and POU96/97 of POU5F1 (Gna.POU5F1, blue) and POU5F3 (Gna.POU5F3, green). Residues found in cyclostomes (Cyclo.POU5, grey) at these positions are also shown. NTD and CTD stand for N-terminal and C-terminal domains, respectively. **b**, Synteny conservations in the vicinity of *Pou5* genes across vertebrates. *Pou5* genes are depicted by coloured arrows, with a single arrow for the three *X. laevis Pou5f3* replicates and two arrows representing the combinations of exons found in inverted order in hagfish *E. burgeri* (see Results and Supplementary Information 1). Black crosses indicate a missing gene in the genomic data analysed. In gnathostomes, genes showing conserved syntenies with *Pou5f1* (in blue) and *Pou5f3* (in green) are respectively shown in left and right panels. In lampreys, the single homologue found for *Pou5* is shown in teal below its gnathostome counterparts. It shares conserved syntenies both with gnathostome *Pou5f1* (*Tcf19*, *Chcr1*) and *Pou5f3* (*Fut7*) loci. Species names are as follows: great white shark, *Carcharodon carcharias*; thorny skate, *Amblyraja radiata*; African lungfish, *Protopterus annectens*; African clawed frog, *Xenopus laevis*; Eastern fence lizard, *Sceloporus undulatus*; Western terrestrial garter snake, *Thamnophis elegans*; green sea turtle, *Chelonia mydas*; Chinese alligator, *Alligator sinensis*; Emu, *Dromaius novaehollandia*; Platypus, *Ornithorhynchus anatinus*; agile opossum, *Gracilinanus agilis*; human, *Homo sapiens*; reedfish, *Erpetoichthys calabaricus*; spotted gar, *Lepisosteus oculatus*; European eel, *Anguilla anguilla* **c**, Conserved syntenies between pairs of paralogous genes (*Clic1*/*Clic3*, *Traf2*/*Traf2* and *Npdc1*/*Npdc1l* respectively in yellow, red and purple arrows) found in the vicinity of gnathostome *Pou51* and *Pou53* genes in the great white shark, lungfish and reedfish. These syntenies are broadly conserved in chondrichthyans, sarcopterygians and actinopterygians (see Supplementary Information 1). **d**, Phylogenetic tree showing the evolutionary rates of cyclostome POU5, gnathostome POU5F1 and POU5F3 sequences in different vertebrate lineages. Evolutionary rates were calculated from the alignment of POU specific domain, linker and homeodomain (as shown in the Supplementary Dataset 1) using BEAST, by imposing the monophyly of gnathostome POU5F1 and POU5F3 and the species phylogeny within these groups. They are represented along tree branches in black, green to red from low, moderate to high. Asterisks show branches in which accelerations of evolutionary rates have taken place. The scale bar corresponds to the average number of amino acid changes per site. Species name abbreviations are listed in Supplementary Information 1.

In osteichthyans, *Pou5f1* and/or *Pou5f3*-related genes can be unambiguously identified in all species analysed. Furthermore, this analysis confirmed a complex pattern of paralogue loss/retention: (i) the presence of both forms in the last common ancestor of sarcopterygians (e.g. lungfish), (ii) independent *Pou5f1* losses/*Pou5f3* retention in actinopterygians (except reedfish), anurans (e.g. frog) and birds (e.g. emu) and (iii) independent *Pou5f3* losses/*Pou5f1* retention in eutherians (e.g. human and mouse) and squamates (e.g. lizard and snake) (Fig. 1b and Supplementary Information 1). It also resolves the timing of paralogue loss/retention events with an increased resolution. For instance, we identified an unambiguous *Pou5f3* coding sequence in the tuatara *Sphenodon punctatus* (Supplementary Table 1), which implies that the loss of this paralogue in squamates followed their split from sphenodonts. Similarly, both *Pou5f1* and *Pou5f3* can be identified in the genome of *Alligator sinensis* (Fig. 1b), in line with a retention of both paralogues not only in turtles as previously documented^24^, but also in archosaurs, their sister group, prior to the loss of *Pou5f1* in birds. In actinopterygians, both paralogues are present in the reedfish *Erpechtoichtys calabaricus*, implying that the loss of *Pou5f1* previously documented in this group followed the split between cladistians and actinopteri (Fig. 1b). In all chondrichthyans (cartilaginous fishes) analysed, we obtained robust evidence for the presence of both paralogues with full length coding sequences found in elasmobranchs, including sharks (small-spotted catshark *Scyliorhinus canicula*, white shark *Carcharodon carcharias*, brownbanded bamboo shark *Chiloscyllium punctatum*, whale shark *Rhincodon typus*) and skates (little skate *Leucoraja erinacea* and thorny skate *Amblyraja radiata*), as well as full length *Pou5f3* and partial *Pou5f1* sequences in the holocephalan *Callorhinchus milii* (Supplementary Table 1; Supplementary Information 1). Finally, in cyclostomes, searches in the genomes of two lampreys, *Lethenteron reissneri* and *Petromyzon marinus*, and in the genome of the hagfish *Eptratetus burgeri* indicated the presence of only one *Pou5*-related coding sequence in each one of these species. These coding sequences could not be assigned to either one of the gnathostome POU5F1 or POU5F3 classes based on amino acid sequence comparisons or phylogenetic analysis (Supplementary Table 1; Supplementary Information 1).

Synteny analyses show that gnathostome *Pou5f1* and *Pou5f3* are respectively located in conserved chromosomal environments, containing in their close vicinity orthologues of *Lsm2*, *Tcf19*, *Cchcr1*, *Ddx39b* for *Pou5f1*, and of *Fut7*, *Abca2*, *Paxx* for *Pou5f3* (Fig. 1b; Supplementary Information 1). Three pairs of paralogues shared between the *Pou5f1* and *Pou5f3* loci (*Clic1/Clic3*; *Traf2l/Traf2*; *Npdc1l/Npdc1*; Fig. 1c) and retained in chondrichthyans, actinopterygians and sarcopterygians can additionally be detected at higher chromosomal distances, in line with their presence in the ancestral locus prior to the duplication generating the two forms (Fig. 1c; Supplementary Information 1). The chromosomal environment of the unique *Pou5* gene identified in the lamprey species shares characteristics of both gnathostome *Pou5f1* and *Pou5f3* loci, including conserved linkages with *Tcf19*/*Cchcr1* and *Fut7* homologues as observed in the vicinity of gnathostome *Pou5f1* and *Pou5f3* genes respectively (Fig. 1b). Taken together, these data highlight the fixation of significant differences between the gnathostome *Pou5f1* and *Pou5f3* genes following the duplication, which generated them.

### Heterogeneous evolutionary rates of POU5 proteins across vertebrates

In order to gain insight into the molecular constraints acting on POU5 protein sequences, we characterised their evolutionary rate variations using a bayesian Markov chain Monte Carlo algorithm. We first focused on the POU domain (containing the POU specific, linker and homeodomain) in a broad sampling of vertebrates, containing all the cyclostome and chondrichthyan sequences available and a representative sampling of osteichthyans, including teleosts, amphibians, sauropsids and mammals (Fig. 1d). This analysis indicates the occurrence of the most pronounced evolutionary rate accelerations in the branches of lamprey POU5 (after their splitting from hagfish), both mammalian and reptilian POU5F1, mammalian POU5F3 (but not sauropsid POU5F3) and the triplicate *Xenopus* POU5F3 proteins (but not their single copy counterpart in salamander). A remarkably high evolutionary rate is also observed in crocodiles for POU5F1, prior to its loss in birds (Fig. 1d). This analysis was refined for mammalian POU5F1 and actinopterygian POU5F3, using more exhaustive species sampling in these taxa and taking into account the C-terminal part of the protein in addition to the POU specific domain and homeodomain (Supplementary Information 1). In the former, higher POU5F1 evolutionary rates are observed in the lineage of therians versus monotremes and in the lineage of eutherians versus marsupials. Heterogeneities are also detected across eutherians, for example, relatively high evolutionary rates in Murinae (mouse and rats) and most rodents, as well as in Chiroptera (bats). Concerning POU5F3, an acceleration in evolutionary rate is detected early in the actinopterygian lineage, with a higher rate of evolution in the actinopterygian versus sarcopterygian (represented by a crossopterygian, coelacanth POU5F3) branch, as well as in the neopterygian (versus chondrostean) lineage. Such an acceleration may explain the reduced capacity of zebrafish POU5F3 to support OCT4-null mouse ESCs^37^. This heterogeneity in evolutionary rates observed in teleosts is unlikely to be related to hidden paralogy in the context of the whole genome duplication known to have occurred early in the teleost lineage^45^. Both copies generated by the teleost-specific duplication of *Traf2*, *Npdc1* and *Fut7* have been retained in the teleosts analysed and, in all cases, the unique *Pou5f3* gene ed lies in synteny with the same paralogues (*Traf2b*, *Npdc1a*, *Fut7a*). In summary, we recurrently observe significant increases in evolutionary rates associated with paralogue gains and losses, suggesting modifications of the functional constraints acting on coding sequences. However, analysis of non-synonymous to synonymous substitution failed to reveal evidence for protein positive evolution, possibly due to the globally very high conservation of the POU specific domain and POU homeodomain.

### The functional capacity of different sarcopterygian POU5 paralogues to rescue OCT4-null ESCs

To explore the functional evolution of POU5F1 and POU5F3 when both genes are retained, we asked whether both paralogous proteins were able to support naïve pluripotency in a heterologous mouse OCT4-rescue assay. We first focused on sarcopterygians and examined the activities of POU5 proteins from a representative sampling of species that carry both paralogues: the coelacanth (*Latimeria chalumnae*), the axolotl (*Ambystoma mexicanum*), the turtle (*Chrysemys picta bellii*) and the tammar wallaby (*Macropus eugenii*). To better visualize evolutionary trends of POU5 activity in sarcopterygians, POU5s from species that have lost either *Pou5f1* or *Pou5f3* were included, African-clawed frog (*Xenopus laevis*) and python (*Python molurus*) (Fig. 2a). Among the three *Pou5f3* paralogues produced by tandem gene duplications in the frog, only two (*XlPou5f3.1* and *XlPou5f3.2,* encoding for X91 and X25 proteins, respectively) were analysed, as the third one (*XlPou5f3.3; X60*) is dispensable for normal development (Fig. 2a). To assess POU5 activity in supporting pluripotency, we used an *Oct4*^-/-^ mouse ESC line carrying a tetracycline (Tc)-suppressible *Oct4* transgene (ZHBTc4)^27^. We introduced cDNAs encoding heterologous POU5 proteins (coding sequences listed in Supplementary Table 1) into ZHBTc4 cells (in the presence or absence of tetracycline) and determined the rescue potential relative to a mouse OCT4 (mOct4) cDNA control (Fig. 2b). Upon OCT4 loss, ESCs differentiate towards trophoblast, while OCT4 over-expression (when both heterologous cDNA and the *Oct4* transgene are expressed simultaneously) induces differentiation towards extra-embryonic mesoderm and endoderm^27^. With the OCT4-rescue assay we can assess the capacity of heterologous proteins to support an undifferentiated ESC phenotype in the absence of mOct4, as well as the capacity to induce differentiation when expressed in the presence of mOct4 (over-expression). The degree to which a particular POU5 rescued mOct4 activity was assessed based on a colony formation assay, comparing the number of alkaline phosphatase positive colonies (AP^+^; purple) in the presence versus the absence of tetracycline (rescue index) (Fig. 2c, upper panel).

**Fig. 2.**
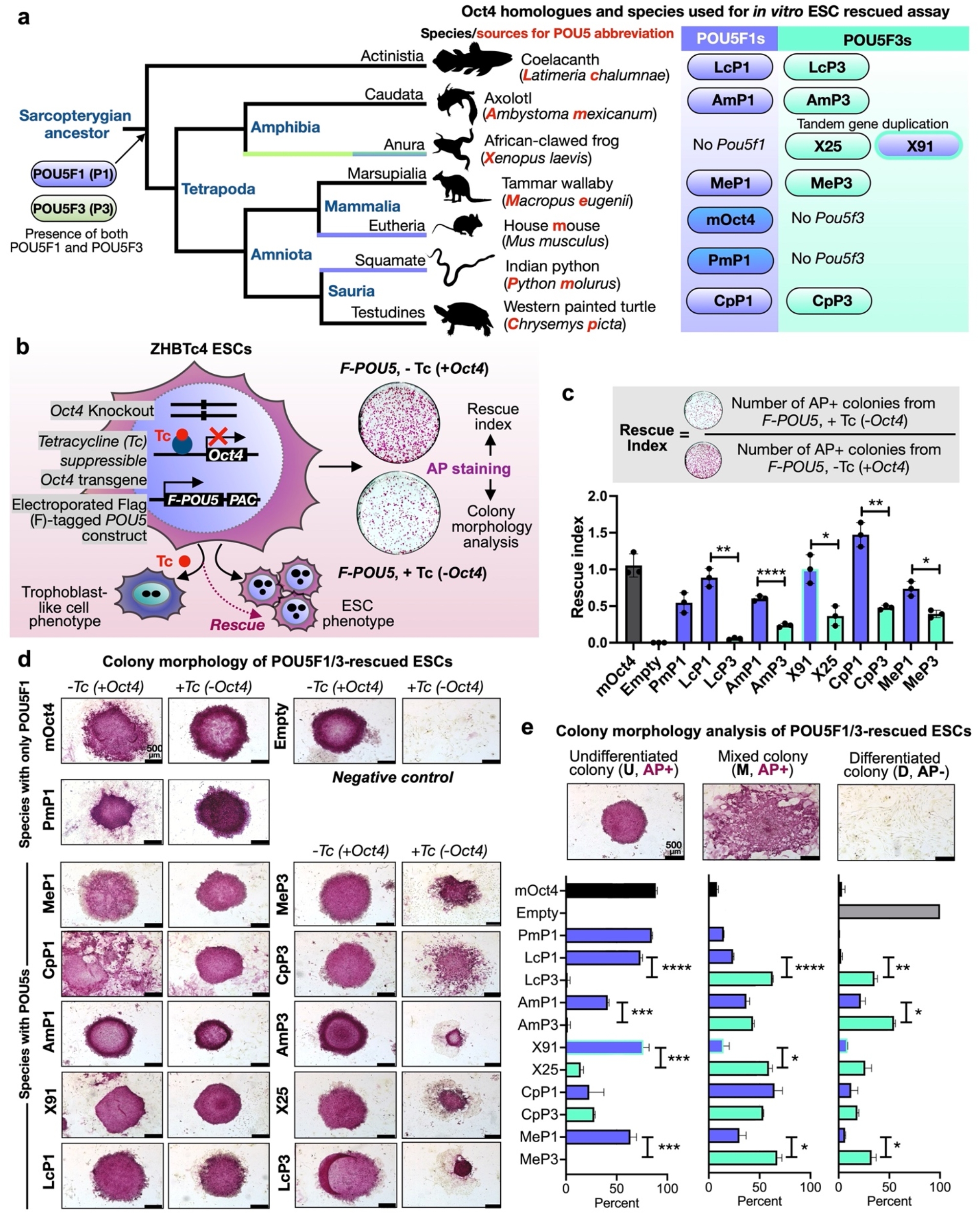
Sarcopterygian POU5F1 proteins have greater capacity to rescue mouse OCT4-null ESC cells than their POU5F3 paralogues. **a**, Schematic illustration showing simplified phylogenetic tree of Sarcopterygian species used for testing POU5 protein activities and introduction of POU5 abbreviations used in this study (letters in red). **b**, Experimental strategy (rescue assay) used to test the capacity of exogenous POU5 proteins (from different vertebrate species) to rescued ZHBTc4 ESCs pluripotency and self-renewal capacity, upon the addition of Tetracycline (Tc). **c**, Rescue indices indicating the capacity of different POU5 homologues to support ESC self-renewal. **d**, Colony phenotypes obtained from POU5-transfected OCT4-null ESC cells grown in the presence or absence of mouse Oct4 (−/+ Tc) at clonal density and stained for Alkaline Phosphatase (AP) activity (purple). **e**, Classification and quantification of ESC colony phenotypes from rescue assay. Colonies were scored as undifferentiated (U), mixed (M), differentiated (D) and AP positive/negative (+/-) colonies. Statistical analyses (Unpaired t-Test) were performed on n=3 biological replicates (3 independent experiments). Error bars indicate standard deviation.

We found that all POU5F1 orthologues from species with either one or two POU5 homologues could rescue OCT4-null ESCs, producing both high levels of undifferentiated colonies (AP^+^) and high rescue indices (Fig. 2c-d, Supplementary Fig. 1a-d). In contrast, the colonies produced by any of the POU5F3 orthologues, except X91, had varied morphologies and overall lower rescue indices (Fig. 2c-d, Supplementary Fig. 1a-b). The majority of POU5F3-rescued colonies retained an undifferentiated centre (AP^+^) surrounded by unstained differentiated cells (Fig. 2d). Quantification of the distinct morphologies produced by these POU5-rescued colonies shows that all POU5F1 proteins produced high percentages of undifferentiated colonies, while POU5F3 proteins supported high numbers of mixed and differentiated colonies (Fig. 2e). Taken together, these observations support the notion that sarcopterygian POU5 paralogues evolved distinct abilities to support pluripotency and self-renewal.

### POU5F1 and POU5F3 support distinct ESC phenotypes

To understand the differences between ESCs supported by the different POU5 proteins, we generated stable cell lines from either POU5F1- or POU5F3-rescued colonies (strategy summarised in Fig. 3a) and confirmed that all cell lines were maintained solely by the heterologous POU5s (Fig. 3b, Supplementary Fig. 2a-c, e). After several passages, almost all clonal lines supported by POU5F1 showed sustained self-renewal and expanded better than those supported by POU5F3 (Supplementary Fig. 2d). POU5F1-rescued ESCs resembled mOct4-rescued controls with homogenous E-cadherin (CDH1) expression and the majority of cells KLF4-positive (Fig. 3b-c). In contrast, POU5F3-rescued ESCs showed mixed morphologies (except for frog X91), with cells expressing either trophectoderm (TE; CDX2^+^) or primitive endoderm (PrE; GATA6^+^) markers (Fig. 3b-c). Moreover, ESCs supported by coelacanth, axolotl or tammar wallaby POU5F3s were prone to differentiate toward TE, while frog X25-rescues differentiated toward PrE and turtle POU5F3-rescues toward both TE and PrE (Fig. 3b-c). Consistent with our previous observations^35,38^, frog X91-rescues were indistinguishable from those supported by mOct4 or the other POU5F1 proteins (Fig. 3b-c).

**Fig. 3.**
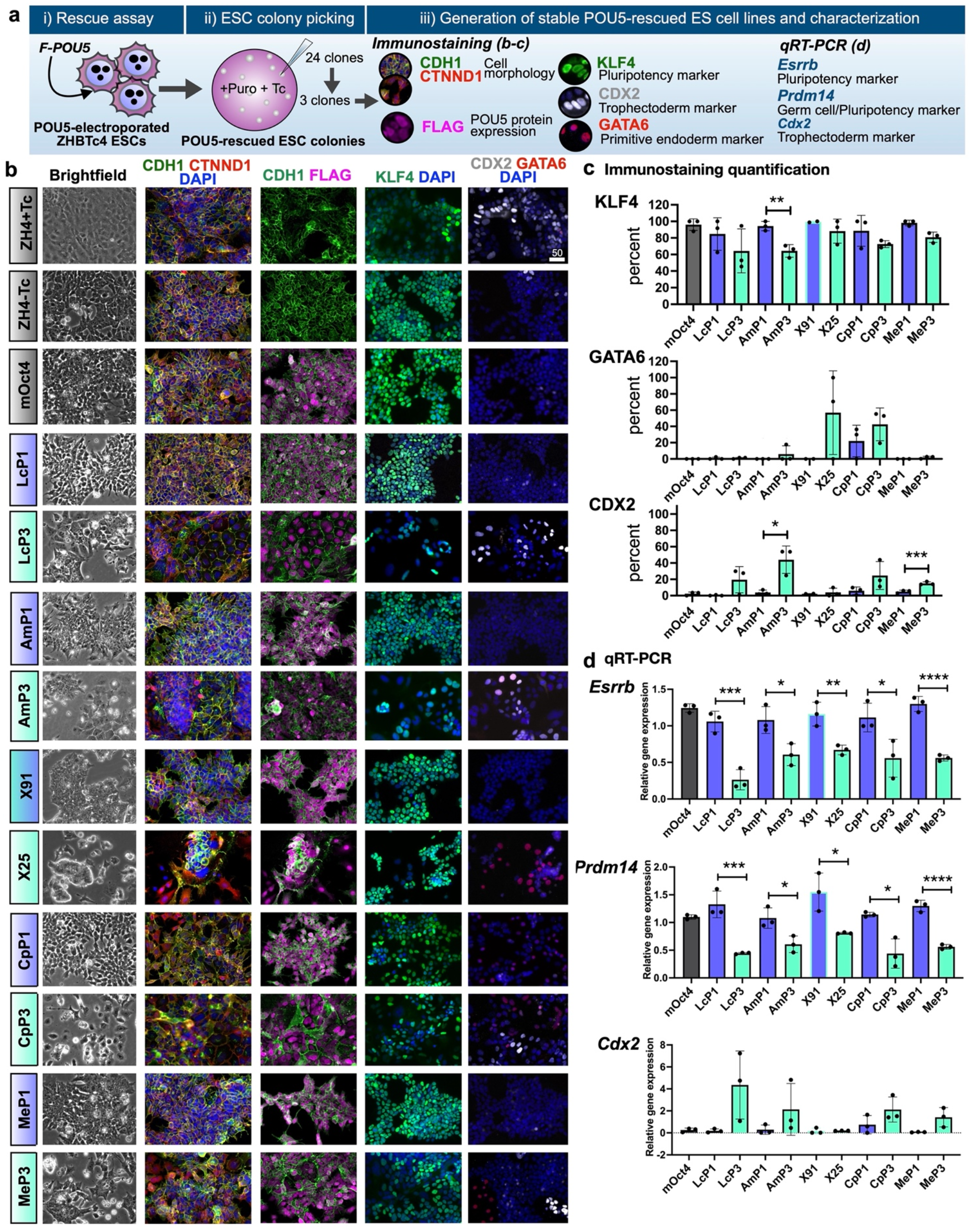
Phenotypes of ESC lines supported by sarcopterygian POU5 proteins. **a**, Experimental strategy used to derive stable ZHBTc4 cell lines rescued by either POU5F1 or POU5F3. **b**, Representative immunofluorescence staining of ZHBTc4-rescued cell lines. Anti-FLAG antibodies were used to detect and localize FLAG-tagged POU5 proteins, anti-KLF4 to assess pluripotency, anti-CDX2 and anti-GATA6 to assess differentiation and anti-CDH1 and anti-CTNND1 to assess cell morphology. **c**, Quantification of biological replicates from immunofluorescence images showing the percentage of KLF4, CDX2 and GATA6 positive cells compared to DAPI (total nuclei). **d**, Relative expression of pluripotency markers (*Esrrb* and *Prdm14*) and differentiation marker (*Cdx2*) in the rescued cell lines quantified by qRT-PCR. The abbreviations for POU5 proteins are the same as in Fig. 2a. Statistical analyses (Unpaired *t*-Test) were performed on n=3 biological replicates (stable clones expanded from the same POU5 rescue experiment).

In agreement with the protein expression data, qRT-PCR showed that POU5F1-rescues expressed high levels of the naïve markers *Esrrb* and *Prdm14* and low levels of the TE marker *Cdx2*, while the reverse generally held true for POU5F3 homologues (with the exception of frog X91) (Fig. 3d). Python POU5F1-rescues expressed *Nanog, Prdm14, Klf4* and *Fgf4* to similar levels as mOct4-rescued cells, suggesting that POU5F1 from species that have lost POU5F3 have similar capacity to support naïve ESC self-renewal (Supplementary Fig. 2f).

### The ability of POU5s to support ESC self-renewal correlates with induction of pluripotency

To test the functionality of the different POU5 homologues in another context, we compared their capacity to support ESC self-renewal with their ability to induce reprogramming. In frog embryos, X60 is expressed maternally and down regulated at gastrulation, both X91 and X25 are expressed in cells about to undergo germ layer induction^38^ and only X91 is expressed in PGCs^46^, correlating with its capacity to rescue OCT4-null ESCs. To explore the ability of these proteins to induce a pluripotent state, as well as monitor reprogramming dynamics, we used a mouse embryonic fibroblast (MEF) line containing a green fluorescent protein expressed from the Nanog locus (Nanog-GFP; Fig. 4a). Reprogramming was performed using a stoichiometric ratio-based infection of equivalent amounts of retroviruses encoding the three factors KLF4, SOX2, c-MYC and a POU5 protein: mOct4, X91 or X25. While both mOct4 and X91 were able to induce Nanog-GFP^+^ colonies, X25 could not. (Fig. 4b, upper panel). However, Nanog-GFP^+^ colonies could be obtained when the dosage of X25 was increased to a 5:1:1:1 ratio with the viruses encoding the other factors (Fig. 4b, lower panel). When compared side by side, X91-iPSCs exhibited less spontaneous differentiation and higher levels of NANOG and SSEA1 (Fig 4c, Supplementary Fig. 3a). Despite the induction of endogenous OCT4 (Fig. 4c), X25-iPSCs exhibited an extensive NANOG negative population (seen in only one X91-iPSC clone), similar to the spontaneous differentiation observed in X25-rescued ESCs (Fig. 3, b and c). Additionally, we observed heterogeneous expression of the pluripotency markers c-KIT and PECAM-1, both within and across different iPSC clones (Fig 4d), with the lowest number of completely reprogrammed cells, both Nanog-GFP^+^ and c-KIT^+^, in X25-iPSCs (Supplementary Fig. 3b). The enhanced capacity of X91 to induce naïve pluripotency was also observed in a higher naïve gene expression signature (Fig. 4e). Similarly, tammar wallaby POU5 proteins (MeP1 and MeP3) could induce AP^+^ iPSCs, although MeP1 was significantly more efficient, correlating with their distinct rescue indices (Supplementary Fig. 3c-d; Fig. 2). Taken together, the difference in reprogramming ability of POU5 proteins validates the functional divergence with regard to pluripotency, as seen in the OCT4-rescue assay.

**Fig. 4.**
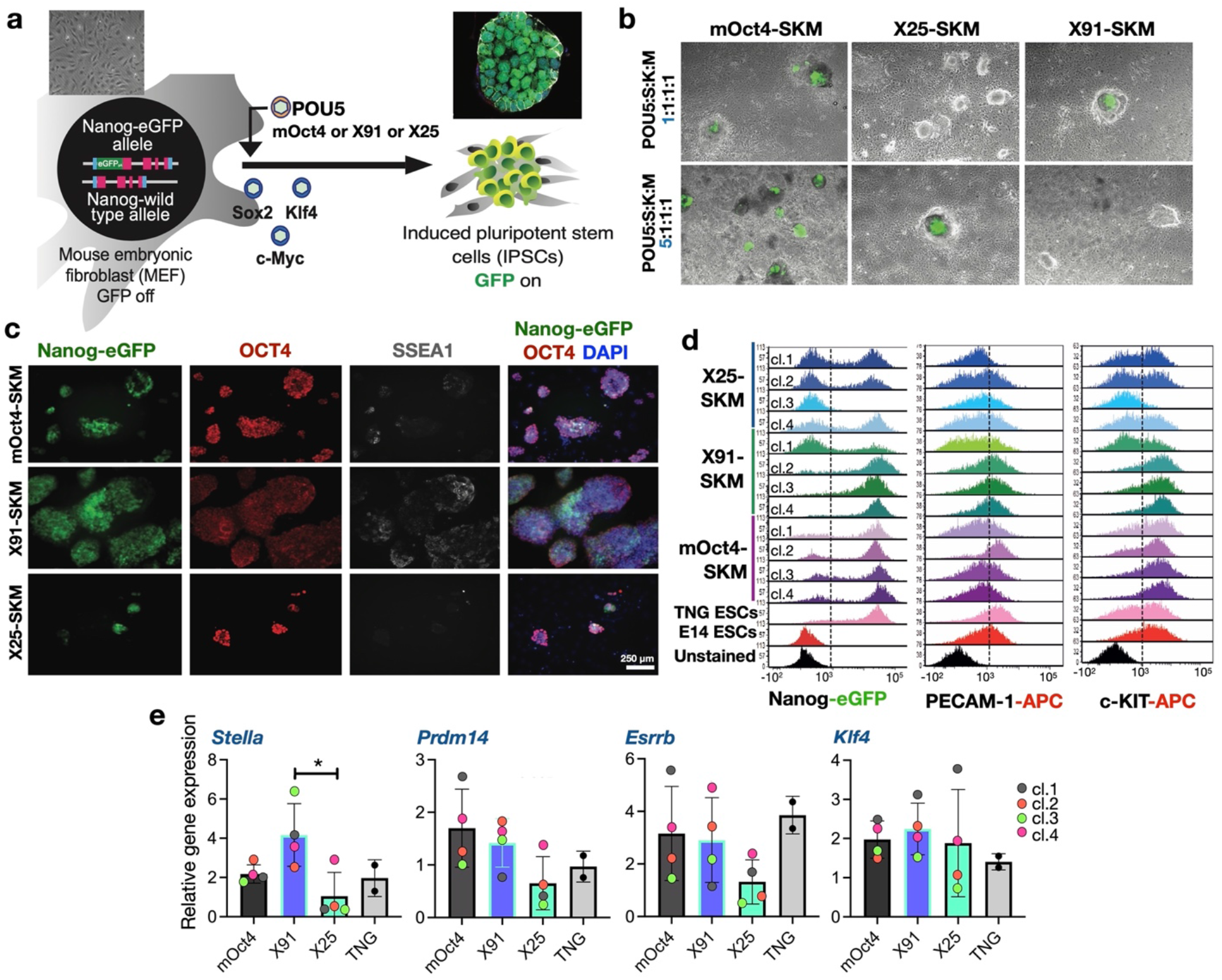
Duplicate POU5 homologues from frog display segregated functions in pluripotency establishment. **a**, Experimental strategy used for murine iPSC generation by retrovirus-based approach. Nanog-GFP mouse fibroblasts were used as a source of somatic cells to monitor the emergence of pluripotency (Nanog positive/green). *Oct4* in OSKM cocktail was replaced with either frog X25 or X91 and either low or high titre of viral infection (for *Oct4* homologues). **b,** Merged images of brightfield and Nanog-GFP represent iPSC colonies at day 24 post-infection. **c**, Characterisation of *Xenopus* POU5s and mOct4-derived iPSC clonal lines by immunofluorescence. Anti-Oct4 and anti-SSEA1 antibodies were used to detect a pluripotency and a germ cell marker, respectively. **d**, Flow Cytometry histograms representing Nanog-GFP, PECAM-1 and c-KIT profiles of mOct4/X91/X25 SKM iPSC clonal lines. **e**, Relative gene expression of germ cells (*Stella* and *Prdm14*) and naïve pluripotency (*Esrrb* and *Klf4*) markers was analysed by qRT-PCR. Data points of each clone (cl.) are also shown. Statistical analyses (Unpaired t-Test) were performed on n=4 biological replicates (from 3 independent experiments).

### Sarcopterygian POU5s exhibit functional segregation of naïve versus primed pluripotency

The functional analyses discussed above suggest that in sarcopterygians retaining both POU5F1 and POU5F3, the former has an enhanced ability to support naïve pluripotency while the latter supports a less stable pluripotent state, giving rise to higher levels of spontaneous differentiation. To characterise this functional divergence and generate a more comprehensive picture of the cell states supported by POU5F1 or POU5F3, we analysed the transcriptome of OCT4-null ESCs rescued by each paralogue. For this analysis, we focused on the coelacanth POU5F1 (LcP1) and POU5F3 (LcP3) forms, which diverged from their tetrapod counterparts around 400 million years ago and exhibit slow rates of evolution (Fig. 1d; Fig. 2a).

Global gene expression analysis on three independent clones for each rescue: LcP1, LcP3 and mOct4, identified 4903 differentially expressed genes (ANOVA with 2-fold change and False Discovery Rate (FDR) ≤ 0.05). We used Morpheus to hierarchically cluster all significantly changing genes based on mean Euclidean distance. The clustering, visualised as a heatmap (Fig. 5a), showed that LcP1-rescued cells were similar to mOct4-rescued cells. Furthermore, naïve pluripotency markers, including germ cell markers, were found in clusters of highly expressed genes in both LcP1- and mOct4-rescued cells while primed pluripotency markers were present in clusters of highly expressed genes in LcP3-rescued cells (Fig. 5a). We also performed pairwise comparisons to explore the subtle differences in global gene expression. This analysis confirmed that the patterns of up-regulated and down-regulated genes of mOct4-rescued cells and LcP1-rescued cells were very similar (Fig. 5b left panel). We next performed GO enrichment analysis on the 605 genes up-regulated in both mOct4 and LcP1-rescued cells compared to LcP3 and identified various naïve state-related categories, e.g. stem cell population maintenance and reproductive process (Fig. 5c top panel). In addition, we found that among the reproduction-related genes up-regulated in both mOct4 and LcP1-rescues, most genes were related to germ cell development, such as spermatogenesis and female gamete generation (Supplementary Fig. 4a). We next identified a set of 1199 genes that were expressed specifically in LcP3-rescued cells and enriched for GO terms including cell junction and tissue development (Fig. 5c lower panel, Supplementary Table 2). Both E-cadherin (*Cdh1*) and N-cadherin (*Cdh2*), as well as other adhesion markers, were significantly up-regulated in POU5F3 as compared to POU5F1-rescued cells (Supplementary Fig. 4b). This link between POU5F3 proteins and positive regulators of adhesion is consistent with what we have previously described^34^ for POU5 protein function as safeguarding epithelial integrity at gastrulation and blocking differentiation as a consequence of epithelial to mesenchymal transition (EMT). Furthermore, among genes common to LcP3- and mOct4-rescued cells, 31 genes were EpiSCs specific (compared to ESCs^47^) and were associated with cell adhesion and extracellular matrix (Supplementary Fig. 4b). In summary, the distinct transcriptomic profiles of ESCs supported by LcP1 and LcP3, suggests alternative roles for these paralogues in naïve versus primed pluripotency, respectively.

**Fig. 5.**
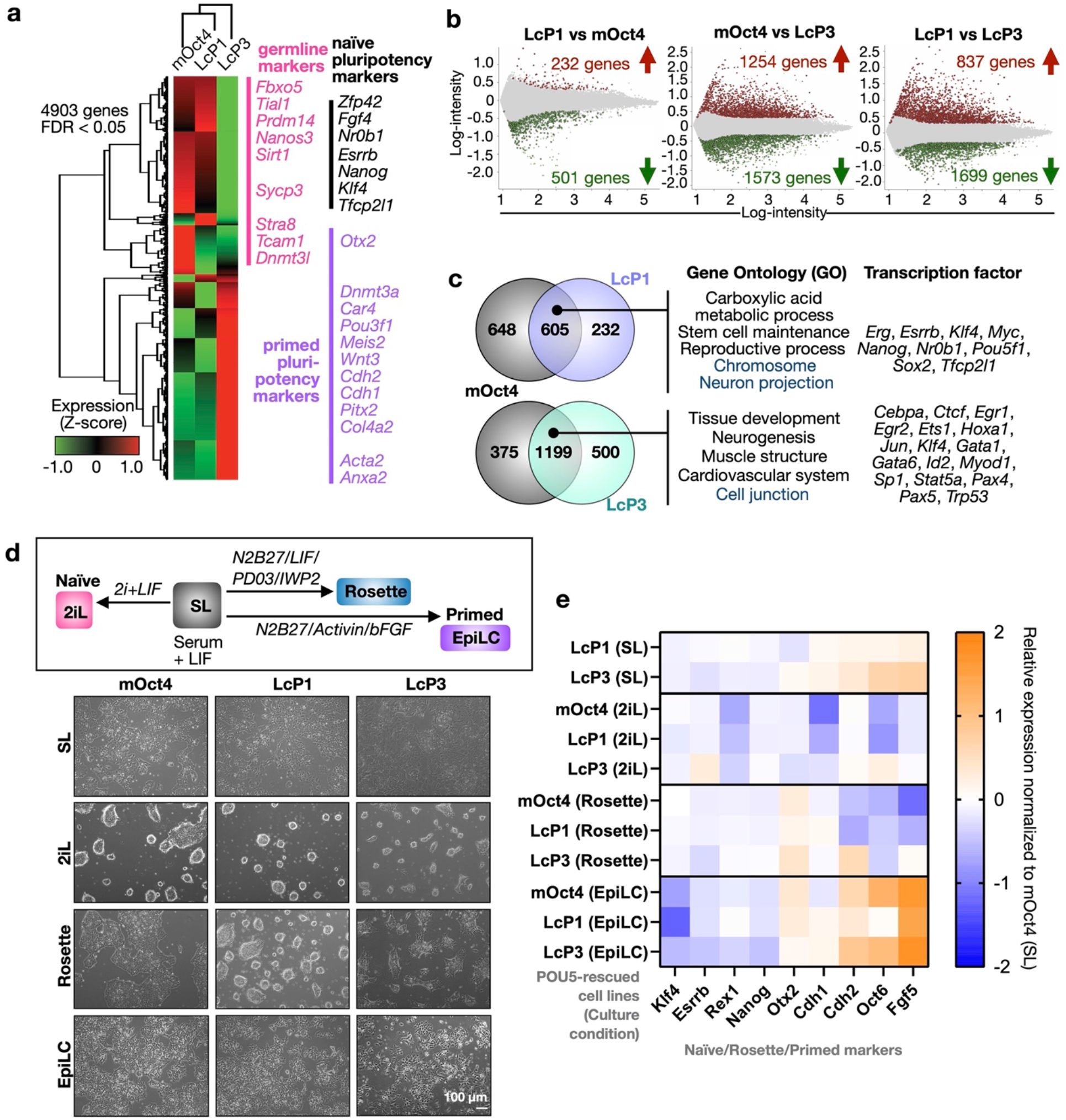
Distinct naïve-primed pluripotency phenotypes are supported by sarcopterygian POU5F1 and POU5F3 proteins. **a**, Heatmap illustrating hierarchical clustering of 4903 differentially expressed genes between the mOct4, LcP1, LcP3-rescued ESC cell lines (fold-change threshold ≥ 2 and FDR ≤ 0.05). The normalised expression level (z-score) of each gene is shown in three-colour format where red, black, green indicate high, medium and low gene expression levels, respectively. **b**, Log-ratio plots showing significantly over-expressed (red) and under-expressed (green) probes, based on the indicated pairwise comparisons (FDR ≤ 0.05). These gene lists were further filtered based on probes corresponding to uniquely annotated genes and with an absolute fold-change cut-off of 2. **c**, Venn diagrams show a common signature in LcP1, LcP3 and mOct4 supported cells. In particular, we find a set of 605 over-expressed genes (a significant majority of both comparisons) and 1199 under-expressed genes common to both LcP1 and mOct4 supported cells, compared to cells supported by LcP3. GO-term analysis (ShinyGO v0.66: Biological Process and Cellular Component, FDR ≤ 0.05) and transcription factor targets (ShinyGO v0.66: TF.Target.RegNetwork, FDR ≤ 0.05) of these gene lists is shown on the right side of the Venn diagram. Full gene list is shown in Supplementary Table 2. **d-e**, Naïve-primed conversion of coelacanth LcP1 and LcP3 rescued ESCs. Cell morphology of the LcP1, LcP3 and mOct4-rescued cells in naïve-primed conversion is shown in bright-field images in panel **d**. The rescued ESC lines, originally cultured in ESC medium (serum + LIF, SL), were driven toward either naïve or primed states: (1) 2i + LIF (2iL) medium, representing the naïve state; (2) Rosette-like stem cells medium, representing the intermediate state between naïve and primed; (3) Epiblast-stem-Cell-Like cells (EpiLC) medium, representing primed state. **e**, Heatmap representing relative gene expression profiles of naïve, rosette and primed pluripotency markers as well as cell adhesion markers. Three clones of LcP1 and LcP3 cell lines into SL/Rosette/EpiLC conditions, alongside with ZHBTc4 control, were analysed by qRT-PCR.

To test the hypothesis that paralogous POU5 proteins have specialized to support either naïve or primed pluripotency, we assessed the ability of both LcP1 and LcP3 to sustain different pluripotent states. Thus, we adapted POU5-rescued cells to either a defined naïve culture with inhibitors of MEK, GSK3 and LIF (2iL), a culture condition that approximates an intermediate pluripotency state, known as rosette-like^13^ or a primed culture Epiblast-like cells (EpiLC)^47^ (Fig. 5d). In line with the transcriptome analysis (Fig. 5a-c), LcP3-rescued cells showed higher levels of primed gene expression in standard serum/LIF (SL) culture (Fig. 5e and Supplementary Fig. 4c). While all rescued cells appeared to eventually adopt a naïve state in 2iL conditions, LcP1 and mOct4-rescued cells adapted faster and showed normal 2iL morphology (Fig. 5d-e). In rosette medium, LcP3-rescued cells showed the highest level of *Otx2*. Finally, when differentiated to EpiLCs, mOct4 and LcP3-rescued cells more effectively up-regulated primed pluripotency markers *Cdh2*, *Oct6* and *Fgf5* (Fig. 5e). Taken together, our data suggest a functional segregation of the sarcopterygian POU5s, with POU5F1 supporting naïve pluripotency and POU5F3 supporting a primed pluripotency gene regulatory network associated with later stages of development, multi-lineage differentiation and gastrulation.

### POU5-mediated mammalian pluripotency first emerged after the gnathostome-cyclostome split

To gain insight into the origin of POU5 pluripotency maintenance activity in vertebrates and into the timing of its functional partitioning between POU5F1 and POU5F3 paralogues in the gnathostome lineage, we analysed the expression pattern of chondrichthyan *Pou5* genes and assessed functionality with the OCT4-rescue assay. We focused on paralogues from one batoid (little skate *Leucoraja erinacea*) and two selachians (whale shark *Rhincodon typus* and small-spotted catshark *Scyliorhinus canicula*). Furthermore, we included the unique POU5 identified in the cyclostome hagfish *Eptatretus burgeri*, which harbours a slowly evolving deduced protein sequence compared to its counterpart in lampreys (Fig. 1d) and is therefore more likely to retain ancestral activities. A simplified phylogenetic tree of the species tested for their POU5 function is depicted in Fig. 6a. First, we analysed the expression of catshark *Pou5f1* (*ScPou5f1)* and *Pou5f3 (ScPou5f3)* from blastocoel formation to neural tube closure (Fig. 6b and Supplementary Information 2). These data show a very similar expression profile for *ScPou5f1* and *ScPou5f3*, with both being broadly expressed in the early embryo, prior to the establishment of the major embryonic lineages (Fig. 6b, i-vi and viii-xiii). At later stages of development, their territories segregate and each paralogue exhibits expression specificities, such as developing PGCs selectively expressing *ScPou5f1* (Fig. 6b, vii) or the anterior hindbrain and tailbud expressing *ScPou5f3* only (Fig. 6b, xiv-xvi).

**Fig. 6.**
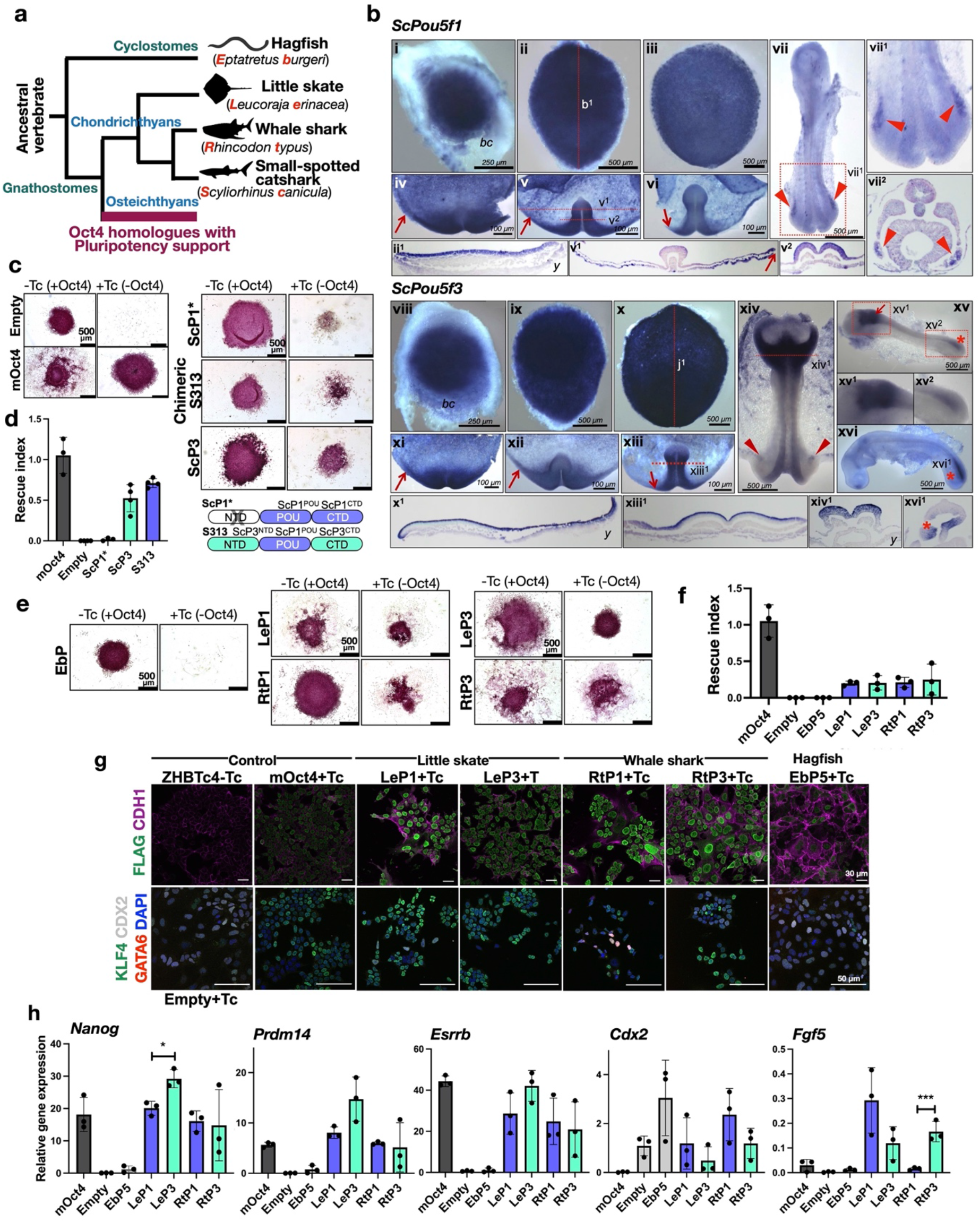
Chondrichthyan but not cyclostome POU5 proteins have the capacity to support pluripotency. **a**, Schematic illustration showing simplified phylogenetic tree of cyclostome and Chondrichthyan species used for testing POU5 protein activities. Abbreviation used in this study are shown in red. **b**, Whole-mount views of catshark embryos following *in situ* hybridisations with probes for *Pou5f1* (*ScP1*) (**i-vii**) or *Pou5f3* (*ScP3*) (**viii-xvi**). Description of each panel is noted in Supplementary Information 3. **c**, AP staining of ZHBTc4 ESC colonies supported by ScP1 and ScP3, cultured in the presence or absence of mOct4 (−/+ Tc). Due to missing N-terminal domain sequence data of ScP1 (ScP1*), a chimeric form of the protein (named S313) was also tested and a cartoon of the swapping construct is shown on the bottom-right. **d**, Rescue indices indicating the capacity of catshark ScP1 and ScP3 to support ESC self-renewal. **e**, AP staining of ZHBTc4 ESC colonies supported by different POU5 proteins including whale shark (Rt), little skate (Le) from chondrichthyans and hagfish (Eb) from cyclostomes. **f**, Rescue indices indicating the capacity of different POU5s to support ESC self-renewal. **g-h**, Phenotypes of rescued ESC lines supported by chondrichthyan POU5 proteins compared to non-rescued cells supported by cyclostome POU5 protein. As Hagfish POU5 (EbP5) cannot rescue, EbP5 expressing colonies were picked and expanded in the absence of Tc, then treated with Tc for four days to remove mOct4 prior to further analysis. **g**, Immunofluorescence staining of POU5 rescued cells using antibodies directed against FLAG-tagged POU5 and CDH1 (E-cadherin) (top panel) and ESC (KLF4)/differentiation (GATA6 and CDX2) markers with DAPI stained nuclei (bottom panel). **h**, Relative expression of naïve pluripotency (*Nanog, Prdm14* and *Esrrb*), primed pluripotency (*Fgf5*) and trophectoderm markers (*Cdx2*) in POU5-rescued cells, quantified by qRT-PCR. Statistical analyses (Unpaired *t*-Test) were performed on n=4 biological replicates (4 independent experiments for rescue index) and n=3 biological replicates (stable clones expanded from the same POU5 rescue experiment).

We then tested the ability of catshark POU5 proteins (ScP1 and ScP3) to support pluripotency using the OCT4-rescue assay (Fig. 2b). Due to a missing N-terminal domain sequence in *ScPou5f1* and based on our finding that the POU domains from frog X91 sufficiently converted the activity of X25 into a POU5F1-like function in the OCT4-rescue assay (Supplementary Fig. 5a-b), we assessed the functionality of ScP1 using a chimeric protein containing the POU domains of ScP1 and the N- and C-terminal domains of ScP3 (named S313) (Fig. 6c). While the chimeric construct was able to support ESC colony formation, differences between the chimeric catshark POU5F1- and POU5F3-supported colonies were hard to distinguish (Fig. 6c-d).

Next, we assessed POU5 homologues from the other chondrichthyans (whale shark *R. typus* and little skate *L. erinacea*) and a cyclostome species (hagfish *E. burgeri*). The number of AP^+^ colonies generated in this OCT4-rescue assay showed that both POU5F1 and POU5F3 proteins from whale shark and little skate (respectively RtP1, RtP3, LeP1 and LeP3) were able to partially support ESC self-renewal in the absence of OCT4, with variable colony morphologies (Fig. 6e). In contrast, hagfish POU5 (EbP5) completely lacked rescue capacity. Unlike sarcopterygians, the average rescue indices obtained with the chondrichthyan paralogues were comparable and generally lower than those obtained with the mOct4 control (Fig. 6f).

To better characterize the functionality of chondrichthyan POU5s, we expanded rescued ESC colonies (cultured in SL +Tc) to generate stable clones and analysed the expression of pluripotency and differentiation markers. As the hagfish POU5 was unable to support any colony formation, clonal lines were generated in the presence of *Oct4* transgene (SL-Tc) and later characterized following subsequent OCT4 removal (Supplementary Fig. 5c). We confirmed that all rescued lines expressed similar levels of both heterologous cDNAs (Supplementary Fig. 5d) and exogenous POU5 proteins (Fig. 6g; Supplementary Fig. 5e). Any variations in the expression of these POU5 proteins did not correlate with their ability to rescue OCT4 activity in ESCs (Supplementary Fig. 5f).

Differences in cellular phenotypes between chondrichthyan POU5F1/3-rescued cells were assessed by immunostaining and qRT-PCR. All rescued lines exhibited a modest level of undifferentiated cells (KLF4^+^) with the exception of EbP5-rescued cells. EbP5-rescues, fixed five days after Tc addition, exhibited similar levels of CDX2 expression as un-rescued control ZHBTc4 cells (Empty) (Fig. 6g). The capacity of chondrichthyan POU5s to rescue pluripotency was confirmed by qRT-PCR, with robust, but variable expression of *Nanog*, *Prdm14* and *Esrrb* (Fig. 6h). Even though chondrichthyan POU5s appeared to support expression of pluripotency genes, they all exhibited low expression of differentiation markers, such as *Cdx2* and *Fgf5* (Fig. 6g-h). Taken together, these data show that all tested chondrichthyan POU5s have some capacity to support mouse ESC self-renewal, with roughly equivalent activities between paralogues, while this capacity is totally absent in the hagfish POU5. This suggests that the determinants underlying specialized POU5 pluripotency-related activities emerged in the gnathostome lineage, after the cyclostome-gnathostome split.

### Structural modelling of POU5 paralogues predicts conserved 3D-elements across vertebrates with the position of specific helices correlating with function in pluripotency

As the POU domains in different homologues, including cyclostomes, have both highly conserved and less conserved regions at the amino acid level (Supplementary Fig. 6), and exhibited variable capacity to rescue OCT4 function, we asked if this difference was reflected at the three-dimensional level of the proteins. For this purpose, we calculated structural predictions for all POU5 homologues using AlphaFold2, an AI system developed by DeepMind to predict three-dimensional protein structures based on their amino acid sequences^48^. AlphaFold2 outputs include measurements of confidence per residue, termed pLDDT, on a scale from 0-100. In all POU5 models, AlphaFold2 predicted the presence of helices in the POU-specific domain (POU-S; α-helices 1-4) and in the POU homeodomain (POU-HD; α-helices 1-3), with folds and most positions being predicted with “very high” confidence (pLDDT>90). In addition, the beginning of the linker between the POU-S and POU-HD was predicted as a helix (Linker α1’), but with variable degrees, from “confident” (90>pLDDT>70) to “low” confidence (70>pLDDT>50). The region between linker α1’ and POU-HD as well as the N- and C-terminal tails were predicted with “low” to “very low” (pLDDT<50) model confidence, suggesting that the latter are unstructured (Supplementary Information 3).

To compare the structures from different species, we asked how the two POU domains, the POU-S (including the linker α1’) and the POU-HD, potentially interacted with DNA. We compared them to an existing crystal structure of mOct4 bound to the *PORE* (Palindromic Oct factor Recognition Element) DNA element (PDB ID: 3L1P, ref^49^). We created POU5-*PORE* DNA structural prediction models for each POU5 homologue by 3D-aligning isolated POU domains with selected conserved amino acid sequences of mOct4 (from PDB 3L1P) and combined them with the 3D coordinates of the *PORE* DNA (from PDB 3L1P). We performed geometry validation and minimization of the resulting POU5-*PORE* DNA models to prevent geometrical clashes (Supplementary Information 3). Before further analysis, isolated structures were verified by Phenix (structural assessment using MolProbity) to ensure a low clash score (cut-off at ten, compared PDB: 3L1P OCT4 on PORE^49^) and to ensure that the analysed residues were Ramachandran favoured (Supplementary Information 3). Finally, we examined how the newly modelled protein-DNA interfaces differed in terms of folds, hydrogen bonding patterns and electrostatic interactions between the POU5 homologue structures and the original mOct4-*PORE* structure.

The super-imposition of all POU5-*PORE* DNA models with mOct4-*PORE* showed similar positioning of all helices, except linker α1’ (Lα1’), with hagfish having the greatest shift in position, suggesting a correlation with its inability to rescue mOct4 activity (Supplementary Fig. 7a). Furthermore, we observed a shift in the orientation of the second helix of the POU-S domain (Sα2) when comparing coelacanth and hagfish proteins (Figure 7a). We then examined the predicted hydrogen bonding (H-bond) interactions between the POU-S-L/POU-HD residues and the *PORE* DNA element for all homologues (Supplementary Fig. 7b). Generally, the predicted protein:DNA H-bonds involved residues located in helices previously reported to interact with DNA^49^ and residues conserved across all species (Supplementary Figures 6 and 7b). Specifically, predicted H-bonds observed in all species, involved residues in the fully conserved third helix of the POU-S domain (Q157^(POU27)^, Q174^(POU44)^, and T175^(POU45)^) and the mostly conserved third helix of the POU-HD (N273^(POU143)^); of note, Q174^(POU44)^ and N273^(POU143)^ have been reported to be essential for iPSCs generation^49^ (Supplementary Figures 6 and 7b). The greatest variation in H-bonds between homologues was predicted for POU-HD residues, showing both species and paralogue-specificity, but not correlating with naïve vs primed POU5 activity.

**Fig. 7.**
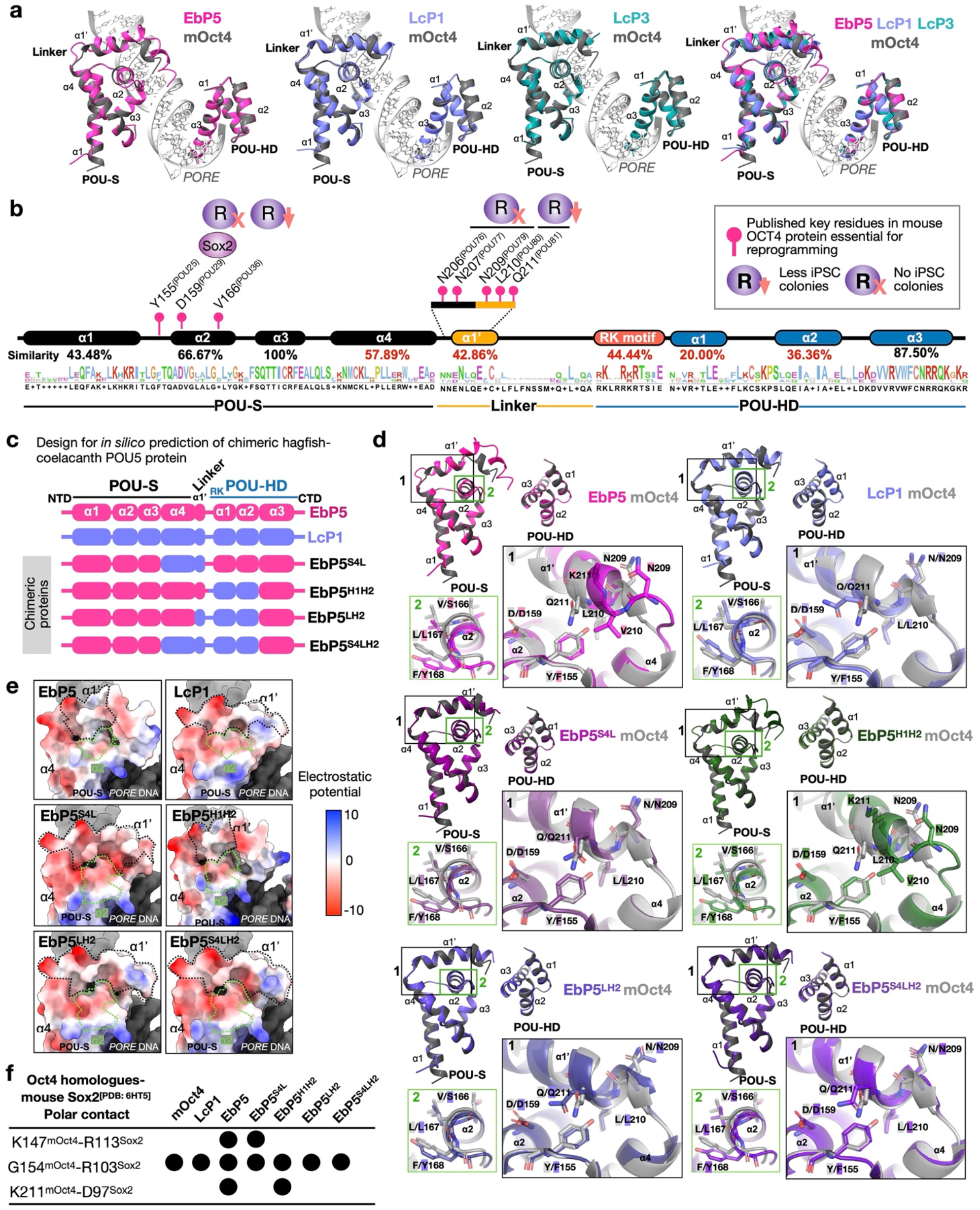
AlphaFold2-based structural models of POU5 homologues predict unique orientations for specific α-helices. **a**, AlphaFold2-based structural prediction models for ordered regions (POU-S-Lα1’ and POU-HD) of coelacanth and hagfish POU5 proteins visualized by ChimeraX with superimposition to mOct4 (grey) on the *PORE* DNA element (PDB ID: 3L1P). **b**, Degree of conservation from the alignment of EbP5, LcP1 and mOct4 protein sequences and key residues of mOct4 (see Supplementary Figure 7 for more details). **c**, Design of EbP5-LcP1 chimeric proteins. We replaced different combinations of un-conserved regions of hagfish EbP5 (pink) with coelacanth LcP1 (lilac). **d**, AlphaFold2-based structural prediction models of chimeric hagfish-coelacanth (Eb-Lc) POU5 proteins, with insets highlighting α2 from POU-S domain (Sα2) and α1’ from Linker (Lα1’). Residues are also marked according to mOct4 numbering in Supplementary Fig. 6. **e,** Predicted electrostatic surface potentials for chimeric proteins with focus on the POU-S-Linker region. Surface charges were determined by ChimeraX, with negatively charged areas shown in red and positively charged in blue. **f,** table summarizing prediction of polar contact interactions between POU5 homologues/chimeric Eb-Lc POU5s and mouse Sox2 using PyMol. Structural models of Eb-Lc chimeric POU5s were generated by AlphaFold2 and the structure of Sox2 was retrieved from 6HT5 (https://www.rcsb.org/structure/6ht5). Number of specific residues indicated in the table are related to mouse Oct4 (as shown in Supplementary Fig. 6) and mouse Sox2. Black dots represent the presence of polar contact interactions.

As the structural changes observed in different POU5 proteins occurred in mOct4 regions identified as essential for reprogramming and support of pluripotency^49–54^ (Figure 7b and Supplementary Fig. 6), we sought to identify the sequences responsible for these structural shifts. For this purpose, we generated a series of *in silico* predictions for chimeras containing elements of the coelacanth POU5F1 and hagfish POU5, focusing on regions less conserved in cyclostomes (Figure 7b, red percentages). The largest swap contained the full region from Sα4 to second helix of the POU-HD (Hα2), with the others containing sections of this region (Fig. 7c). The structures show that only the EbP5^S4LH2^ and EbP5^LH2^ chimeras could re-align Lα1’ and Sα2, which were shifted in the hagfish POU5, as compared to mOct4 (Fig. 7d and Supplementary 7c). In particular, the linker together with Hα1-2 from LcP1 were required to bring Sα2 of EbP5 back in close proximity with Lα1’, making the interaction of key residues (the interface formed by L210^(POU80)^ and Q211^(POU81)^ with Y/F155^(POU25)^) more favourable (Fig. 7d, box 1, Supplementary Fig. 7c). Furthermore, we investigated the electrostatic surface potentials of the POU5-*PORE* structural models, specifically focusing on the solvent-exposed surface areas with low amino acid sequence conservation, Sα2, Sα4, Lα1’ and Hα1-2 (Supplementary Fig. 8). While we observed general differences in surface charge distribution between homologues, the hagfish POU5 solvent-exposed surfaces appeared to be the most neutral. Specifically, the surface charge distribution observed in the region of Sα2 and Lα1’ was rescued by chimeric proteins EbP5^S4LH2^ and EbP5^LH2^, but not by EbP5^S4L^ or EbP5^H1H2^ (Fig. 7e). Similarly, when mOct4-mSox2 polar contacts were predicted (from PDB 6HT5), we found that EbP5 had two additional interactions that were not in mOct4 and were rescued in EbP5^S4LH2^ and EbP5^LH2^ (Fig. 7f). Taken together, our *in silico* modelling suggests that the region including the linker and the first two helices of the homeodomain play a key role in orienting the structure, resulting in specific helix-helix and protein-protein interactions.

To test whether the re-orientation of Lα1’ and Sα2 was sufficient to support pluripotency *in vitro*, we engineered two chimeras, EbP5^S4LH2^ and EbP5^LH2^, and evaluated their functionality using the OCT4-rescue assay (Fig. 2b; Fig. 8a). Both chimeras supported formation of undifferentiated colony, but showed differences in their proliferative ability, as seen by the reduced size of the EbP5^LH2^-rescued colonies (Fig. 8b, c). To understand the phenotypic differences between EbP5^S4LH2^ and EbP5^LH2^-rescued cells, we established clonal cell lines with stable chimeric protein expression (Fig. 8d) and compared their gene expression profiles by qRT-PCR (Fig. 8e). Both chimeras supported the expression of key pluripotency markers, such as *Nanog*, *Prdm14*, *Esrrb* and *Fgf4* and efficiently suppressed *Cdx2* expression, similarly to mOct4 and LcP1.

**Fig. 8.**
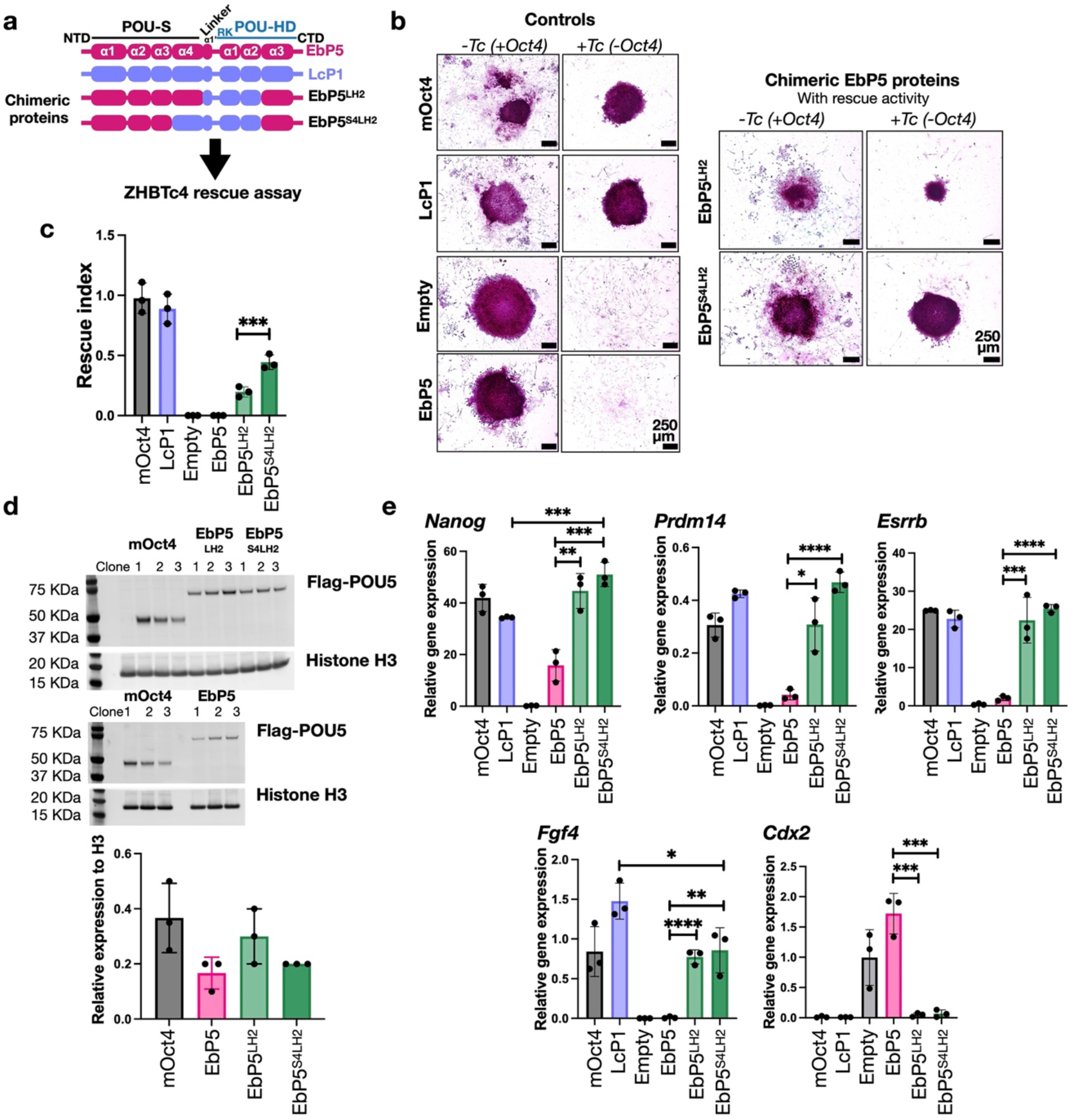
Replacing specific regions of hagfish POU5 with their coelacanth POU5F1 counterparts is sufficient to rescue pluripotency in OCT4-null mouse ESCs. **a**, Design of the EbP5-LcP1 chimeras used to test rescue capacity in OCT4-null mouse ESCs **b**, Colony phenotypes of ZHBTc4 cells transfected with different chimeric POU5 proteins grown in the presence or absence of mouse Oct4 (−/+ Tc) and stained for alkaline phosphatase (AP) activity (purple). **c**, Rescue index of OCT4-null ESCs rescued by chimeric hagfish POU5 proteins. **d**, Western blot showing protein expression of 3xflag-tagged chimeric Eb-Lc POU5 proteins from three rescued clones per species, with quantification below. **e**, Relative expression of naïve pluripotency (*Nanog, Prdm14, Esrrb* and *Fgf4*) and trophectoderm markers (*Cdx2*) in chimeric hagfish POU5-rescued ESC clonal cell lines, quantified by qRT-PCR. Statistical analyses (Unpaired *t*-Test) were performed on n=3 biological replicates (stable clones expanded from the same POU5 rescue experiment).

In conclusion, with a combination of sequence alignments, structural modelling and domain swapping, we pinpointed the region of gnathostome POU5F1 that is sufficient to inhibit differentiation and support naïve ESC self-renewal in the absence of mOct4. This suggests that the 3D position of the linker and the regions flanking it may have been key evolutionary targets for the establishment of an Oct4-centric gene regulatory network associated with pluripotency.

## Discussion

Here we show that since their emergence in vertebrates, POU5 proteins have undergone a complex step-wise evolution, enabling the eventual emergence of the naïve and primed pluripotency states of mammals. This evolutionary history involves the segregation and integration of multiple spatial and temporal inputs into a core network safe-guarding cell potency, which can be traced back to the origin of gnathostomes (Fig. 9).

**Fig. 9.**
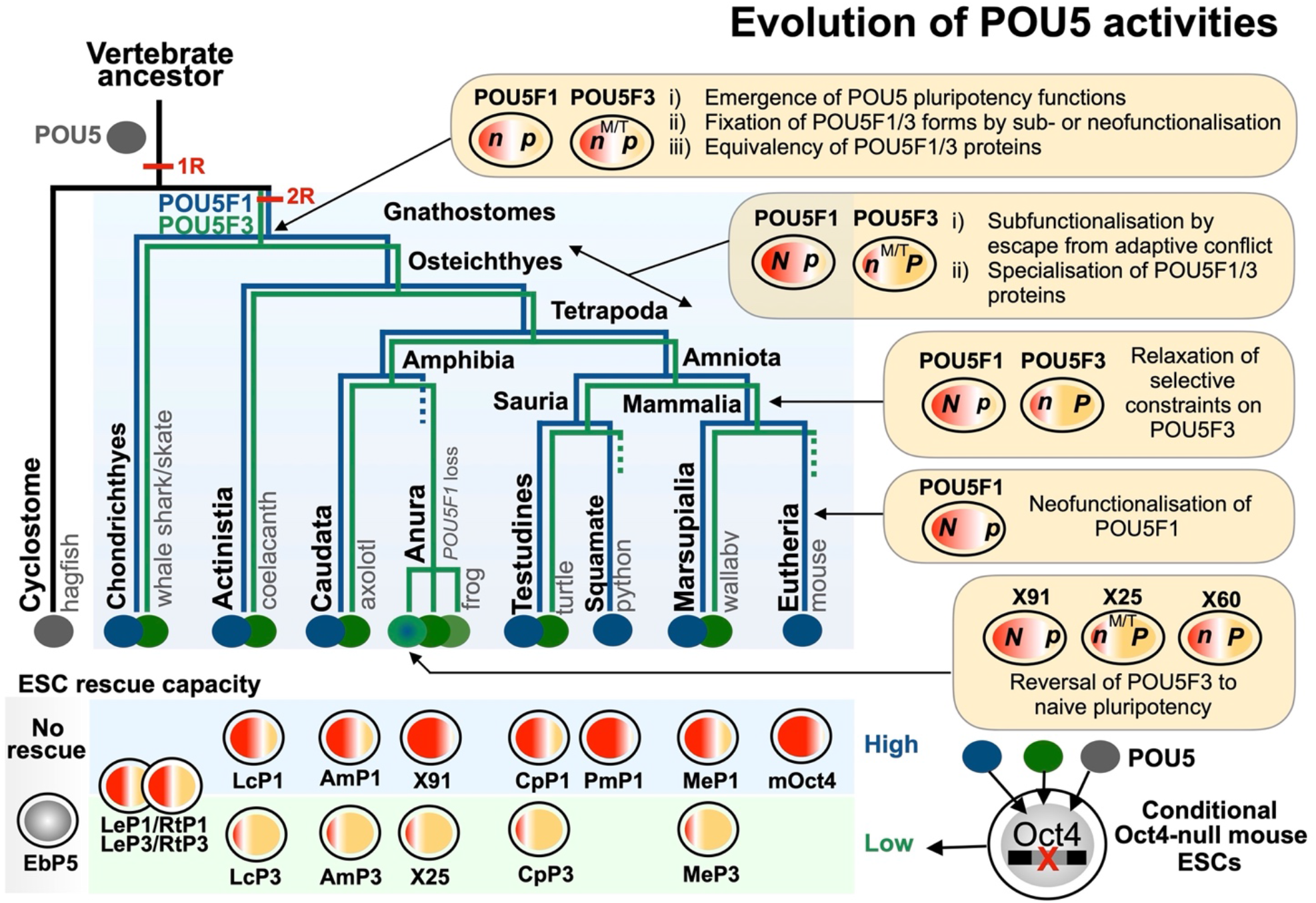
Summary of the evolution of POU5 activities in vertebrates. A simplified phylogenetic tree summarises the evolution of the *Pou5* gene family in vertebrates as inferred from genomic searches and sequence analysis. Most salient events were (1) the emergence of the family in the vertebrate lineage (black branch), (2) the maintenance of one copy in cyclostomes (grey branch) and (3) a duplication giving rise to the two gnathostome *Pou5f1* and *Pou5f3* paralogues (blue and green branches respectively), which may have been part of the 1R/2R whole genome duplications that took place in vertebrates. Dotted lines indicate lineages in which one paralogue was lost. Right panels point at key nodes of the tree and indicate major milestones in the functional evolution of the gene family, as inferred from expression and functional analyses of POU5F1 and POU5F3 proteins from selected species (box at the bottom of the figure shows a summary of the activities observed in the Oct4 rescue assay). These include (1) the emergence of the capacity of POU5 proteins to support pluripotency, which predated the duplication generating the *Pou5f1* and *Pou5f3* genes, but is not shared by cyclostome POU5 proteins, (2) the preservation of both gnathostome paralogues possibly related to expression specificities, fixed for each form prior to the gnathostome radiation, (3) functional specialisations of paralogous proteins that took place early in the sarcopterygian lineage and could have paved the way to elaborations of naïve and primed pluripotency states of eutherians. Additional evolutionary changes, including reversals or innovations, have paralleled losses of one paralogue in anurans and in eutherians (*Pou5f1* and *Pou5f3* respectively). N/n and P/p refers to the capacity of POU5 paralogous proteins to support naïve and primed pluripotency in OCT4 rescue assays (“N” and “P” refer to a strong activity, “n” and “p” to a low one). M/T refers to specific expression traits of *Pou5f3* at the neural tube, anterior hindbrain and tailbud, conserved across gnathostomes, including chondrichthyans, which may have contributed to the preservation of this paralogue following the *Pou5f1/Pou5f3* gene duplication.

Pinpointing the timing of gene losses and duplications is an essential stepping stone in understanding functional evolution of a multigene family. The identification of *Pou5f1* and *Pou5f3* orthologues in chondrichthyans and a single gene in cyclostomes, without a clear relationship to a specific class, indicates that the gene family emerged in the vertebrate lineage prior to the split between gnathostomes and cyclostomes, and that the *Pou5f1*/*Pou5f3* duplication predated gnathostome radiation (Fig. 9). Analysis of key amino acid positions supports the monophyly of the group containing the lamprey and hagfish *Pou5* genes, in line with the phylogenetic position of these species and of the group containing both gnathostome paralogues^55^. The detection of extended conserved syntenies between gnathostome *Pou5f1* and *Pou5f3* loci suggests that the duplication generating these two forms was a part of the two rounds of whole genome duplications (WGDs) that took place in early vertebrate evolution^56,57^. The timing of these two rounds was recently clarified, with the first round occurring prior to the gnathostome-cyclostome split, and the second one occurring in the gnathostome lineage, prior to its radiation^58^. The detection of extended syntenies between the gnathostome *Pou5f1* and *Pou5f3* loci indicates that these classes were generated by a large-scale, possibly genome wide, duplication event and the monophyly of gnathostome *Pou5* supported by our sequence analyses further suggests that this duplication may have been part of the second round of vertebrate WGD (Supplementary Fig. 9a-c).

A key finding of our OCT4-rescue assay is that the hagfish POU5 protein is unable to support pluripotency, while all osteichthyan POU5F1 or POU5F3 proteins tested exhibit some capacity to do so. This also holds true for chondrichthyan POU5 proteins, although the degree to which they support pluripotent phenotypes is variable. Altogether, this indicates that the origin of the structural determinants, which underlie the regulation of the OCT4-centric pluripotency network in ESCs, can be traced back to the origin of gnathostomes, prior to the duplication, which generated the two classes. The broad expression of both paralogues in chondrichthyans combined with their similar modest functional activity is consistent with potential roles in germ cell and gastrulation stage pluripotency, suggesting these properties were fixed early in the gnathostome lineage.

Our systematic search for *Pou5* genes in a broad sampling of gnathostomes shows that despite losses of either paralogue (ref^24^; this study), both *Pou5f1* and *Pou5f3* were retained at the base of the chondrichthyan, actinopterygian and sarcopterygian lineages, as well as in the last common ancestor of actinistians, amphibians, sauropsids and mammals (Fig. 9). This suggests that distinct selective forces acted to preserve both paralogues shortly after duplication, in agreement with evolutionary models for maintenance of duplicate genes^59^. In line with an early specialization of each form in gnathostomes, chondrichthyan POU5F1 and POU5F3 display unique expression characteristics, selectively maintained in osteichthyans. For instance, in the catshark, the anterior hindbrain expresses *Pou5f3*, similar to chick, frog and zebrafish^36,38,60,61^ while the developing yolk sac endoderm exhibits a *Pou5f1* expression reminiscent of *Oct4* in the primitive endoderm of mammals^30,62^. These territories may reflect ancient class-specific expression features, fixed prior to gnathostome radiation, which contributed to the early preservation of both paralogues, either by neo-functionalisation, or duplication-degeneration-complementation. A dosage selection effect may also have been involved, consistent with the expression of the two catshark paralogues and with the dose sensitivity of OCT4 in ESCs^27^. This initial expression diversification of the two classes at the gene level may have paved the way to subsequent specializations at the protein level, further contributing to their maintenance. In agreement, our analysis shows that sarcopterygian POU5F1 orthologues, from species harbouring both paralogues, were significantly more able to support naïve pluripotency, while POU5F3s showed a higher capacity to support primed pluripotency, unlike chondrichthyan paralogues. These findings suggest that the dual functionality observed for mOct4, has an alternative resolution in sarcopterygians that retain both genes, through the segregation of paralogues into either naive or primed pluripotency functions. Specialisation of duplicates is consistent with an evolutionary mode of escape from adaptive conflict, whereby the duplication of an ancestral bi-functional gene results in the specialisation of each paralogue, optimising its capacity to fulfil one function, while impeding its capacity to perform the other^59^. We propose that this process led to a functional diversification of POU5F1 and POU5F3 proteins early in the osteichthyan lineage, such that POU5F1 orchestrated the preservation of the germ line, insulating it from extrinsic differentiation signals, while POU5F3 specialised to manage gastrulation specific signals, through the regulation of adhesion, migration and differentiation.

Even though the functional specialisation of POU5F1 and POU5F3 appeared to be maintained in a number of sarcopterygian taxa, our data highlight multiple losses of either one of the two paralogues (Fig. 9). How can organisms cope with the loss of genes harbouring specialised functions? Perhaps evolutionary innovation focused on the region that was responsible for the emergence of the POU5-centric pluripotency network. Supporting this idea, we found a coding sequence in LcP1 that influences key structural elements in POU5 proteins and conveys POU5 activity to the hagfish protein, not only endowing this protein with chondrichthyan-like POU5 activity, but with POU5F1-like capacity to support naïve pluripotency. Central to these structural elements are a number of residues that are crucial for the support or induction of pluripotency by mOct4 (Fig. 7b and Supplementary Fig. 6), such as the POU-S domain residues D159^(POU29)^ required for the mOct4-mSox2 interaction and iPSC formation^53^ and V166^(POU36)^ required for optimal reprogramming^49^. In addition, multiple positions in the first helix of the linker region have been identified as important for reprogramming^49^, including positions N206^(POU76)^, N207^(POU77)^, N209^(POU79)^, L210^(POU80)^ and Q211^(POU81)^. Simultaneous mutation of N206^(POU76)^, N207^(POU77)^, N209^(POU79)^ and L210^(POU80)^ abolishes OCT4-rescue activity^49^. However, all of these amino acids are ultimately conserved in both POU5F1 and POU5F3, and as result their identity does not explain the differences in naïve versus primed pluripotency observed here. Therefore, we looked for residues that were unique to the specific paralogues. Although we identified variations within the linker, no obvious naïve motif was apparent. While position D205^(POU75)^ in mOct4 is conserved in LcP1, but is an E in LcP3, the RK motif in LcP3 contains an extra R and there is homeodomain position, L250^(POU120)^, that is conserved in mOct4 and LcP1, but is a S in LcP3. However, these specific differences are not found in X91, have not been identified via mutational screens, and have no clear assigned function. Therefore, it is not our contention that these residues give POU5F1 its capacity to support naïve pluripotency. Instead, we favour an explanation that involves the coevolution of multiple changes that preserve the structural integrity of protein-protein interaction surfaces, including the influence of positions in the homeodomain on the structure of the linker and the POU-S domain. In *Xenopus*, where loss of *Pou5f1* was followed by gene duplication, one of the three POU5F3 proteins evolved the ability to support a naïve-like pluripotency. Sequence comparisons highlighted a rapid rate of evolution and extensive divergence of the POU domain relative to other POU5F3 proteins, suggesting multiple compensatory interactions that could re-orient the two key structural motifs discussed here.

Upon the loss of a paralogue, perhaps higher concentrations of the remaining POU5 could compensate for the loss of the other form and any co-evolved binding partner specificity; potentially also resulting diversifications of developmental strategies. For instance, the timing and mechanism whereby PGCs segregate from somatic cells extensively vary across metazoans, and the study of model organisms has highlighted two radically different modes: pre-formation and epigenesis. The first relying on an early specification by maternal determinants, while the second depends on a later induction from surrounding tissues^63^. Intriguingly, all osteichthyans that have lost *Pou5f1* employ pre-determination, a derived trait in vertebrates (chick, *Xenopus*, sturgeon, zebrafish^63,64^), while closely related species that have retained this paralogue use induction, such as the axolotl in amphibians, or the turtle in amniotes^65,66^. This correlation suggests that an epigenesis strategy for PGC specification was a driving force to preserve *Pou5f1* in osteichthyans, in line with the specialisation of the protein into naïve pluripotency. This selective constraint was relaxed upon the transition to a pre-formation mode, involving an early determination of the germ line. Supporting this hypothesis, a remarkably high evolutionary rate of POU5F1 is observed in crocodilians, while the gene is lost in the bird lineage. The biological significance of the *Pou5f3* losses observed in eutherians and squamates is less clear. While mouse and human OCT4 have robust capacity to support primed and naïve pluripotency, we have also found a similar naïve POU5F1 activity in snakes. The specific expression of the *Pou5f3* class in the anterior hindbrain and tailbud, as well as their support of primed pluripotency, suggest that related developmental processes may have diversified. All sarcopterygian POU5F1 proteins tested were endowed with the capacity to repress spontaneous trophoblast differentiation in ESC cultures. This property was not shared by chondrichthyan POU5 proteins, but is clearly encoded in the region spanning the POU-S-L and POU-HD domains encoded in the earliest sarcopterygian POU5F1 (coelacanth) tested and can be transferred to heterologous proteins. These data suggest an early emergence of the corresponding structural determinants of POU5 proteins in gnathostomes, followed by an elaboration phase taking place selectively in the POU5F1 lineage, after the gnathostome radiation. In line with this hypothesis, repression of a *Cdx* family member by POU5 proteins has been reported in *Xenopus*^38^, and *Cdx2, Pou5f1* and *Pou5f3* expression at the level of elongating posterior arms in the catshark are consistent with an ancient origin of this regulatory node (this study; ref^48^). A key innovation of mammals may have been its co-option into the developmental context of the blastocyst, regulating the trophoblast lineage commitment, as observed in the mouse^30,32^.

Pluripotency is a specific functional definition that was initially coined to describe the capacity of mammalian cells to differentiate in response to experimental manipulation and evolved to become a developmental concept describing the state or the potential of early embryonic progenitors, as compared to immortal cell lines derived from early mammalian embryos. While underlying gene regulatory networks, or more specifically, pluripotency networks, have been extensively analysed in eutherian mammals, attempts to extend this notion to species outside mammals have been plagued by ambiguous sequence comparisons or non-conservation of functional activities. Despite the fundamental importance of preserving potency in early development, the extent to which key regulators of the pluripotency network have shifted during evolution has been surprising. By exploring the functional evolution of one of the fundamental regulators in the pluripotency network, we have traced the origins of an OCT4-centric network to the emergence of gnathostomes and showed that its evolution is intimately linked to the strategy used to preserve the germ line from extrinsic differentiation signals. Our work sheds light on the evolutionary forces, which drive the extensive diversification of pluripotency networks across gnathostomes, including developmental contexts, the mode of germ line specification and variations in early embryonic architecture. In conclusion, we present a highly nuanced story describing the evolution of POU5 family and suggest that phenotypic studies restricted to a single model organism can only provide a snapshot of the pluripotency network linked to this pivotal component.

## Methods

### Plasmid construction

Expression plasmids carrying *Pou5* coding sequence (CDS) were generated for ZHBTc4 ESC rescue experiment by inserting the triple flag-tagged (3xflag) *Pou5* coding sequences into pCAGIP vector^43,67^ between the CAG promoter and the *IRES-PAC* (Puromycin resistant gene encoding puromycin N-acetyl-transferase). The sources of *Pou5* genes used for the rescue assay are listed in Supplementary Table 1. *Pou5* CDS for *CpP1*, *CpP3*, *EbP5*, *LcP1*, *LcP3*, *LeP1*, *LeP3*, *MeP1*, *MeP3*, *RtP1*, *RtP3*, *ScP3* and chimeric constructs *S313*, *EbP5^LH2^* and *EbP5^S4LH2^* were synthesised by gBlock (IDT) and Gene synthesis (Invitrogen) services. *Xho*I/*Not*I sites were used to insert *Pou5* fragments into the pCAG 3xflag mOct4 vector in replace of the mouse *Oct4* CDS. For *LcP1*, *AmP1*, *AmP3* with *Xho*I sites present in the CDS, GeneArt® Seamless Cloning & Assembly (Invitrogen) was used to subclone the *Pou5* CDS into pUCL19 carrying a 3xflag sequence. The *3xflag Pou5* CDS were then inserted by transfer a XbaI/Not1 fragment into the same sites in the pCAG vector. DNA sequencing was performed by GATC Biotech.

### Mouse ESC culture

Mouse ESCs were routinely cultured as described by ref^38^. Briefly, complete mouse ESC medium was composed of Glasgow Minimum Essential Medium (GMEM) containing 0.1 mM non-essential amino acids, 2 mM L-glutamine, 1.0 mM sodium pyruvate, 0.1 mM β – mercaptoethanol, 10% Fetal Bovine Serum (FBS) and murine LIF (homemade). The flasks/dishes (Corning) for ESC culture were coated with 0.1% gelatin in PBS. 2iL, Rosette and EpiLC culture conditions were described in Supplementary Table 3.

### ZHBTc4 ESC rescue experiment

pCAGIP-POU5 expression vectors were linearised with *Sca*I or *Pvu*I. ZHBTc4 ESCs (1X10^7^) were electroporated with 100 μg of linearised pCAG-IP-POU5 plasmid (Gene Pulser Xcell™ Electroporation Systems at 0.8 kV, 10 μF, 0.4 mm cuvette). Electroporated cells (1X10^6^) were then plated onto gelatinised 100 mm culture dishes containing ESC medium with and without tetracycline (Tc, 2 μg/mL). At day 2 post electroporation, the medium was replaced with ESC medium supplemented with 1 μg/mL puromycin (with and without Tc) to select the cells expressing transfected *POU5* genes and the medium was changed every other day thereafter. At day 9 post electroporation, several ESC colonies were big enough to be picked for expansion and used to generate stable ESC lines from both plus and minus Oct4 conditions (without and with Tc). The ESC colonies were also fixed and stained for alkaline phosphatase activity. To better elucidate the phenotypes of stable POU5 rescued lines, three clonal cell lines were characterised at passage 6, for each POU5-rescue experiment.

### IPSC generation

To produce retrovirus particle for infecting Nanog-GFP MEF cells, packaging cell lines Plat-E were transiently transfected using Lipofectamine LTX (Invitrogen) with two expression vectors: pMXs-vector carrying gene of interest and pCL-ECO containing modified gene encoding retroviral components. Retrovirus supernatant or medium containing virus particles was harvested at day 2 post transfection and concentrated by Retro-Concentrator (Clontech) solution. The titer of retrovirus was measure by Retro-X qRT-PCR Titration Kit (Clontech). For iPSC generation, transgenic mouse embryos at embryonic stage 13.5 were collected for MEF derivation. The embryos were from the cross of male Nanog-eGFP mice (Ian Chambers, University of Edinburgh) with female 129S2/ScPasCrl (Charles Reiver) Nanog-eGFP/129. For ethical approval, the mice were maintained, bred, and manipulated at University of Copenhagen, SUND transgenic core facility (animal work was authorized by project licenses 2012-15-2934-00142, 2013-15-2934-00935 and 2018-15-0201-01520, Danish National Animal Experiments Inspectorate (Dyreforsøgstilsynet)). For iPSC induction, Nanog-GFP MEF cells were infected with ectopic retrovirus carrying Oct4 or POU5 homologue gene (X25 or X91) together with other retrovirus carrying Sox2, Klf4 and c-Myc. The infection was done at day 0 and day 1 under MEF medium. On day 3, MEF medium was replaced with defined iPSC induction medium. On day 4, induced cells were seeded onto irradiated feeders. Medium was changed daily from day 6 to day 10 and every two-day from day 12 onward. Infected cells and iPSCs were cultured on the irradiated feeders and in defined iPSC induction medium composed of DMEM high glucose (ThermoFisher), 20% KnockOut Serum Replacement (ThermoFisher), 0.1 mM non-essential amino acids (Sigma), 2 mM L-glutamine (ThermoFisher), 0.1 mM β –mercaptoethanol (Sigma), LIF (homemade), 20 µg/mL Vitamin C (L-ascorbic acid, Sigma), 0.5 µM Alk5 inhibitor (A83-01, Tocris).

### Alkaline phosphatase (AP) staining

The Leukocyte Alkaline phosphatase kit was used for AP staining according to the manufacturer’s instructions. Briefly, cells were fixed with a fresh mixture of 25 mL citrate solution, 8 mL 37% formaldehyde and 65 mL acetone. The fixed cells were then washed twice with tap water and stained with fresh AP solution, which was generated by mixing 400 μL of FRV alkaline phosphatase solution and 400 μL of sodium nitrate solution and incubating the mixture in the dark for 2 minutes. Subsequently, the mixture was added to 18 mL of water and mixed well, followed by addition of 400 μL of naphthol. A volume of 5 mL of this mixture was immediately added to the fixed cells, followed by a 25 minutes incubation in the dark at room temperature. The stained cells were washed twice with tap water and air dried overnight. Images of AP colonies were acquired using a Leica-5500B microscope and then processed using Fiji ImageJ image processing software. The stained colonies were categorised into 3 classes, undifferentiated, mixed and differentiated, based on the intensity of AP staining. The rescue index was calculated by dividing (1) the number of rescued AP positive ESC colonies obtained in the absence of endogenous Oct4 with (2) the number of colonies obtained in the presence of endogenous Oct4 for a given transfection.

### Immunofluorescence

POU5-rescued ESCs at passage 6 were seeded onto 8-well 15μ-Slide (Ibidi) at a density 20,000 cells/well. The cells were grown for two days and then fixed with 4% paraformaldehyde (PFA). The list of antibodies and details of their application is provided in Supplementary Table 3. Primary antibodies were diluted in blocking solution (containing TritonX 100, serum and BSA) and used to stain cells overnight at 4° C. Cells were then stained with secondary antibodies diluted 1:800 in blocking solution for 1 hour at room temperature in the dark. Cells were washed three times with PBS after each antibody incubation. Samples were imaged on a Leica AP6000 microscope and within each experiment, all images were acquired using identical acquisition settings and analysed by Fiji^68^. E-cadherin (CDH1) and p120 catenin (CTNND1) was chosen as membrane-associated marker to observe cell morphology. KLF4, CDX2 and GATA6 are markers for undifferentiated naïve ESCs, trophectodermal lineage and PrE, respectively. Immunofluorescence quantification was performed using CellProfiler^69^. Briefly, fluorescent images for KLF4, GATA6, CDX2 or DAPI staining of POU5-rescued cells were uploaded and run on CellProfiler software using a revised pipeline (Supplementary Information 2). The output showing the number of accepted objects indicates the number of cells with specific signals. Number of KLF4, GATA6 or CDX2 positive cells against DAPI positive cells (total cells in fluorescent image) were calculated as a percentage to compare between different POU5-rescued lines. Data points in the bar charts are the percentage of each biological clone.

### Western Blots

Cells were washed once with PBS and then lysed directly on the plate by addition of 2x Laemmli buffer (4% (w/v) SDS, 20% (v/v) glycerol, 120 mM Tris-HCl pH 7.4). Samples were heated for 5 min at 70° C, sonicated for 10 secs at 40% power using a Sonopuls mini20 (Bandelin) and centrifuged for 10 mins at 14,000 x g to clear the lysates. Protein concentration was determined using NanoDrop 2000 (Thermo Scientific). A sample volume of 20 µl containing 40 µg of protein, supplemented with 2 µl of 1 M DTT and 1 µl of bromophenol blue, was loaded per lane on NuPAGE 4–12% Bis-Tris Protein Gels (Invitrogen). Electrophoresis was performed in 1x NuPAGE MES SDS running buffer (Invitrogen) at 190 V for 45 min. Proteins were transferred to Nitrocellulose blotting membranes (GE Healthcare) at 400 mA for 70 min on ice in cold transfer buffer (25 mM Tris base, 190 mM Glycine, 20% Methanol). After washing in TBST (20 mM Tris (pH 7.5), 150 mM NaCl, 0.1% Tween 20), membranes were blocked for >1 hr at RT in TBST containing 10% Skim milk powder. All primary antibody incubations (overnight at 4 °C) were performed in TBST containing 5% BSA, followed by three washes in TBST and secondary antibody incubations (2hrs at RT) were performed in TBST containing 5% Skim milk powder. Blots were imaged on a Chemidoc MP (Biorad), and then quantified using ImageJ.

### Quantitative RT-PCR (qRT-PCR)

RNA and cDNA preparations were performed using the RNeasy^TM^ Mini Kit and SuperScript® III Reverse Transcriptase, respectively, according to manufacturer’s instructions. Quantitative RT-PCR was performed using the Roche Universal ProbeLibrary (UPL) System and UPL primers were designed using the Roche Assay Design Centre. All UPL primers and probes used in this study are listed in Supplementary Table 3. PCR reactions were performed using the LightCycler® 480 Probes Master Mix. Briefly, a 10 µl reaction of UPL qRT-PCR was composed of 5 μL of Probes Master Mix, 0.45 μL of 10 μM forward/left primer, 0.45 μL of 10 μM reverse/right primer, 0.1 μL of specific probe, 2 μL of diluted first strand cDNA, and 2 μL of RNase-free water. Concentration value for each gene of interest were normalised to that of the housekeeping gene *Tbp* to obtain the relative transcript level.

### Microarray Processing and Analysis

Global gene expression profiles of POU5-rescued ESC lines were obtained using Agilent one-colour microarray-based gene expression analysis according to the manufacturer’s instructions. High quality total RNA (RNA integrity number = 10) was labelled with Cyanine 3 CTP using the Low Input Quick Amp Labelling Kit (Agilent Technologies-5190-2305), and purified using Qiagen’s RNeasy Mini Spin Columns. The quantity of purified Cy3 labelled cRNA was measured using a Nanodrop spectrophotometer. Fragmentation was performed on 600 ng of cRNA from each sample and the fragmented cRNA was then hybridised to Agilent Mouse 8X60K slides (Grid_GenomicBuild, mm9, NCBI37, Jul2007) for 17 hours at 65 °C. Hybridised slides were then washed with Agilent wash buffers and scanned on an Agilent Scanner (Agilent Technologies, G2600D SG12524268), and probe intensities were obtained by taking the gProcessedSignal from the output of Agilent feature extraction software using default settings. Probe annotation and statistical testing was performed using the NIA Array Analysis Tool as described in ref^70^. Significant genes were clustered and heatmap analysis was performed using Morpheus (https://software.broadinstitute.org/morpheus). Gene lists in each cluster were analysed for enriched Gene annotation (GO)-term for Biological Process and Cellular Components using ShinyGO v0.66^71^ to generate lists of functional enrichment.

### Flow cytometry

ESCs were collected and stained with the indicated primary antibody dilutions (Supplementary Table 3) in FACS buffer (10% FBS in PBS) for 15 minutes on ice. The cells were washed three times with FACS buffer and re-suspended in cold FACS buffer containing DAPI (1 μg/mL). If secondary antibodies were required, the cells were further stained with a dilution 1:800 of secondary antibodies for 15 minutes on ice, washed three times with PBS and re-suspended in cold FACS buffer containing DAPI. All experiments included unstained E14Tg2A ESCs as a non-fluorescent control that was used to establish appropriate gates. Flow cytometry was carried out on a BD LSRFortessa (BD Bioscience) and data analysis was performed in FCS Express (De Novo Software).

### *In situ* hybridisation

Whole-mount *in situ* hybridisations (ISH) and sections of catshark embryos were conducted using standard protocols, as described in ref^72^. Catshark breeders were purchased from local professional fishermen and maintained by the Oceanological Observatory Aquariology Service of Banyuls sur Mer (France) with aquatic infrastructure based on national regulations (license number A6601601).

### Structural model prediction by AlphaFold2

Protein sequences of POU5 homologues used for AlphaFold2 structural prediction^48^ are listed in Supplementary Information 3. We performed AlphaFold2 with Colab notebook (Link is noted in Supplementary Table 3). We obtained 3D coordinates, per-residue confidence metric called pLDDT and Predicted Aligned Error from each POU5 structure (shown in Supplementary Information 3). From AlphaFold2 output, non-structural regions including N-/C-terminal domains and a region between α1’-helix of the linker and α1 helix of POU-HD were removed by PyMol^73^ to obtain isolated POU-S-Linker (POU-S-L) and isolated POU-HD. In PyMol, isolated domains were also superimposed to each corresponding domain in mOct4 on *PORE* sequence (PDB: 3L1P^49^). Isolated domains of POU5 protein and *PORE* sequence were saved to obtain new structural model on *PORE* DNA (POU5-*PORE* structure). This combined POU5-*PORE* structures were verified for the clash score (steric clashes) by Phenix^74^ using MolProbity^75^ (Supplementary Information 3-Table 3). The structures with low clash score (<10) were further analysed for H-bonding interaction to *PORE* DNA using ChimeraX^76^ H-bonding prediction parameters included distance tolerance at 0.750Å and angle tolerance at 20.000°.

## Acknowledgements

We thank Gillian Morrison for POU5 plasmids, Yasunori Murakami and Chris Amemiya for assisting with preliminary cyclostome *POU5* analysis, Charlotte Bouleau and Charline Jamin for technical help in *ScPou5f1* ISH, William Hamilton for critical comments and revision, Gelo Victoriano Dela Cruz for flow cytometry technical assistance. We appreciate the contribution of python *Pou5f1* mRNA sequence from Moises Mallo. We also thank the Oceanological Observatory Aquariology Service of Banyuls sur Mer for care of catsharks and EMBRC-France for the support of marine infrastructure. This work was supported by University of Copenhagen studentship (to W.S.), Lundbeck Foundation Grant (to J.M.B, R198-2015-412), région Bretagne doctoral fellowship (to B.G.) and AsymBrain ANR grant (to S.M., ANR-16-CE13-0013-02). Work in the NNF Center for Stem Cell Medicine is funded by the Novo Nordisk Foundation (NNF17CC0027852 and NNF21CC0073729).

## Accession Numbers

The Gene Expression Omnibus (GEO) accession number for the DNA microarray data reported in this study is GSE148167 (LcP1/LcP3/mOct4 rescued ESCs).

## Author contributions

W.S., E.M., M.L., S.M. and J.M.B. designed the study. S.F. obtained tammar wallaby, turtle, coelacanth POU5F1 and POU5F3 coding sequences. S.M. and H.M. obtained hagfish and chondrichthyan POU5 sequences, generated phylogenetic trees and evolutionary rate analyses of POU5. Oct4 null ESC rescue assay and the generation of clonal cell lines was conducted by W.S (for coelacanth, turtle, axolotl, tammar wallaby POU5), E.M. (for catshark, whale shark, little skate and hagfish POU5), H.P. and F.R. (for catshark and chimeric coelacanth-catshark POU5), A.L. (for *Xenopus* XlPOU25 (XlPOU5F3.2) and XlPOU91 (XlPOU5F3.1)), and F.H. (for *Xenopus* POU5 chimeric proteins). W.S. and E.M. performed analysis of POU5 rescued clonal lines by immunofluorescence, qRT-PCR and microarray. J.H. analysed the microarray dataset. M.W., G.M. and W.S. performed AlphaFold2 structural modelling prediction and interpreted the results. B.G. conducted *in situ* hybridizations in catshark embryos. F.S. and S.K. provided the arctic lamprey unpublished transcriptome database, lamprey embryos for POU5 sequence analysis and assisted with lamprey POU5 protein sequence analysis. W.S., M.L., E.M., S.M. and J.M.B. interpreted the results. W.S., E.M., M.L. S.M. and J.M.B. wrote the paper with input from all authors.

## Declaration of Interests

The authors declare no competing interests.

## Supplementary Information

**Supplementary Fig. 1, related to Fig. 2.**
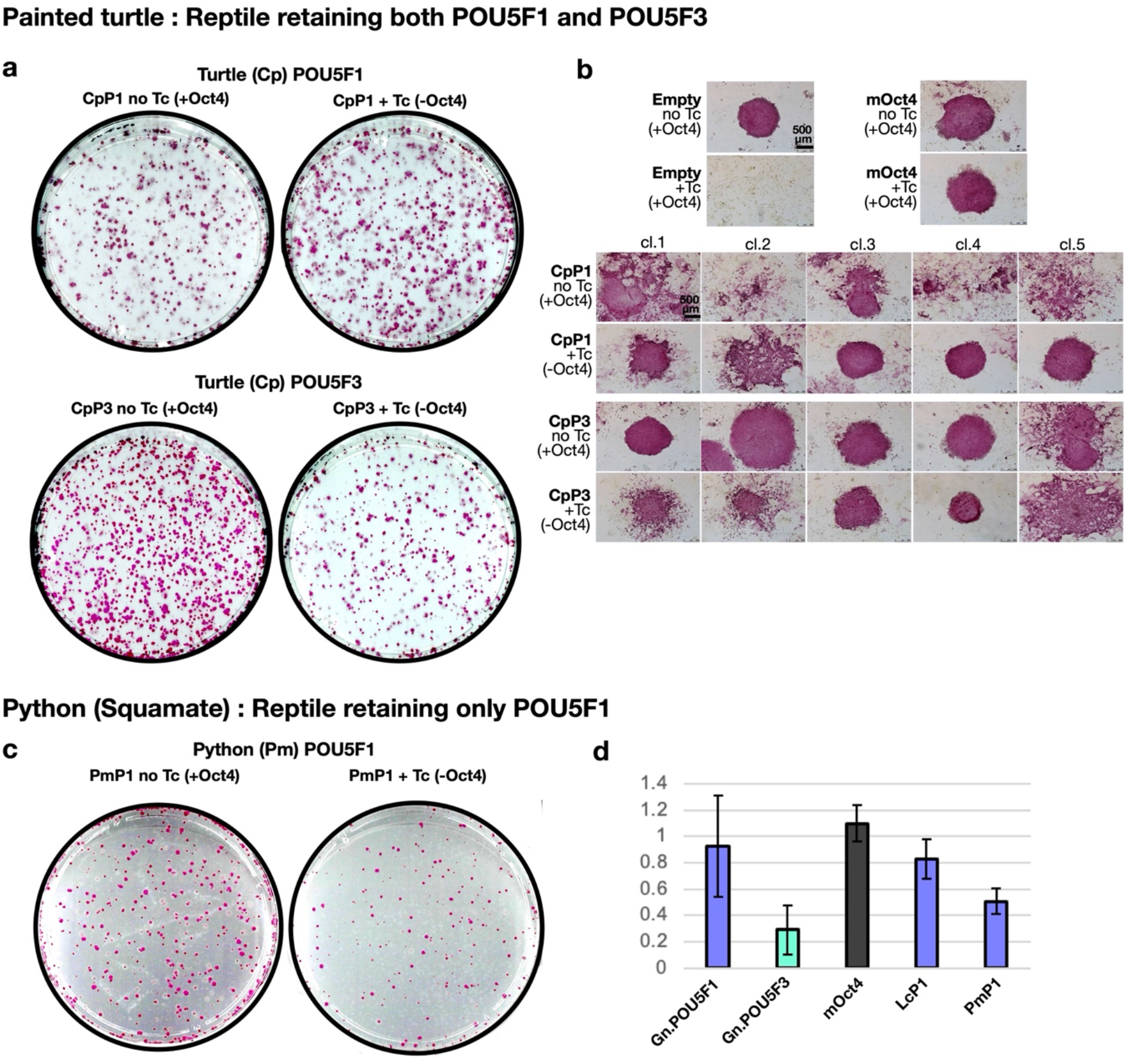
ESC colony morphology of reptilian POU5-rescued Oct4-null ESC cell lines. **a-c,** Alkaline Phosphatase (AP) stainings. **a**, Overview of AP stained colonies (grown in 10cm dishes) of ZHBTc4 cells +tc rescued with painted turtle (*Chrysemys picta*, Cp) POU5F1 and POU5F3. ZHBTc4 cells were electroporated with POU5 vectors and cultured in the absence or presence of tetracycline (= presence of Oct4 expression/absence of Oct4 expression). **b**, Colony morphologies of Cp POU5-rescued ESC colonies. **c**, Overview of AP stained colonies (grown in 10cm dishes) of OCT4-null ESCs rescued with python (*Python molurus*, Pm) POU5F1 (PmP1). **d**, Rescue index of OCT4-null ESCs rescued with PmP1 compared to coelacanth and mouse POU5F1s and average of gnathostome POU5F1s and POU5F3s. Abbreviations: Tc, Tetracycline.

**Supplementary Fig. 2, related to Fig. 3.**
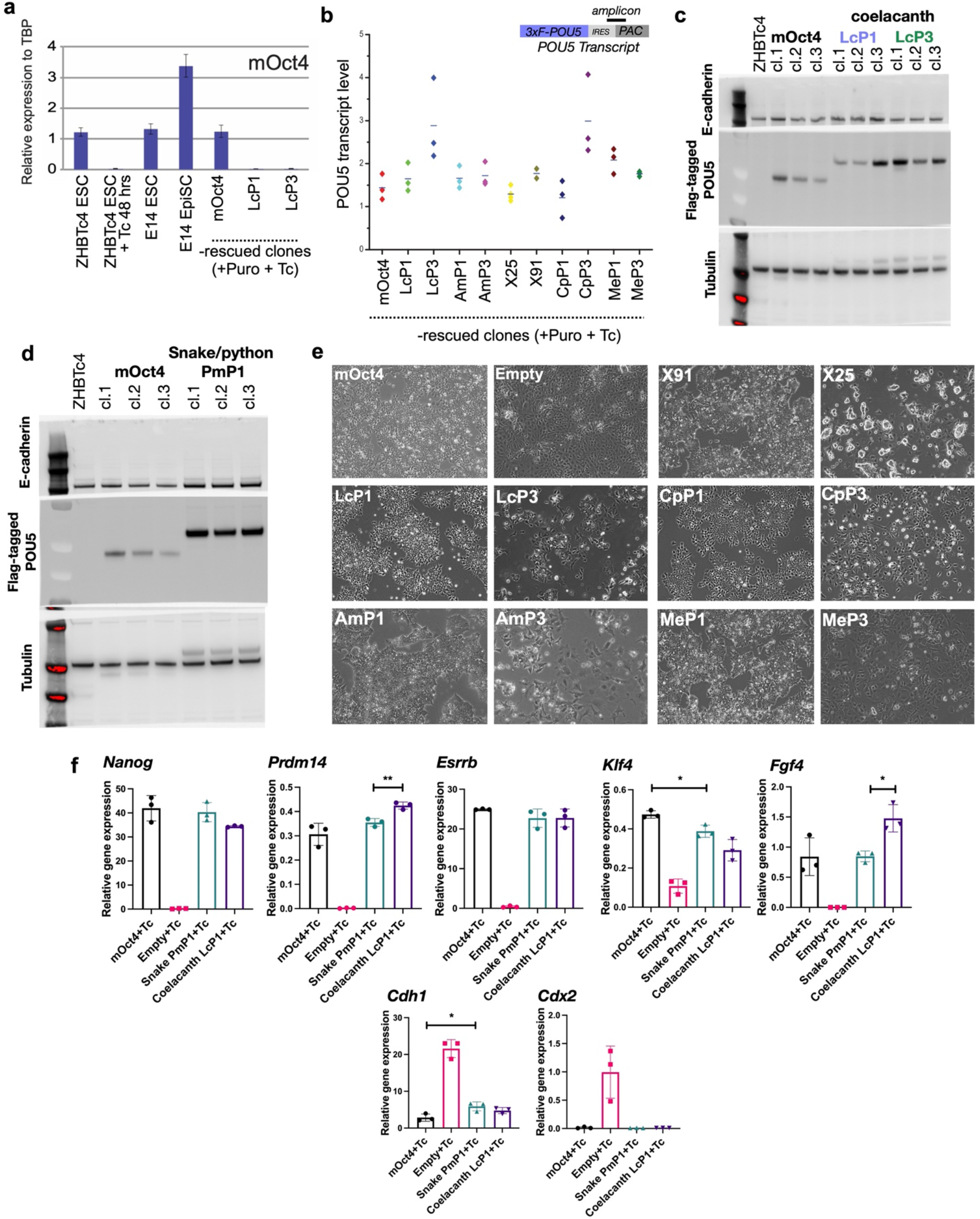
Additional phenotypes of POU5-rescued OCT4-null ESC cell lines. **a**, Relative expression of mouse *Oct4* mRNA, measured by qRT-PCR, to confirm that POU5-rescued ESC clonal lines were maintained solely by transfected POU5 constructs. **b**, qRT-PCR showing expression of the puromycin-resistance gene used to indirectly measure POU5 transcript levels among different POU5-rescued clonal ESC lines. Data represented in the plot show average values of three independent clonal lines with the horizontal bars between diamond symbols representing the mean of these values. **c**, Western blot showing expression of flag-tagged POU5 proteins. **d**, Bright-field images of OCT4-null ESC clonal lines rescued by different POU5 proteins. Colonies from POU5-rescued ESCs were picked and expanded in ESC culture containing tetracycline (Tc) for 6 passages before further cell analysis using immunofluorescence and qRT-PCR (as shown in Fig. 3). OCT4-null cells don’t grow in normal ESC medium because POU5 activity is required for colonies expansion, therefore no rescued colony could be picked from empty vector controls. To obtain ESCs without OCT4 expression, used as control for immunofluorescence and qRT-PCR in Fig. 3, ZHBTc4 cells were electroporated with the empty vector and cultured in the absence of Tc (presence of Oct4 expression). *Oct4* was later removed by addition of tetracycline for four days, to generate differentiated control cultures. **e**-**f**, additional phenotypes of python POU5F1-rescued cell lines. **e**, Western blot showing expression of flag-tagged PmP1 proteins. **f**, qRT-PCR, of OCT4-null ESCs rescued with PmP1, for pluripotency markers (*Nanog*, *Prdm14*, *Esrrb*, *Klf4* and *Fgf4*), cell adhesion (*Cdh1*) and differentiation markers (*Cdx2*). Abbreviations: Empty: empty vector; Puro, Puromycin; abbreviations for POU5 proteins are the same as in Fig. 2. Statistical analyses (Unpaired *t*-Test) were performed on n=3 biological replicates (stable clones expanded from the same POU5 rescue experiment).

**Supplementary Fig. 3, related to Fig. 4.**
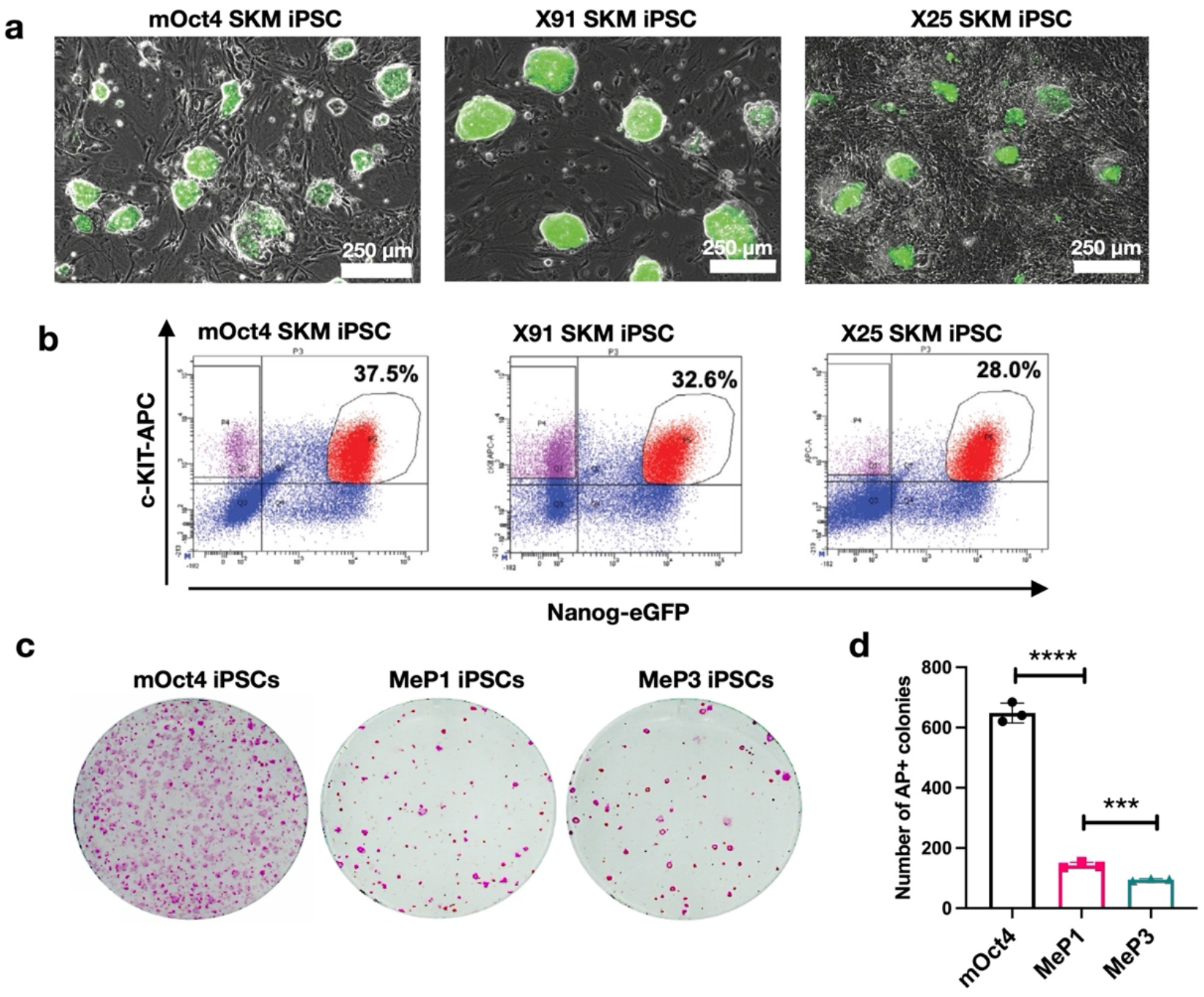
Additional phenotypes of mouse iPSCs generated by different POU5s. **a**, Merged brightfield and Nanog-eGFP (green) expression of iPSCs generated with different POU5 proteins: mOct4, X91 or X25 together with Sox2, Klf4 and c-Myc (SKM). **b,** Nanog-eGFP and a cell surface marker c-KIT profile for assessing naïve pluripotency of iPSCs generated from X91, X25 or mOct4 were analysed by flow cytometry and showed as dot plots. **c**, Overview of AP stainings (purple) of iPSCs generated by tammar wallaby Pou5f1 (MeP1) and Pou5f3 (MeP3) together with mSox2, mKlf4 and mc-Myc. **d**, Reprogramming efficiency of iPSCs generation comparing tammar wallaby POU5F1/3 (MeP1 and MeP3) to mouse Oct4. Statistical analyses (Unpaired *t*-Test) were performed on n=3 biological replicates (3 times of infection (the same batch for virus production) and in 3 different seeding onto irradiated MEF).

**Supplementary Fig. 4, related to Fig. 5.**
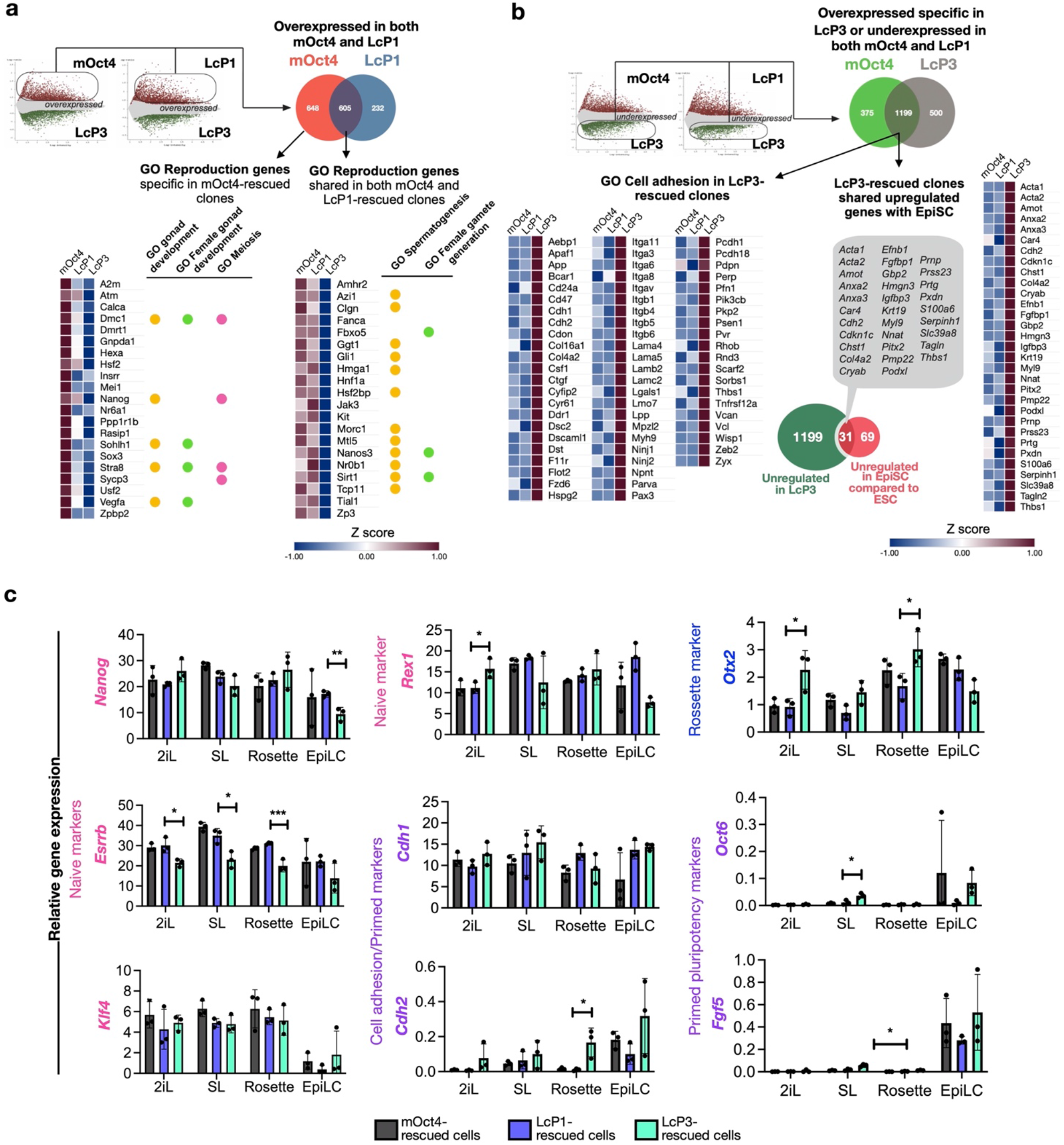
Difference in the gene expression profiles of OCT4-null ESCs supported by LcPOU5F1, mOct4 and LcPOU5F3. **a-b**, Log-ratio plots showing significantly over-expressed (in red) and under-expressed (in green) probes, based on indicated pairwise comparisons (FDR ≤ 0.05). These gene lists were further filtered based on probes corresponding to uniquely annotated genes and with an absolute fold-change cut-off of 2. Differentially expressed genes from both comparisons were used to produce Venn diagrams to further identify common genes expressed in both mOct4 and LcP1-supported ESCs or specific to LcP3-supported ESCs. **a**, Over-expressed genes (605 genes) in common to mOct4 and LcP1-supported cells and over-expressed genes (648 genes) specific to mOct4-supported cells were analysed by the GO term analysis tool ShinyGO (FDR ≤ 0.05 and GO terms with more the 5 genes are shown). Expression of genes in the ‘Reproduction’ GO term were enriched in both mOct4- and LcP1-rescued clones were shown as heatmaps. **b**, Over-expressed genes (1199) in LcP3-supported cells, but down-regulated in both mOct4- and LcP1-suppoted cell lines were analysed by GO term analysis (Left panel). Heatmaps of genes from the ‘biological adhesion’ GO term were enriched in the LcP3-rescued clones (Right panel). The set of genes specifically up-regulated in LcP3-supported ESCs (1199 genes) were compared to top 100 most significant genes up-regulated in EpiSC when compared to naïve ESCs^47^. List of genes shared in both EpiSC and LcPOU5F3-rescued clones were used to construct a heatmap to visualise their expression. **c**, Relative gene expression of naïve, rosette and primed pluripotency markers and cell adhesion markers of mOct4, LcP1 or LcP3-rescued cells under SL/Rosette/EpiLC culture conditions were analysed by qRT-PCR, as shown in Fig. 5e. Statistical analyses (Unpaired *t*-Test) were performed on n=3 biological replicates (stable clones expanded from the same POU5 rescue experiment).

**Supplementary Fig. 5, related to Fig. 6.**
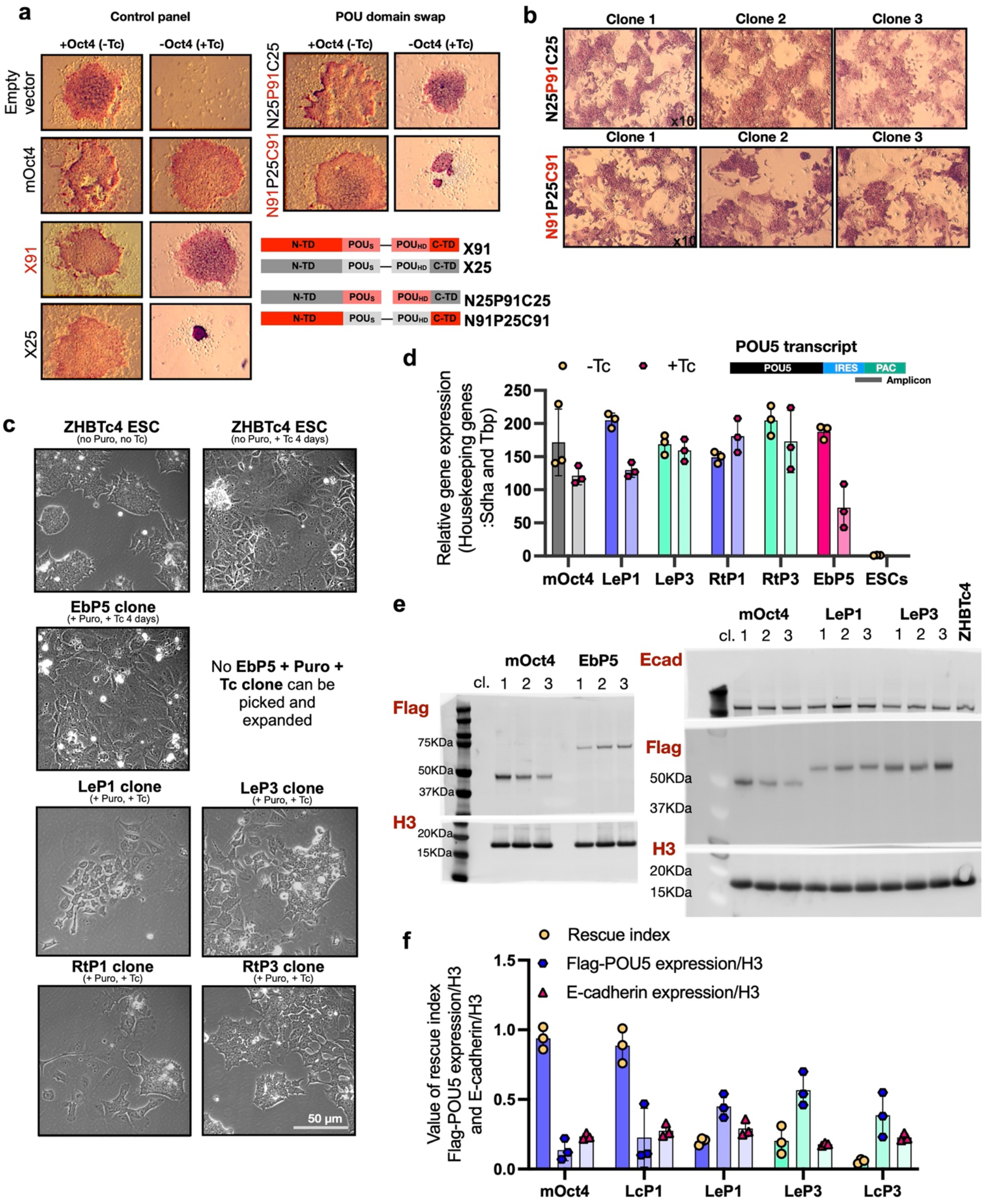
Chondrichthyans POU5 rescue activity and functional domains. **Mapping the functional domains of POU5 proteins responsible for rescue of OCT4-null ESCs.** As *Xenopus* proteins have a similar coding sequence, but differ dramatically in their rescue activity, we used these proteins to initially map the functional domains responsible for rescue. **a**, Morphology of chimeric POU5-rescued ESC colonies stained for alkaline phosphatase (AP, purple) activity. OCT4-null ESCs were rescued by X91-X25 chimeric proteins and data for the POU domain swap is shown. X91 is as effective as mOct4 at supporting ESC self-renewal. Swapping POU domain of X91 into X25 improved rescue activity of X25 while replacement of POU domain in X91 from X25 diminished the capacity of X91 to support ESCs. **b**, Stable OCT4-null ESCs cultures rescued with X91-X25 chimeric proteins were also stained for AP activity. **Chondrichthyan, but not cyclostome POU5 proteins rescue Oct4 phenotypes.** **c-f**, Analysis of expanded, stable OCT4-null ESC clonal lines rescued by Chondrichthyan POU5 proteins. As Hagfish POU5 (EbP5) cannot rescue, EbP5 expressing colonies were picked and expanded in the absence of Tc, then treated with Tc for four days to remove OCT4 before imaging. **c**, Bright-field images of expanded colonies. **d**, Expression of the puromycin-resistant gene used to indirectly measure POU5 transcript levels between different POU5-rescued clonal lines, measured by qRT-PCR. **e**, Western blots showing levels of FLAG tagged proteins, representing levels of POU5 and E-cadherin (Ecad), in three clones per rescued line with H3 as a loading control. **f**, Bar chart showing the comparison between the rescue index for each species and the relative POU5 and E-cadherin protein levels from stable rescued lines (see Supplementary fig. 2). Individual data points in the bar graphs represent values from three independent clonal lines.

**Supplementary Fig. 6, related to Fig. 7.**
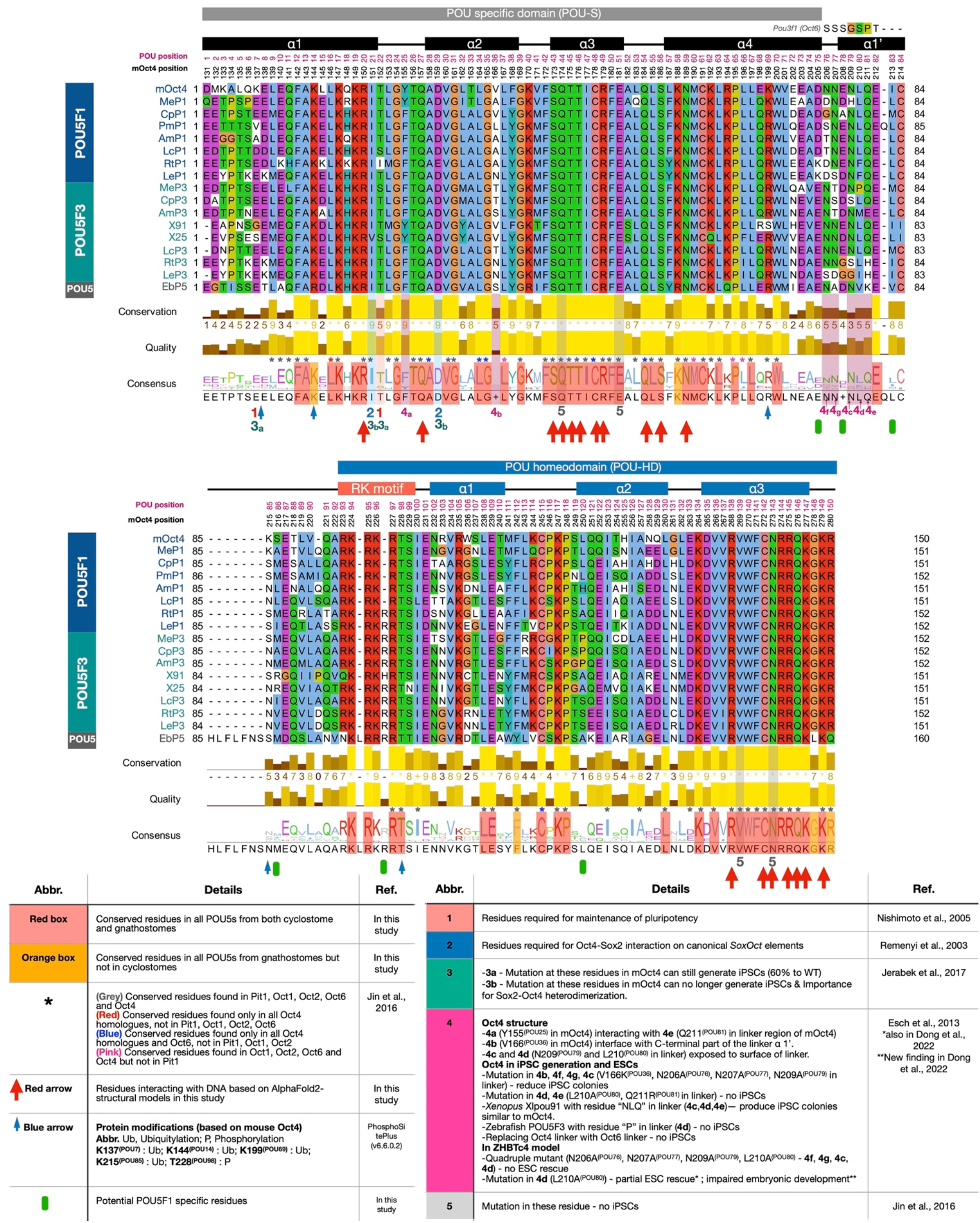
Multiple sequence alignment of POU5 proteins used in Oct4 rescue assays. Protein sequences from POU5s were aligned using MUSCLE and visualized in Jalview^77^. Consensus logo indicating conservation level is shown below the alignment. The uniqueness of each residue among POU5 proteins was identified based on literature review and AlphaFold2-based protein-DNA interaction in this study. Residue conservation among POU proteins (PIT1, OCT1, OCT2, OCT4 and OCT6) is marked with coloured asterisks on the top of consensus logo and level of conservation is shown with coloured boxes over the consensus sequence, explanation below the alignment (lower left). POU domain-DNA interactions from AlphaFold2-based mOct4 structural models using ChimeraX are noted with red arrows and residues with reported post-translational modifications are noted with blue arrows. Literature references for functional analysis of residues are noted under the consensus logo as a table (1-5^49–54,78,79^), with the role of the residue detailed below the alignment (lower right).

**Supplementary Fig. 7, related to Fig. 7.**
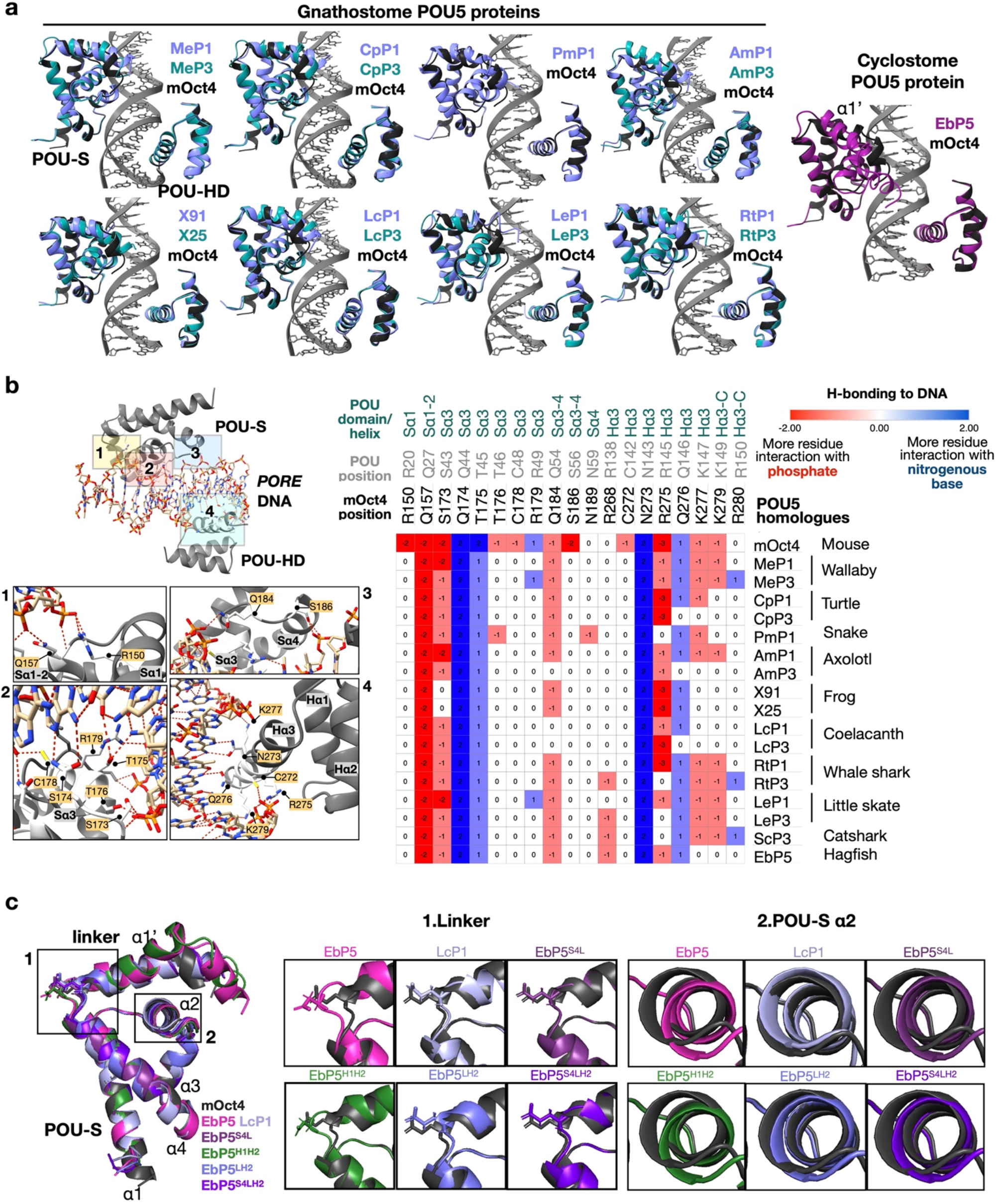
AlphaFold2-based structural models for POU5 proteins used in Oct4 rescue assays. **a**, Predicted structures for all gnathostome and cyclostome POU5 homologues analysed for rescue potential in OCT4-null ESCs (Figures 2 and 6) using AlphaFold2. The POU-S domain together with the linker helix (POU-S-L) and the POU-HD of each pair of POU5 homologues are shown in superimposition to mOct4 (black) on the *PORE* DNA element (PDB ID: 3L1P) and visualized by ChimeraX. **b**, POU-DNA H-bond interactions were predicted by ChimeraX. The number of H-bonds from each residue interacting with nitrogenous base (blue, positive value) and/or phosphate (red, negative value) of the DNA are shown as a heatmap, produced by Morpheus^80^. Zoom-ins on the mOct4-*PORE* interaction regions to visualize specific residues (boxes 1-4). **c**, AlphaFold2 predicted structures for chimeric proteins. Superimposition of POU-S domains from mOct4, LcP1, EbP5 and the four chimeras (left side), with zoom-in 1 focusing on the E208^(POU78)^ residue found altered in the bend between the POU-S *α*4 and the Linker *α*1’ and the zoom-in 2 focusing on the positioning of POU-S *α*2, each structure is compared to mOct4 (dark grey).

**Supplementary Fig. 8, related to Fig. 7.**
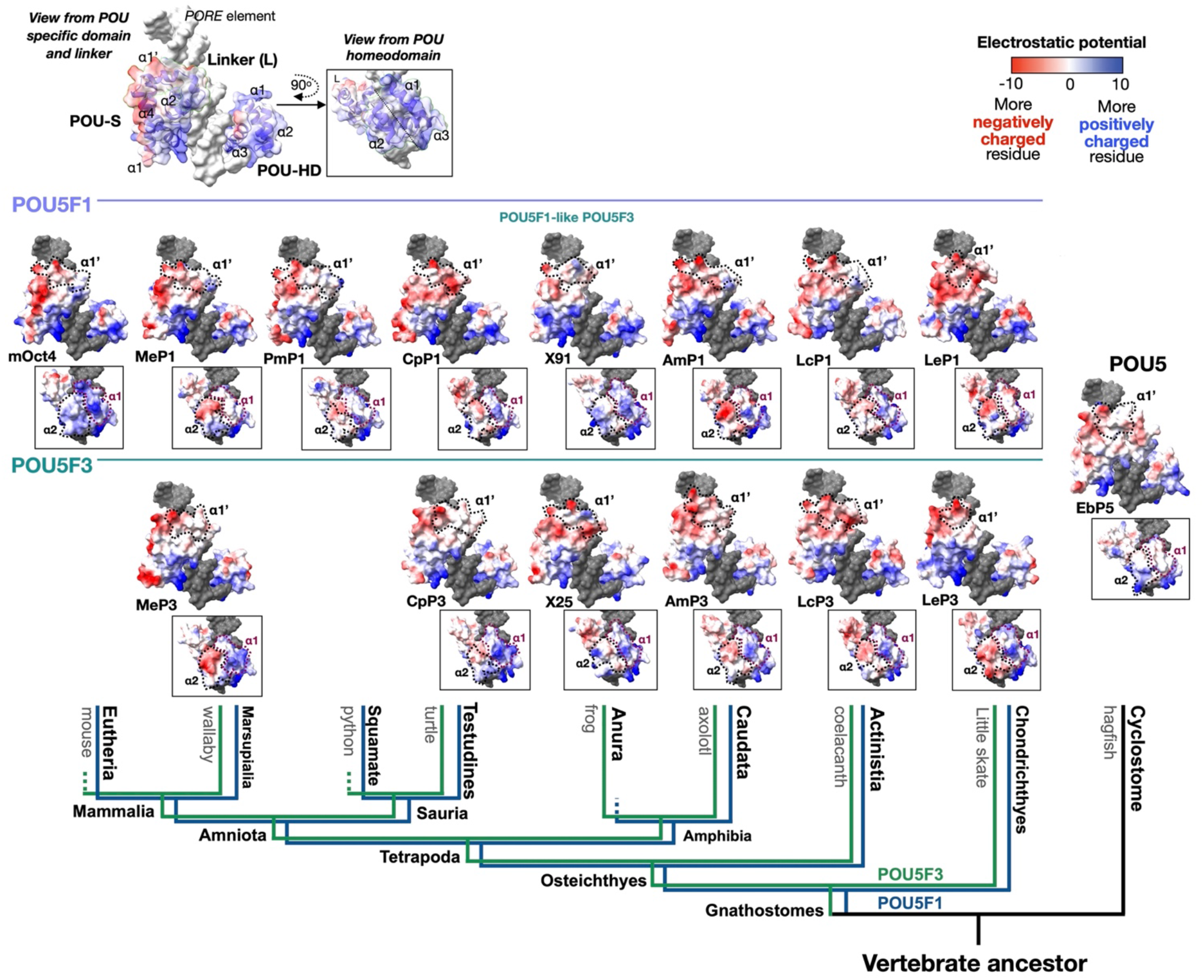
Electrostatic potential of all AlphaFold2-based POU5 structural models. Predicted electrostatic surface potentials of POU5 domains on the *PORE* DNA element for all gnathostome and cyclostome POU5 homologues analysed. Surface charges, determined by ChimeraX, with negatively charged areas shown in red and positively charged in blue. Each POU5 protein has an orientation focusing on the POU-S-L (top) and the POU-HD (boxed below) with a description of the views above. The structures are organized based on the phylogeny tree shown at the bottom.

**Supplementary Fig. 9, related to Fig. 9.**
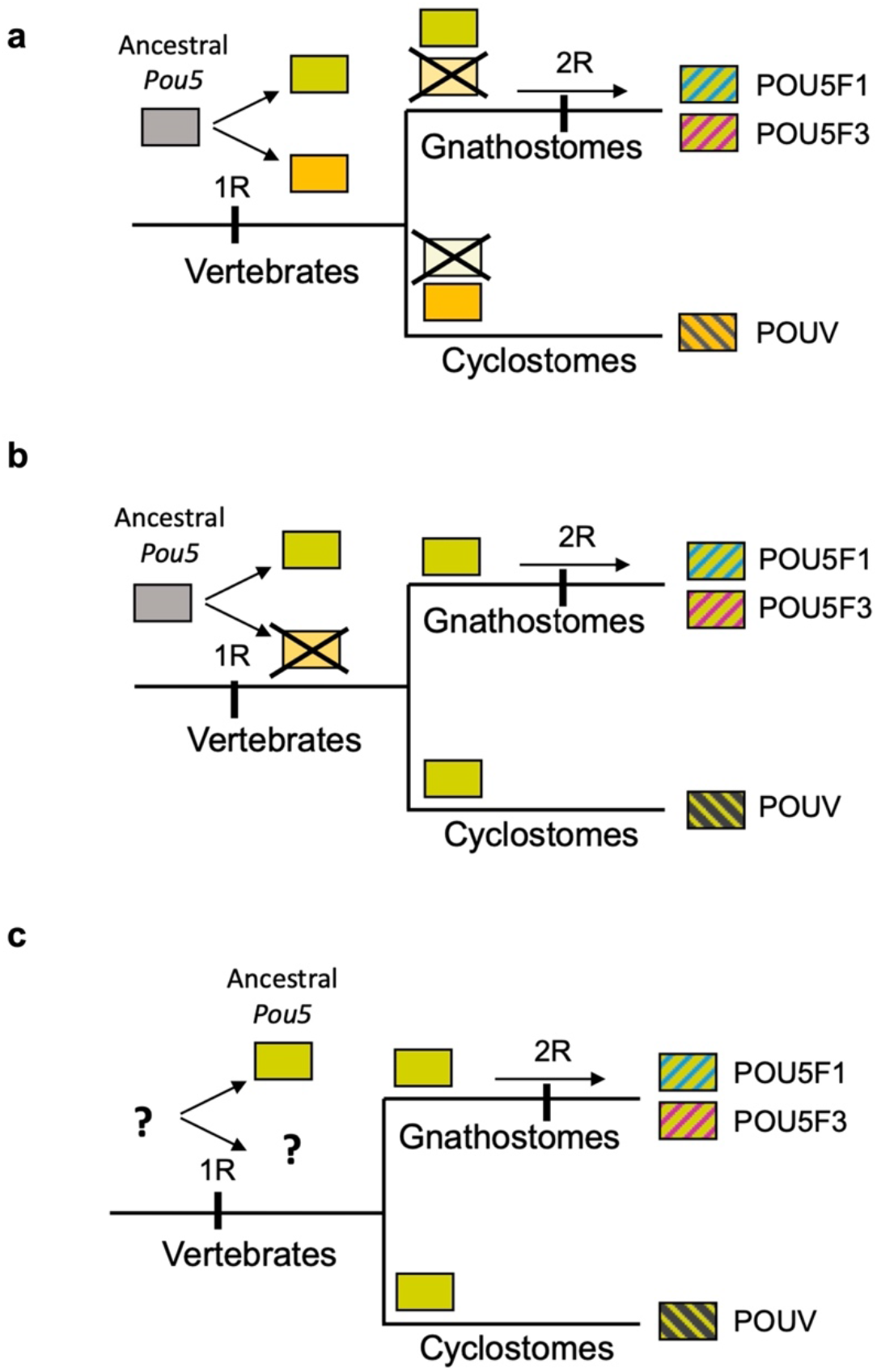
Evolution of the POU5 family prior to the gnathostome radiation. Our data suggest that the *Pou5f1*/*Pou5f3* duplication was part of the second round of vertebrate WGD (whole genome duplications), recently proposed to have followed the split between gnathostomes and cyclostomes^58^. This hypothesis is consistent with both the detection of extensive syntenies found between the paralogous loci and the monophyly of gnathostome *Pou5* genes, POU5F1 and F3 proteins appearing more similar to each other than to their cyclostome counterparts. Different possibilities consistent with these observations are shown. **a**-**b**, Early rise of *Pou5* genes in the vertebrate lineage, prior to the first round of WGD (1R) with (**a**) differential losses of either one of 1R duplicates between gnathostomes and cyclostomes or (**b**) loss of one 1R duplicate prior to the gnathostome-cyclostome split. **c**, variation of **b**, with the rise of *Pou5* genes following the 1R WGD.

**Supplementary Table 1, related to Fig. 1 Accession numbers of the sequences used in synteny and phylogenetic analyses and predictions of *POU5* coding sequences from genomic databases.**

**Supplementary Table 2, related to Fig. 5 Gene Expression Data and Gene Ontology Analysis based on the transcriptomic comparisons of coelacanth (Lc) POU5-rescued and mOct4-rescued ESC lines.**

**Supplementary Table 3. Resources used in and generated by this study.** Reagents, plasmids, cell lines, antibodies, software and deposited data are listed, together with the places they can be obtained from. For all antibodies, the dilution used is indicated. Catalogue numbers are listed where relevant.

**Supplementary Dataset 1, related to Fig. 1 Alignment POU5_Ali.fst.**

Alignment of the POU specific domain (POU_S_), Linker domain and POU homeodomain (POU_HD_) of POU5 proteins used to generate the consensus sequence of POU5 protein (Fig. 1a) and the phylogenetic tree showing the evolutionary rates of vertebrate POU5 proteins (Fig. 1c).

**Supplementary Information 1 Syntenic analysis, phylogenetic analysis and the estimation of rate of POU5 protein evolution.**

**Supplementary Information 2 Additional information for figures.**

**Supplementary Information 3 Structural models of POU5 homologues from AlphaFold2.**

## Notes

### Competing Interest Statement

The authors have declared no competing interest.

