## Supplementary Information 1 for "Evolutionary Origin of Vertebrate OCT4/POU5 Functions in Supporting Pluripotency"

### Supplementary Information 1 (Related to Fig. 1)

**<u>Supplementary information related to Fig. 1a.</u> Alignment of POU5 predicted sequences: (a) and (b) chondrichthyan full-length POU5F1 and POU5F3 respectively, (c) cyclostome full-length POU5 and (d) vertebrate POU5 over the POU domain, linker and homeodomain.** Amphioxus (*Branchiostoma floridae*) POU2 and POU3 are included as outgroups for comparisons in (d). A dot denotes a missing amino acid, an X undetermined sequence. In (d), POU5 synapomorphies, as identified in Gold et al. 2013, are boxed and shown in red, with the ancestral residue in black. The N-terminal conserved domain (NTD), POU specific domain and POU homeodomain are shown respectively in blue, grey and green. Residues conserved between gnathostome POU5F1 and POU5F3 but not with cyclostome POU5 in the C-terminal part of the protein are shaded in magenta in (a-c) and sequences, which can be aligned between both classes, are in bold characters. *Amblyraja radiata* sequences were retrieved from a genome assembly, which does not meet VGP quality standard and were edited to remove putative frame shifts in the sequence (see Supplementary Table 1). Species abbreviations: Ar, *Amblyraja radiata*; Cm, *Callorhinchus milii*; Cpu, *Chyloscyllium punctatum*; Hz, *Heterodontus zebra*; Le, *Leucoraja erinacea*; Ok, *Okamejei kenojei*; Rh/Rt, *Rhincodon typus*; Sc, *Scyliorhinus canicula*; St, *Scyliorhinus torazame*.

**a**

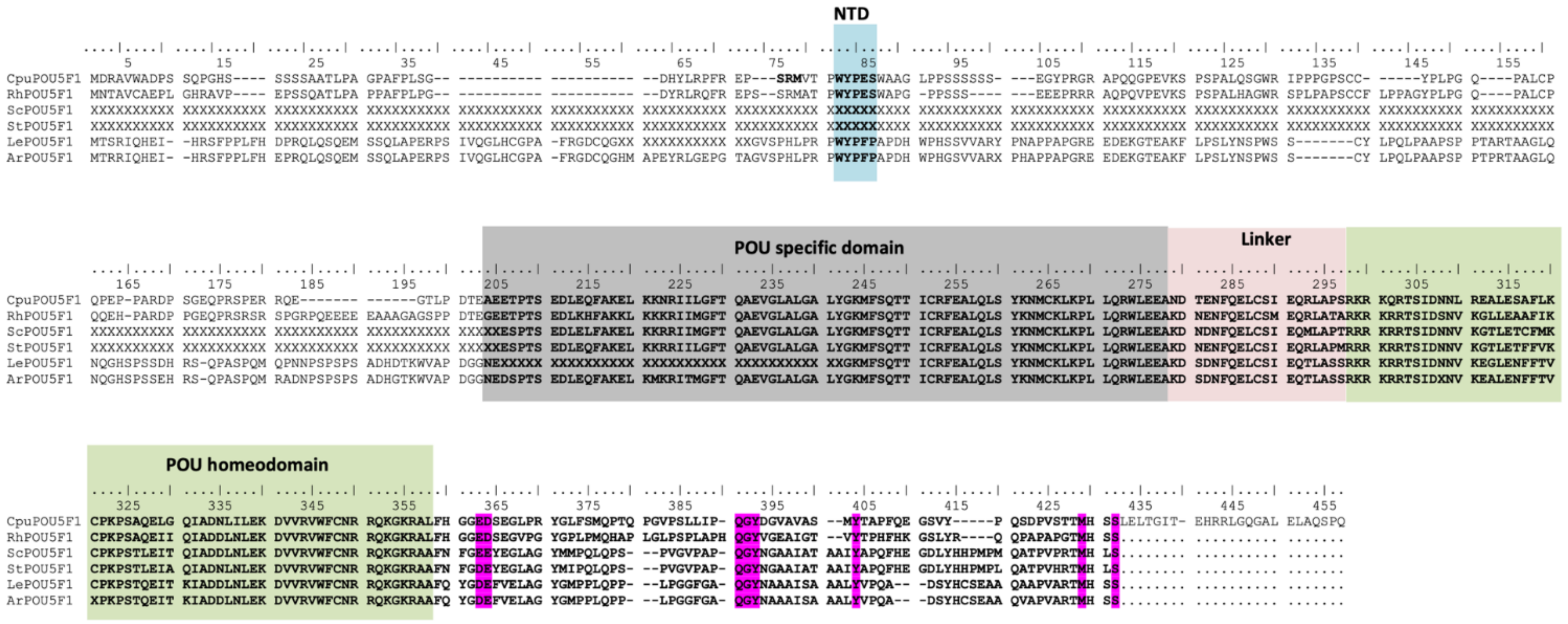

**b**

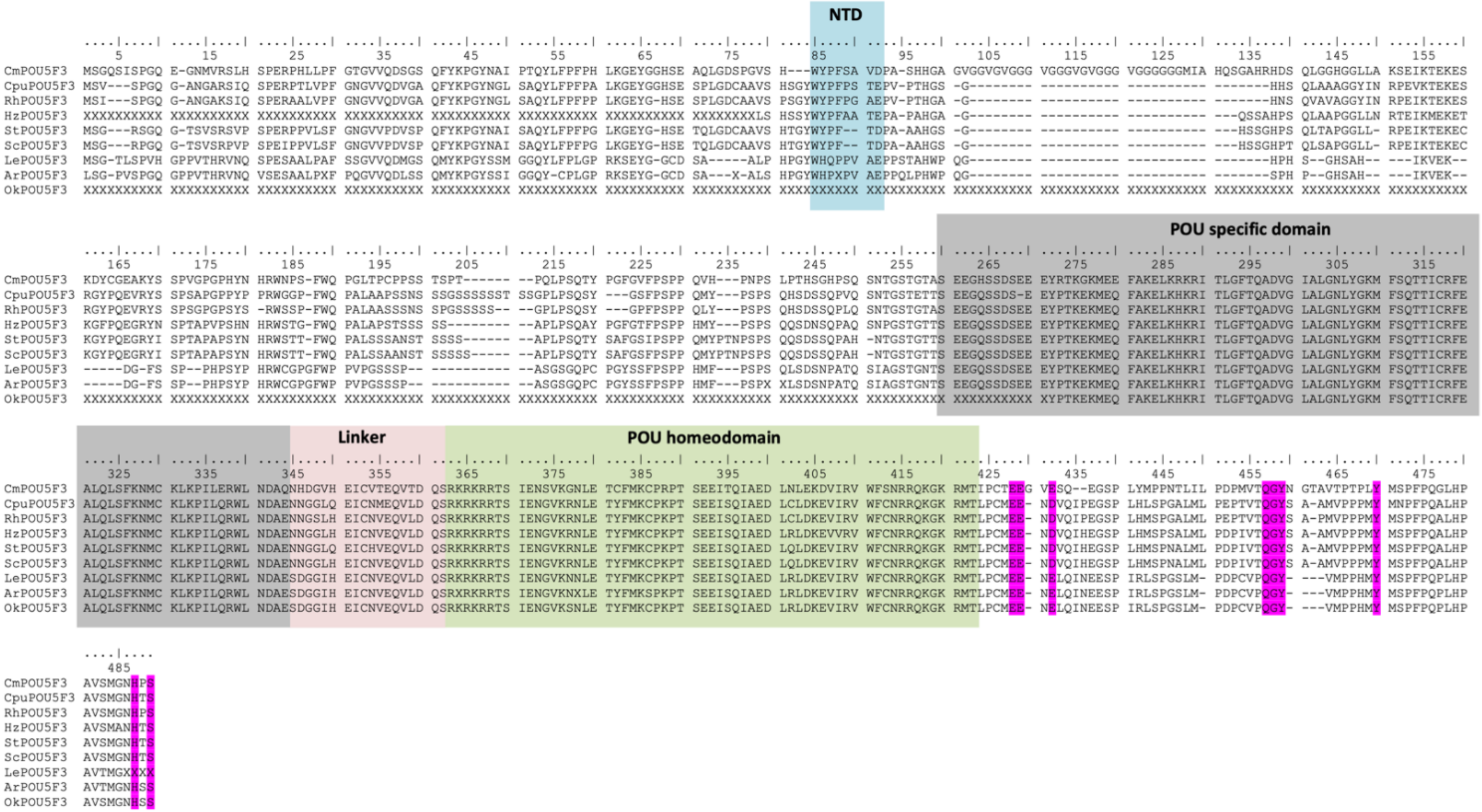
NTD
POU specific domain
Linker
POU homeodomain
CmPOU5F3
CpuPOU5F3
RhPOU5F3
HzPOU5F3
StPOU5F3
ScPOU5F3
LePOU5F3
ArPOU5F3
OkPOU5F3

c

```
                                                                                                       NTD
         5          15         25         35         45         55         65         75         85         95         105        115        125        135        145        155
EbPOU5   MSSLSIKVDA ANEL-QLMYE SYSEASAAQS SVVRPYMEQA AQQRGACCVP GQGSGHGHVH LVPISPLSLG YSGELYEPSL QCPSRAHGAV PGPAWYAYPG PSAAGSEAAQ AAASWAAARG VEIKCEEGVG VGEHVESEVK RIRYQDFKYG AVYGSGYVLP
PmPOU5   METQSLQTSA L-DIPQSGPS LDNRF----- RAYGPHQGVS VQRSPHEQTA GAGGQMMDAF NNSSLSSSSS SSSSSSSS-L PAPGTPLGHR MGTNWFATIN IP----DYYP PQPQE--YHH QQQQQQQQQQ ---------- ---EGPPRHP STYGSTVATS
LjPOU5   METQSLQTSA L-DIPQPGPS LDNRF----- RAYGPHPGVS AQSSPPEEPA GAGGQVMDAF NNSSLSSSSS SSLSSSSSLL QAPDTPLGHR MGTNWFATIN IP----DYYS LQLQQQQYHQ QQYHQQQQQY HQQQQQYHQQ QQQEGPPRHP STYGPTVATT

         165        175        185        195        205        215        225        235        245        255        265        275        285        295        305        315
EbPOU5   NAAMHGGHGQ QNPHSLSHLN LHPHLQLNPS PHLQPNPNLH LQLNPNSHLQ PSSHLQPNPS SHLQPNPNSH LQLSSHLQQN SHLQPSSHLQ PNSHLQPNPN SHLQPNPNSH LQPNTNSHLQ PNPNSHLQPN PNSHLQPNPN SHLQLSSHSQ QNPLSLLNVH
PmPOU5   AVSRATVHNN GVATDETLAH QQHQQHLHQQ HQQQQHQQHQ QQHQQRRQRQ LAPRSIGYAA PFPSGLGDAG SPADDLAVAG TSASFLASSV PCQPPPP--- ---HHHHHH- ------GWQN LPVPPPAASR YQESVVVIVD DDDDDDDDDS SG-------D
LjPOU5   AGSLVAVHNN GVATDETLAY QHHQQQQHHH QQQHHHQQQ- -----RRQRQ LTPRSIGFAA PFLAGLG-AE SPDDDLAVAS TSARFLASTV PCQPPPPPPP PPLHHHQHHH QHHQHQGWPG LPPA-PAASR FEESVVIDIV DDDDDGSSSS SGGGGGGGGD

         245        255        265        275        285        295        305        315        325        335        345        355        365        375        385        395
EbPOU5   SHLQPSSHLQ PNSHLQPNPN SHLQPNPNSH LQPNTNSHLQ PNPNSHLQPN PNSHLQPNPN SHLQLSSHSQ QNPLSLLNVH PQAYPHSQSP APTPLAAALA STQSPGASSG TEDGGCARER SRNSEHDLGS ------GDE- --G-TISSET LAQFARDLKH
PmPOU5   TSASFLASSV PCQPPPP--- ---HHHHHH- ------GWQN LPVPPPAASR YQESVVVIVD DDDDDDDDDS SG-------D IVVSVSDDEC HGAMMQQQQQ QQ-LLMHHLL QQQQQQQLPH QPHHGQRPGY EWHGAMGEAA --S-DGNIDE LARFAKELKV
LjPOU5   TSARFLASTV PCQPPPPPPP PPLHHHQHHH QHHQHQGWPG LPPA-PAASR FEESVVIDIV DDDDDGSSSS SGGGGGGGGD TVVSVSDDEC HGTMMHQQQQ QQLLLMHHLL RQQQQQQQPR QLHHGQRPAY EWHGALGEVG --S-DGNIDE LARFAKELKM

                         POU specific domain                                           Linker                                                  POU homeodomain
         405        415        425        435        445        455        465        475        485        495        505        515        525        535        545        555
EbPOU5   KRITLGFTQA DVGVALGSLY GRIFSQTTIC RFEALQLSYR NMCKLQPLLE RWMIEAENAD NVKEVCHLFL FNSSMD---- ---------- ---------- ---QSLANVN KLRRRRTTIE NGVRDTLEAW YLVCSKPSAK EIARIAGELN LDKEVVRVWF
PmPOU5   KRVNLGFTQA EMGLSIGSLC GRVFSQTTIC RFEALQLSQR NLCKLQPLFK LWMDDVGVNG GSGGVSSVGG GVGGK----- ---------- ---------- ---TPREVGY RMRKKRTYFA SYTRMQLEAY YNVCSKPNMS AIASIAQQLK LENSVVRLWF
LjPOU5   KRVNLGFTQA EMGISIGSLC GRVFSQTTIC RFEALQLSQR NLCKLQPLFK LWMEDVQSNG SISSNGGNGG NGGNGGNGGN GGNGGNGGND TKENAAAGQT LSKTPREVVQ RIRKKRTYFA SYTRMQLEAY YNVCSKPNMS AIASIAQHLK LENNVVRLWF

         565        575        585        595        605        615        625        635        645        655        665        675        685        695        705        715
EbPOU5   CNRRQKLKQL SPPLPKEEE- -PEMSSQHQL MPHIQ----- ---------- ---------- ---EFGTLPT SSSSPAYDIH TYNMQASM-N APILPS.... .......... .......... .......... .......... .......... ........
PmPOU5   SNRRQKCRKA STSLADNNVP QFHANGQNGP GAAAPVAVAR DNIDTDGSTG GV-------- -ETVAA--MA YDGAFAACHR -GGVCKMETS ASSCLRSAGA PRGADAHGFS GYG------- -GGGDDGG RDGNGDGSDLFP TAVHTCLRSF AGSGLNDT
LjPOU5   SNRRQKCKKA CTSFPDDNVP PFHANGHNGP GAAAPVVIVH DNIDG--STG GGGGGGGGGG GDIVAAAAMA YDGAFAACHD EGGVCKMETS DDTCLRGAGA PRSGDAGACG FNGDAGYDGG DGGGDDGGRD GSGDGSNLFP TAVHTYMRSF AGSGLNDT
```

d

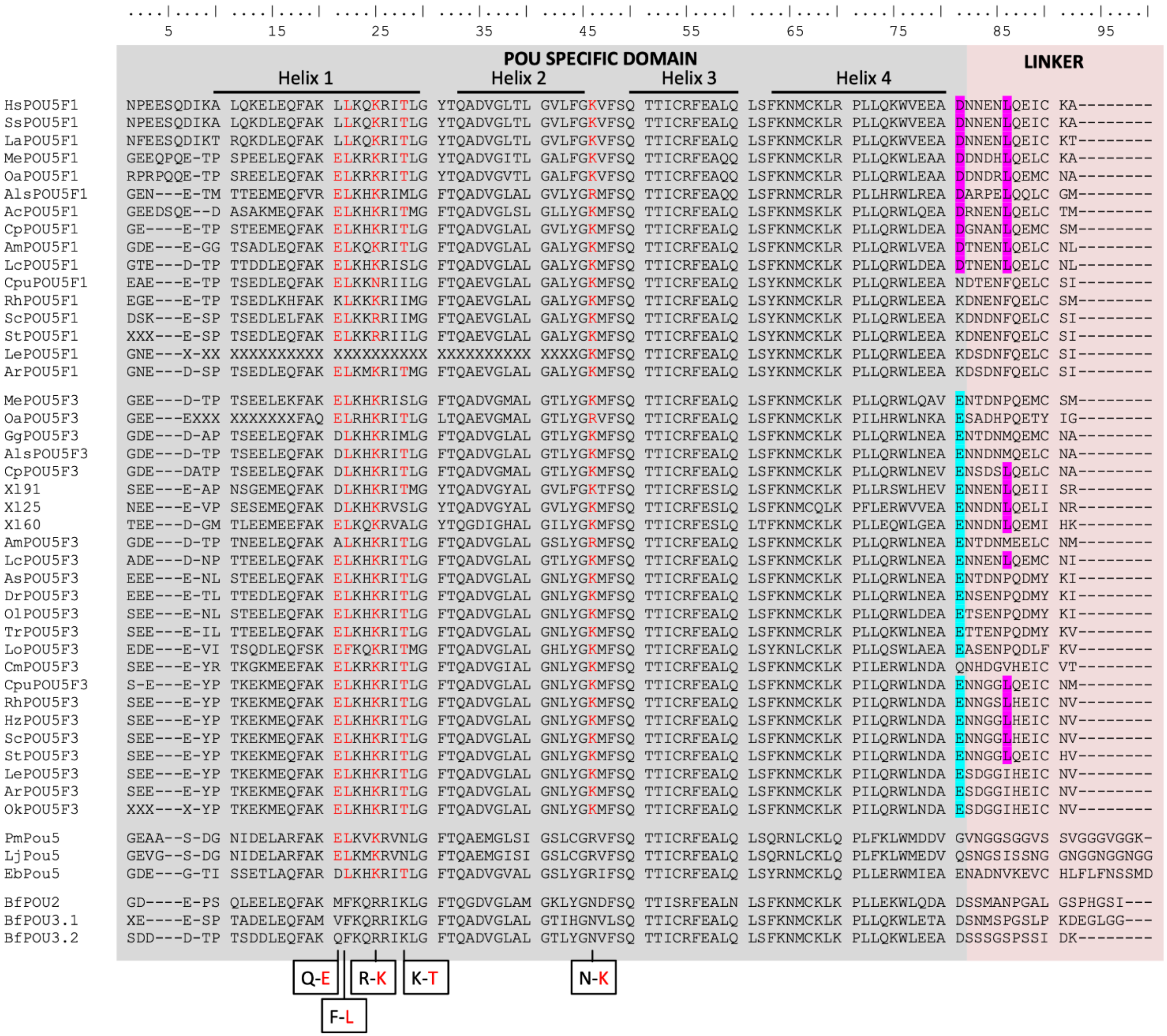
5 15 25 35 45 55 65 75 85 95
POU SPECIFIC DOMAIN
LINKER
Helix 1
Helix 2
Helix 3
Helix 4
HsPOU5F1 NPEESQDIKA LQKELEQFAK LLKQKRITLG YTQADVGLTL GVLFGKVFSQ TTICRFEALQ LSFKNMCKLR PLLQKWVEEA DNNENLQEIC KA--------
SsPOU5F1 NPEESQDIKA LQKDLEQFAK LLKQKRITLG YTQADVGLTL GVLFGKVFSQ TTICRFEALQ LSFKNMCKLR PLLQKWVEEA DNNENLQEIC KA--------
LaPOU5F1 NFEESQDIKT RQKDLEQFAK LLKQKRITLG YTQADVGLTL GVLFGKVFSQ TTICRFEALQ LSFKNMCKLR PLLQKWVEEA DNNENLQEIC KT--------
MePOU5F1 GEEQPQE-TP SPEELEQFAK ELKRKRITLG YTQADVGITL GALFGKVFSQ TTICRFEAQQ LSFKNMCKLR PLLQKWLEAA DDNDHLQELC KA--------
OaPOU5F1 RPRPQQE-TP SREELEQFAK ELKRKRITLG YTQADVGVTL GALFGKVFSQ TTICRFEAQQ LSFKNMCKLR PLLQRWLEAA DDNDRLQEMC NA--------
AlsPOU5F1 GEN---E-TM TTEEMEQFVR ELKHKRIMLG FTQADVGLAL GVLYGRMFSQ TTICRFEAQQ LSFRNMCRLR PLLHRWLREA DARPELQQLC GM--------
AcPOU5F1 GEEDSQE--D ASAKMEQFAK ELKHKRITMG FTQADVGLSL GLLYGKMFSQ TTICRFEALQ LSFKNMSKLK PLLQRWLQEA DRNENLQELC TM--------
CpPOU5F1 GE----E-TP STEEMEQFAK ELKHKRITLG FTQADVGLAL GVLYGKMFSQ TTICRFEALQ LSFKNMCKLR PLLQRWLDEA DGNANLQEMC SM--------
AmPOU5F1 GDE---E-GG TSADLEQFAK ELKQKRITLG FTQADVGLAL GALYGKMFSQ TTICRFEALQ LSFKNMCKLR PLLQRWLVEA DTNENLQELC NL--------
LcPOU5F1 GTE---D-TP TTDDLEQFAK ELKHKRISLG FTQADVGLAL GALYGKMFSQ TTICRFEALQ LSFKNMCKLK PLLQRWLDEA DTNENLQELC NL--------
CpuPOU5F1 EAE---E-TP TSEDLEQFAK ELKKNRIILG FTQAEVGLAL GALYGKMFSQ TTICRFEALQ LSYKNMCKLK PLLQRWLEEA NDTENFQELC SI--------
RhPOU5F1 EGE---E-TP TSEDLKHFAK KLKKKRIIMG FTQAEVGLAL GALYGKMFSQ TTICRFEALQ LSYKNMCKLK PLLQRWLEEA KDNENFQELC SM--------
ScPOU5F1 DSK---E-SP TSEDLELFAK ELKKRRIIMG FTQAEVGLAL GALYGKMFSQ TTICRFEALQ LSYKNMCKLK PLLQRWLEEA KDNDNFQELC SI--------
StPOU5F1 XXX---E-SP TSEDLEQFAK ELKKRRIILG FTQAEVGLAL GALYGKMFSQ TTICRFEALQ LSYKNMCKLK PLLQRWLEEA KDNENFQELC SI--------
LePOU5F1 GNE---X-XX XXXXXXXXXX XXXXXXXXXX XXXXXXXXXX XXXXGKMFSQ TTICRFEALQ LSYKNMCKLK PLLQRWLEEA KDSDNFQELC SI--------
ArPOU5F1 GNE---D-SP TSEDLEQFAK ELKMKRITMG FTQAEVGLAL GALYGKMFSQ TTICRFEALQ LSYKNMCKLK PLLQRWLEEA KDSDNFQELC SI--------
MePOU5F3 GEE---D-TP TSEELEKFAK ELKHKRISLG FTQADVGMAL GTLYGKMFSQ TTICRFEALQ LSFKNMCKLK PLLQRWLQAV ENTDNPQEMC SM--------
OaPOU5F3 GEE---EXXX XXXXXXXFAQ ELRHKRITLG LTQAEVGMAL GTLYGRVFSQ TTICRFEALQ LSFKNMCKLK PILHRWLNKA ESADHPQETY IG--------
GgPOU5F3 GDE---D-AP TSEELEQFAK DLKHKRIMLG FTQADVGLAL GTLYGKMFSQ TTICRFEALQ LSFKNMCKLK PLLQRWLNEA ENTDNMQEMC NA--------
AlsPOU5F3 GDE---D-TP TSEELEQFAK DLKHKRITLG FTQADVGLAL GTLYGKMFSQ TTICRFEALQ LSFKNMCKLK PLLQRWLNEA ENNDNMQELC NA--------
CpPOU5F3 GDE---DATP TSEELEQFAK DLKHKRITLG FTQADVGMAL GTLYGKMFSQ TTICRFEALQ LSFKNMCKLK PLLQRWLNEV ENSDSLQELC NA--------
X191 SEE---E-AP NSGEMEQFAK DLKHKRITMG YTQADVGYAL GVLFGKTFSQ TTICRFESLQ LSFKNMCKLK PLLRSWLHEV ENNENLQEII SR--------
X125 NEE---E-VP SESEMEQFAK DLKHKRVSLG YTQADVGYAL GVLYGKMFSQ TTICRFESLQ LSFKNMCQLK PFLERWVVEA ENNDNLQELI NR--------
X160 TEE---D-GM TLEEMEEFAK ELKQKRVALG YTQGDIGHAL GILYGKMFSQ TTICRFESLQ LTFKNMCKLK PLLEQWLGEA ENNDNLQEMI HK--------
AmPOU5F3 GDE---D-TP TNEELEQFAK ALKHKRITLG FTQADVGMAL GSLYGKMFSQ TTICRFEALQ LSFKNMCKLK PLLQRWLNEA ENTDNMEELC NM--------
LcPOU5F3 ADE---D-NP TTEELEQFAK ELKHKRITLG FTQADVGLAL GTLYGKMFSQ TTICRFEALQ LSFKNMCKLK PLLQRWLNEA ENNENLQEMC NI--------
AsPOU5F3 EEE---E-NL STEELEQFAK ELKHKRITLG FTQADVGLAL GNLYGKMFSQ TTICRFEALQ LSFKNMCKLK PLLQRWLNEA ENTDNPQDMY KI--------
DrPOU5F3 EEE---E-TL TTEDLEQFAK ELKHKRITLG FTQADVGLAL GNLYGKMFSQ TTICRFEALQ LSFKNMCKLK PLLQRWLNEA ENSENPQDMY KI--------
OlPOU5F3 SEE---E-NL STEELEQFAK ELKHKRITLG FTQADVGLAL GNLYGKMFSQ TTICRFEALQ LSFKNMCKLK PLLQRWLDEA ETSENPQDMY KI--------
TrPOU5F3 SEE---E-IL TTEELEQFAK ELKHKRITLG FTQADVGLAL GNLYGKMFSQ TTICRFEALQ LSFKNMCRLK PLLQKWLNEA ETTENPQDMY KV--------
LoPOU5F3 EDE---E-VI TSQDLEQFSK EFKQKRITMG FTQADVGLAL GHLYGKMFSQ TTICRFEALQ LSYKNLCKLK PLLQSWLAEA EASENPQDLF KV--------
CmPOU5F3 SEE---E-YR TKGKMEEFAK ELKRKRITLG FTQADVGIAL GNLYGKMFSQ TTICRFEALQ LSFKNMCKLK PILERWLNDA QNHDGVHEIC VT--------
CpuPOU5F3 S-E---E-YP TKEKMEQFAK ELKHKRITLG FTQADVGLAL GNLYGKMFSQ TTICRFEALQ LSFKNMCKLK PILQRWLNDA ENNGGLQEIC NM--------
RhPOU5F3 SEE---E-YP TKEKMEQFAK ELKHKRITLG FTQADVGLAL GNLYGKMFSQ TTICRFEALQ LSFKNMCKLK PILQRWLNDA ENNGSLHEIC NV--------
HzPOU5F3 SEE---E-YP TKEKMEQFAK ELKHKRITLG FTQADVGLAL GNLYGKMFSQ TTICRFEALQ LSFKNMCKLK PILQRWLNDA ENNGGLHEIC NV--------
ScPOU5F3 SEE---E-YP TKEKMEQFAK ELKHKRITLG FTQADVGLAL GNLYGKMFSQ TTICRFEALQ LSFKNMCKLK PILQRWLNDA ENNGGLHEIC NV--------
StPOU5F3 SEE---E-YP TKEKMEQFAK ELKHKRITLG FTQADVGLAL GNLYGKMFSQ TTICRFEALQ LSFKNMCKLK PILQRWLNDA ENNGGLQEIC HV--------
LePOU5F3 SEE---E-YP TKEKMEQFAK ELKHKRITLG FTQADVGLAL GNLYGKMFSQ TTICRFEALQ LSFKNMCKLK PILQRWLNDA ESDGGIHEIC NV--------
ArPOU5F3 SEE---E-YP TKEKMEQFAK ELKHKRITLG FTQADVGLAL GNLYGKMFSQ TTICRFEALQ LSFKNMCKLK PILQRWLNDA ESDGGIHEIC NV--------
OkPOU5F3 XXX---X-YP TKEKMEQFAK ELKHKRITLG FTQADVGLAL GNLYGKMFSQ TTICRFEALQ LSFKNMCKLK PILQRWLNDA ESDGGIHEIC NV--------
PmPou5 GEAA--S-DG NIDELARFAK ELKVKRVNLG FTQAEMGLSI GSLCGRVFSQ TTICRFEALQ LSQRNLCKLQ PLFKLWMDDV GVNGGSGGVS SVGGGVGGK-
LjPou5 GEVG--S-DG NIDELARFAK ELKMKRVNLG FTQAEMGISI GSLCGRVFSQ TTICRFEALQ LSQRNLCKLQ PLFKLWMEDV QSNGSISSNG GNGGNGGNGG
EbPou5 GDE---G-TI SSETLAQFAR DLKHKRITLG FTQADVGVAL GSLYGRIFSQ TTICRFEALQ LSYRNMCKLQ PLLERWMIEA ENADNVKEVC HLFLFNSSMD
BfPOU2 GD----E-PS QLEELEQFAK MFKQRRIKLG FTQGDVGLAM GKLYGNDFSQ TTISRFEALN LSFKNMCKLK PLLEKWLQDA DSSMANPGAL GSPHGSI---
BfPOU3.1 XE----E-SP TADELEQFAM VFKQRRIKLG FTQADVGLAL GTIHGNVLSQ TTICRFEALQ LSFKNMCKLK PLLQKWLETA DSNMSPGSLP KDEGLGG---
BfPOU3.2 SDD---D-TP TSDDLEQFAK QFKQRRIKLG FTQADVGLAL GTLYGNVFSQ TTICRFEALQ LSFKNMCKLK PLLQKWLEEA DSSSGSPSSI DK--------
Q-E
R-K
K-T
N-K
F-L

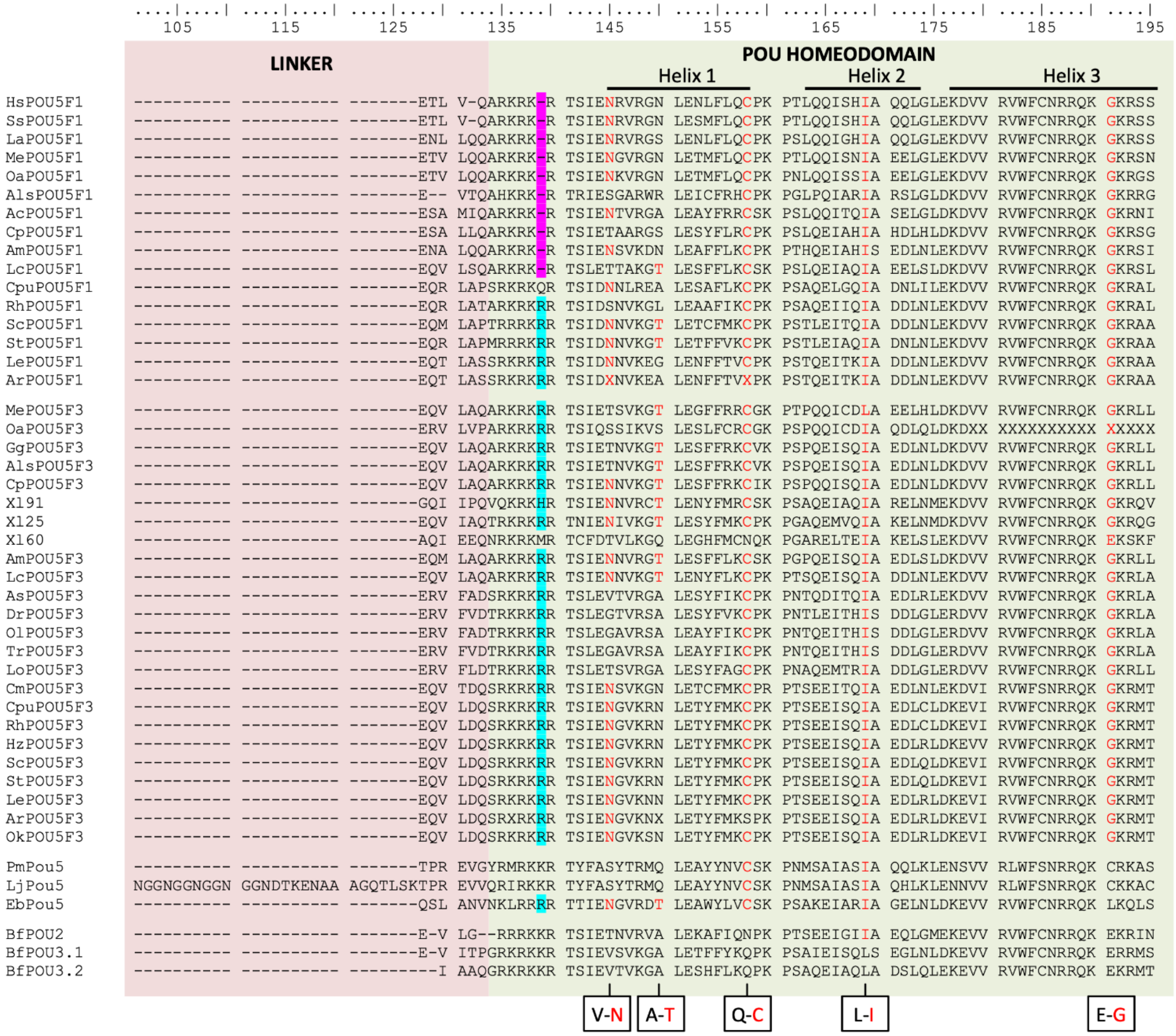
105 115 125 135 145 155 165 175 185 195
LINKER
POU HOMEODOMAIN
Helix 1
Helix 2
Helix 3
HsPOU5F1 ---------- ---------- -------ETL V-QARKRK-R TSIENRVRGN LENLFLQCPK PTLQQISHIA QQLGLEKDVV RVWFCNRRQK GKRSS
SsPOU5F1 ---------- ---------- -------ETL V-QARKRK-R TSIENRVRGN LESMFLQCPK PTLQQISHIA QQLGLEKDVV RVWFCNRRQK GKRSS
LaPOU5F1 ---------- ---------- -------ENL LQQARKRK-R TSIENRVRGS LENLFLQCPK PSLQQIGHIA QQLGLEKDVV RVWFCNRRQK GKRSS
MePOU5F1 ---------- ---------- -------ETV LQQARKRK-R TSIENGVRGN LETMFLQCPK PTLQQISNIA EELGLEKDVV RVWFCNRRQK GKRSN
OaPOU5F1 ---------- ---------- -------ETV LQQARKRK-R TSIENKVRGN LETMFLQCPK PNLQQISSIA EELGLEKDVV RVWFCNRRQK GKRGS
AlsPOU5F1 ---------- ---------- -------E-- VTQAHKRK-R TRIESGARWR LEICFRHCPK PGLPQIARIA RSLGLDKDVV RVWFCNRRQK GKRRG
AcPOU5F1 ---------- ---------- -------ESA MIQARKRK-R TSIENTVRGA LEAYFRRCSK PSLQQITQIA SELGLDKDVV RVWFCNRRQK GKRNI
CpPOU5F1 ---------- ---------- -------ESA LLQARKRK-R TSIETAARGS LESYFLRCPK PSLQEIAHIA HDLHLDKDVV RVWFCNRRQK GKRSG
AmPOU5F1 ---------- ---------- -------ENA LQQARKRK-R TSIENSVKDN LEAFFLKCPK PTHQEIAHIS EDLNLEKDVV RVWFCNRRQK GKRSI
LcPOU5F1 ---------- ---------- -------EQV LSQARKRK-R TSLETTAKGT LESFFLKCSK PSLQEIAQIA EELSLDKDVV RVWFCNRRQK GKRSL
CpuPOU5F1 ---------- ---------- -------EQR LAPSRKRKQR TSIDNNLREA LESAFLKCPK PSAQELGQIA DNLILEKDVV RVWFCNRRQK GKRAL
RhPOU5F1 ---------- ---------- -------EQR LATARKRKRR TSIDSNVKGL LEAAFIKCPK PSAQEIIQIA DDLNLEKDVV RVWFCNRRQK GKRAL
ScPOU5F1 ---------- ---------- -------EQM LAPTRRKKRR TSIDNNVKGT LETCFMKCPK PSTLEITQIA DDLNLEKDVV RVWFCNRRQK GKRAA
StPOU5F1 ---------- ---------- -------EQR LAPMRRRKRR TSIDNNVKGT LETFFVKCPK PSTLEIAQIA DNLNLEKDVV RVWFCNRRQK GKRAA
LePOU5F1 ---------- ---------- -------EQT LASSRKRKRR TSIDNNVKEG LENFFTVCPK PSTQEITKIA DDLNLEKDVV RVWFCNRRQK GKRAA
ArPOU5F1 ---------- ---------- -------EQT LASSRKRKRR TSIDXNVKEA LENFFTVXPK PSTQEITKIA DDLNLEKDVV RVWFCNRRQK GKRAA
MePOU5F3 ---------- ---------- -------EQV LAQARKRKRR TSIETSVKGT LEGFFRRCGK PTPQQICDLA EELHLDKDVV RVWFCNRRQK GKRLL
OaPOU5F3 ---------- ---------- -------ERV LVPARKRKRR TSIQSSIKVS LESLFCRCGK PSPQQICDIA QDLQLDKDXX XXXXXXXXXX XXXXX
GgPOU5F3 ---------- ---------- -------EQV LAQARKRKRR TSIETNVKGT LESFFRKCVK PSPQEISQIA EDLNLDKDVV RVWFCNRRQK GKRLL
AlsPOU5F3 ---------- ---------- -------EQV LAQARKRKRR TSIETNVKGT LESFFRKCVK PSPQEISQIA EDLNLDKDVV RVWFCNRRQK GKRLL
CpPOU5F3 ---------- ---------- -------EQV LAQARKRKRR TSIENNVKGT LESFFRKCIK PSPQQISQIA EDLNLDKDVV RVWFCNRRQK GKRLL
X191 ---------- ---------- -------GQI IPQVQKRKHR TSIENNVRCT LENYFMRCSK PSAQEIAQIA RELNMEKDVV RVWFCNRRQK GKRQV
X125 ---------- ---------- -------EQV IAQTRKRKRR TNIENIVKGT LESYFMKCPK PGAQEMVQIA KELNMDKDVV RVWFCNRRQK GKRQG
X160 ---------- ---------- -------AQI EEQNRKRKMR TCFDTVLKGQ LEGHFMCNQK PGARELTEIA KELSLEKDVV RVWFCNRRQK EKSKF
AmPOU5F3 ---------- ---------- -------EQM LAQARKRKRR TSIENNVRGT LESFFLKCSK PGPQEISQIA EDLSLDKDVV RVWFCNRRQK GKRLL
LcPOU5F3 ---------- ---------- -------EQV LAQARKRKRR TSIENNVKGT LENYFLKCPK PTSQEISQIA DDLNLEKDVV RVWFCNRRQK GKRLA
AsPOU5F3 ---------- ---------- -------ERV FADSRKRKRR TSLEVTVRGA LESYFIKCPK PNTQDITQIA EDLRLEKDVV RVWFCNRRQK GKRLA
DrPOU5F3 ---------- ---------- -------ERV FVDTRKRKRR TSLEGTVRSA LESYFVKCPK PNTLEITHIS DDLGLERDVV RVWFCNRRQK GKRLA
OlPOU5F3 ---------- ---------- -------ERV FADTRKRKRR TSLEGAVRSA LEAYFIKCPK PNTQEITHIS DDLGLERDVV RVWFCNRRQK GKRLA
TrPOU5F3 ---------- ---------- -------ERV FVDTRKRKRR TSLEGAVRSA LEAYFIKCPK PNTQEITHIS DDLGLERDVV RVWFCNRRQK GKRLA
LoPOU5F3 ---------- ---------- -------ERV FLDTRKRKRR TSLETSVRGA LESYFAGCPK PNAQEMTRIA DDLGLERDVV RVWFCNRRQK GKRLL
CmPOU5F3 ---------- ---------- -------EQV TDQSRKRKRR TSIENSVKGN LETCFMKCPR PTSEEITQIA EDLNLEKDVI RVWFSNRRQK GKRMT
CpuPOU5F3 ---------- ---------- -------EQV LDQSRKRKRR TSIENGVKRN LETYFMKCPK PTSEEISQIA EDLCLDKEVI RVWFCNRRQK GKRMT
RhPOU5F3 ---------- ---------- -------EQV LDQSRKRKRR TSIENGVKRN LETYFMKCPK PTSEEISQIA EDLCLDKEVI RVWFCNRRQK GKRMT
HzPOU5F3 ---------- ---------- -------EQV LDQSRKRKRR TSIENGVKRN LETYFMKCPK PTSEEISQIA EDLRLDKEVV RVWFCNRRQK GKRMT
ScPOU5F3 ---------- ---------- -------EQV LDQSRKRKRR TSIENGVKRN LETYFMKCPK PTSEEISQIA EDLQLDKEVI RVWFCNRRQK GKRMT
StPOU5F3 ---------- ---------- -------EQV LDQSRKRKRR TSIENGVKRN LETYFMKCPK PTSEEISQIA EDLQLDKEVI RVWFCNRRQK GKRMT
LePOU5F3 ---------- ---------- -------EQV LDQSRKRKRR TSIENGVKNN LETYFMKCPK PTSEEISQIA EDLRLDKEVI RVWFCNRRQK GKRMT
ArPOU5F3 ---------- ---------- -------EQV LDQSRXRKRR TSIENGVKNX LETYFMKSPK PTSEEISQIA EDLRLDKEVI RVWFCNRRQK GKRMT
OkPOU5F3 ---------- ---------- -------EQV LDQSRKRKRR TSIENGVKSN LETYFMKCPK PTSEEISQIA EDLRLDKEVI RVWFCNRRQK GKRMT
PmPou5 ---------- ---------- -------TPR EVGYRMRKKR TYFASYTRMQ LEAYYNVCSK PNMSAIASIA QQLKLENSVV RLWFSNRRQK CRKAS
LjPou5 NGGNGGNGGN GGNDTKENAA AGQTLSKTPR EVVQRIRKKR TYFASYTRMQ LEAYYNVCSK PNMSAIASIA QHLKLENNVV RLWFSNRRQK CKKAC
EbPou5 ---------- ---------- -------QSL ANVNKLRRRR TTIENGVRDT LEAWYLVCSK PSAKEIARIA GELNLDKEVV RVWFCNRRQK LKQLS
BfPOU2 ---------- ---------- -------E-V LG--RRRKKR TSIETNVRVA LEKAFIQNPK PTSEEIGIIA EQLGMEKEVV RVWFCNRRQK EKRIN
BfPOU3.1 ---------- ---------- -------E-V ITPGRKRKKR TSIEVSVKGA LETFFYKQPK PSAIEISQLS EGLNLDKEVV RVWFCNRRQK ERRMS
BfPOU3.2 ---------- ---------- ---------I AAQGRKRKKR TSIEVTVKGA LESHFLKQPK PSAQEIAQLA DSLQLEKEVV RVWFCNRRQK EKRMT
V-N
A-T
Q-C
L-I
E-G

**<u>Supplementary information related to Fig. 1b-c.</u> Detailed syntenies between *POU5* loci in vertebrates and related phylogenetic analyses.**

**(e-g) detailed conserved syntenies between gnathostome *POU5F1* (e), *POU5F3* (f) and cyclostome (g) POU5 loci.** Genes identified in the genomes of the species analysed are depicted by coloured arrows. Black crosses over empty arrows indicate that the corresponding gene was not identified in the genome version analysed (see Suppl. Table 1). Continuous black lines connect genes found in synteny. The loci are identified by chromosome or scaffold names. Accession numbers are provided in Suppl. Table 1.

**e Gnathostome *POU5F1***

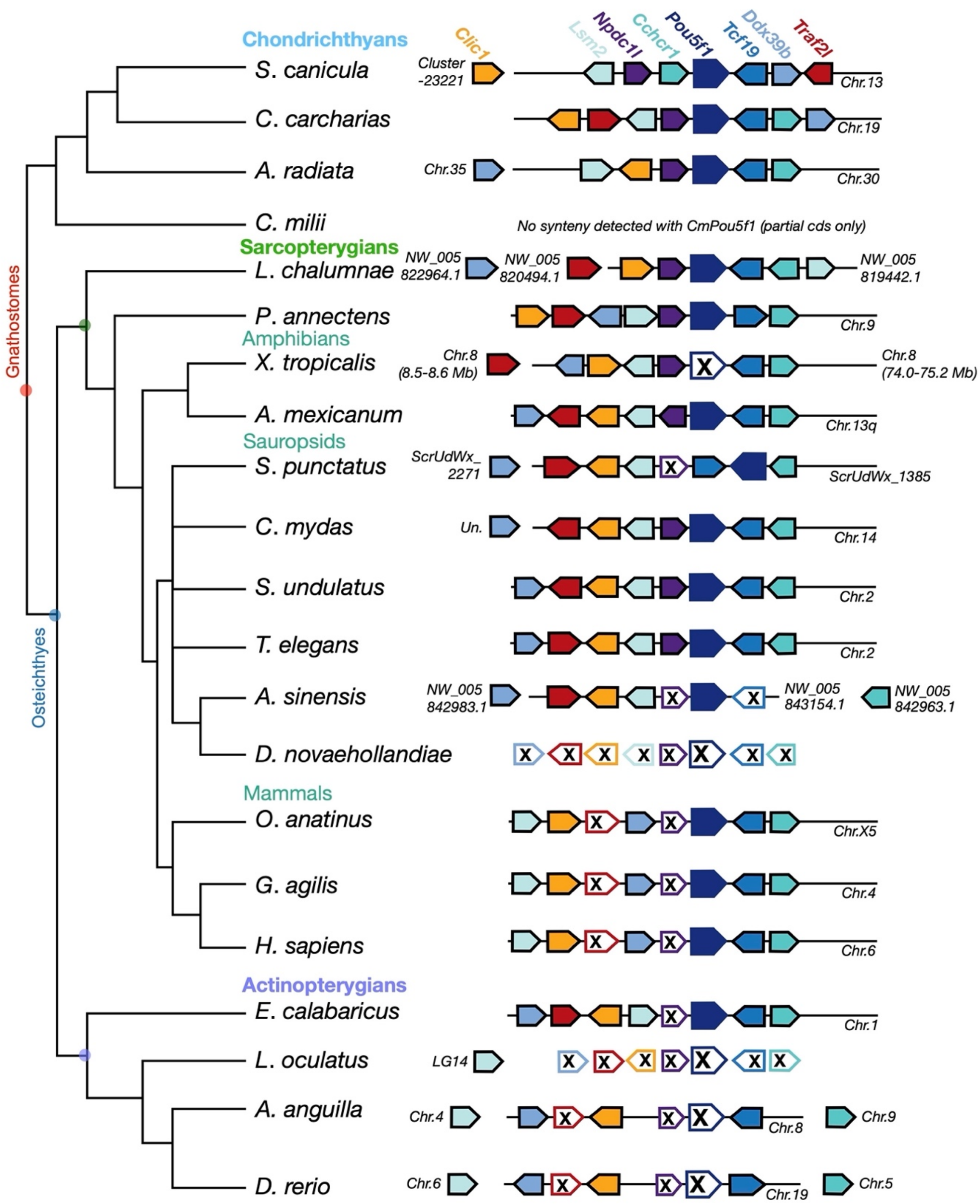

**f** Gnathostome *POU5F3*

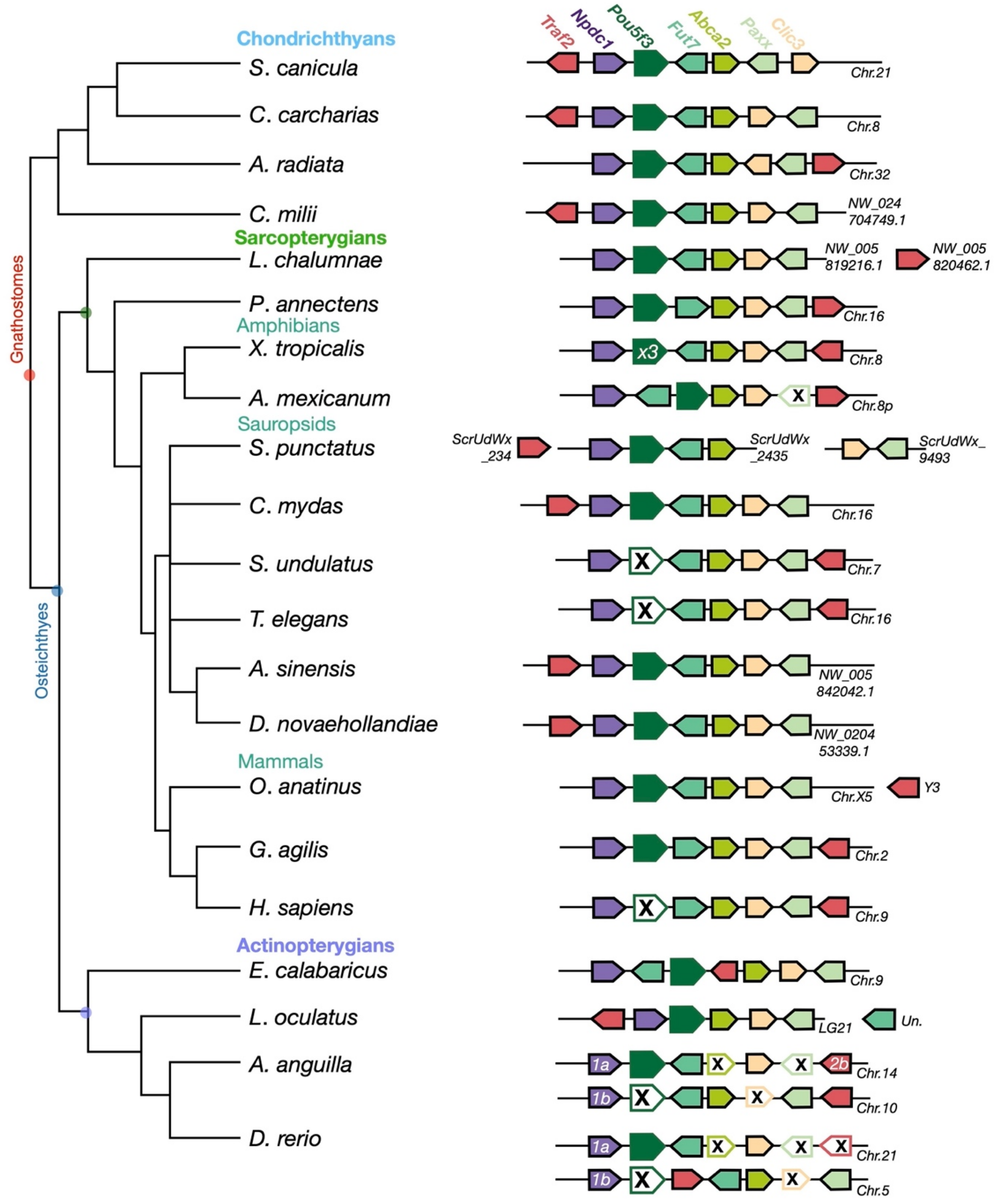

**g** **Cyclostome *POU5***

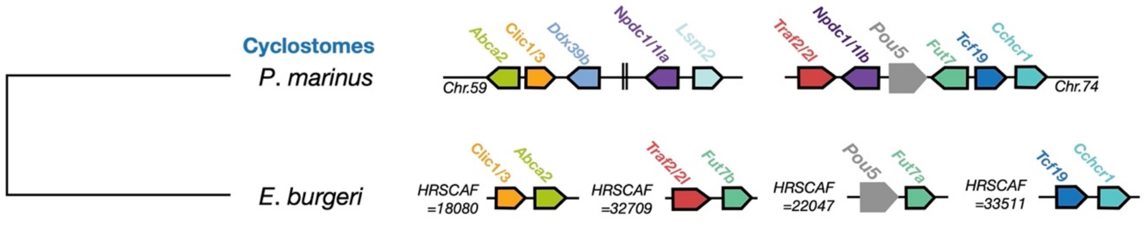

**(h-k) Phylogenetic trees showing the relationships between gnathostomes POU5F1 and POU5F3 proteins (h) and between paralogous genes used in synteny analyses: TRAF2/2L (i), CLIC1/3 (j), NPDC1/NPC1L (k).** **(h)** is unrooted, **(j-k)** were rooted using amphioxus or hemichordate sequences. The trees were calculated using IQTree with the following parameters: NNI moves, LG+R4, aLRT branch support. The numbers at each node indicate posterior probabilities of group occurrence (in %). In (j-k), genes exhibiting conserved linkages with cyclostome *POU5* genes, gnathostome *POU5F1* and *POU5F3* are shaded in grey, blue and green respectively. Species abbreviations: Aa, *Anguilla anguilla*; As, *Acipenser sinensis*; Ac, *Anolis carolinensis*; Als, *Alligator sinensis*; Ar, *Amblyraja radiata*; Am, *Ambystoma mexicanum*; Amx, *Astyanax mexicanum*; Ao, *Amphiprion ocellaris*; Bb, *Branchiostoma belcheri*; Bf, *Branchiostoma floridae*; Cl, *Columbia livia*; Cm, *Callorhinchus milii*; Cp, *Chrysemys picta*; Cm, *Chelonia mydas*; Cpu, *Chiloscyllium punctatum*; Dn, *Dromaius novaehollandiae*; Dr, *Danio rerio*; Eb, *Eptatretus burgeri*; Gag, *Gracilinanus agilis*; Gg, *Gallus gallus*; Gm, *Gadus morhua*; Hs, *Homo sapiens*; Hz, *Heterodontus zebra*; La, *Loxodonta africana*; Lc, *Latimeria chalumnae*; Le, *Leucoraja erinacea*; Lj: *Lethenteron japonicum*; Lo, *Lepisosteus oculatus*; Md, *Monodelphis domestica*; Me, *Macropus eugenii*; Oa, *Ornithorhynchus anatinus*; Ok, *Okamejei kenojei*; Ol, *Oryzias latipes*; On, *Oreochromis niloticus*; Pa, *Protopterus annectens*; Pm, *Petromyzon marinus*; Ps, *Pelodiscus sinensis*; Rh/Rt, *Rhincodon typus*; Sc, *Scyliorhinus canicula*; Sk, *Saccoglossus kowalevskii*; Ss, *Sus scrofa*; Ssl, *Salmo salar*; St, *Scyliorhinus torazame*; Su, *Sceloporus undulatus*; Te, *Thamnophis elegans*; Tr, *Takifugu rubripes*; Xl, *Xenopus laevis*; Xt, *Xenopus tropicalis*.

h

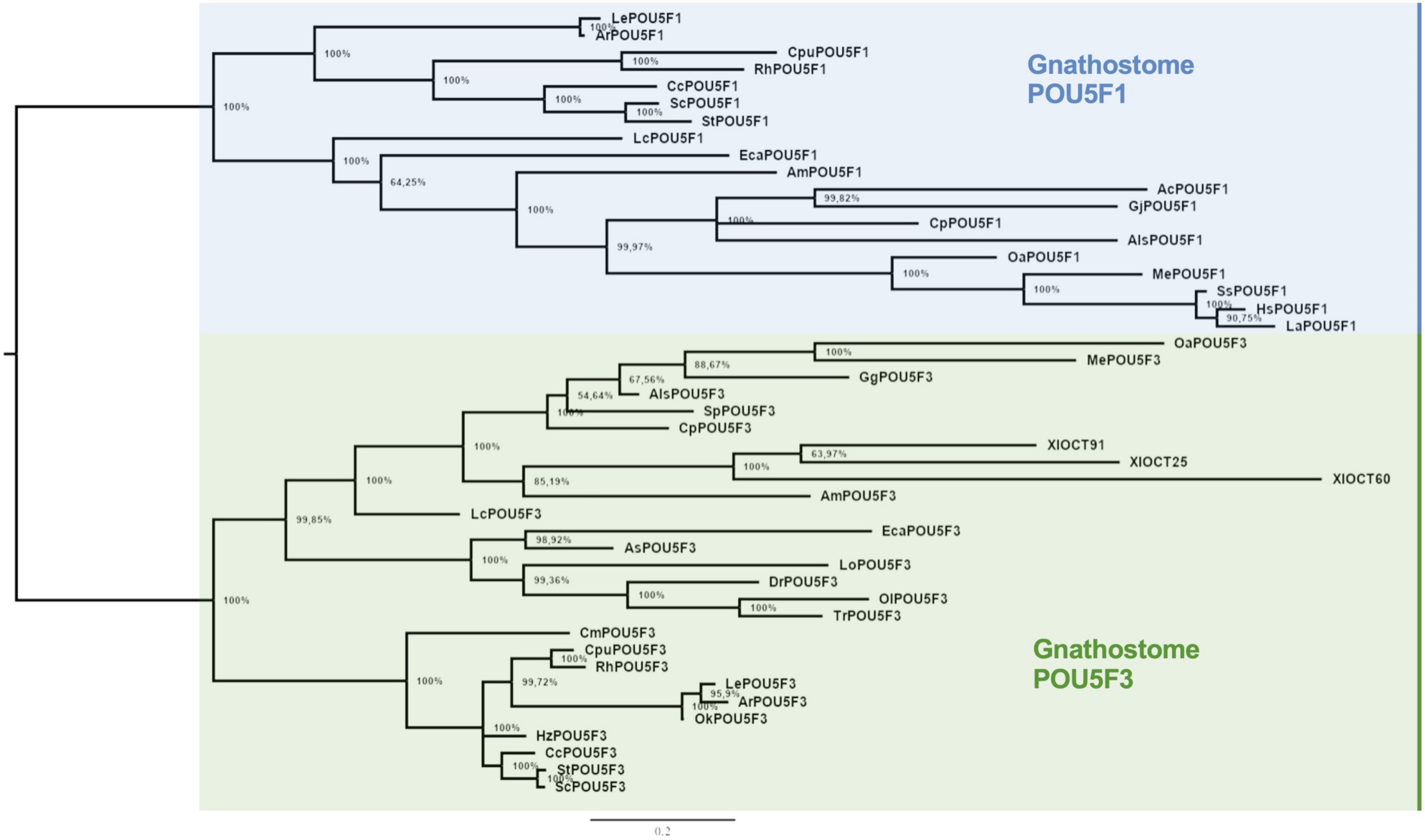

Gnathostome POU5F1
LePOU5F1
ArPOU5F1
CpuPOU5F1
RhPOU5F1
CcPOU5F1
ScPOU5F1
StPOU5F1
LcPOU5F1
EcaPOU5F1
AmPOU5F1
AcPOU5F1
GjPOU5F1
CpPOU5F1
AlsPOU5F1
OaPOU5F1
MePOU5F1
SsPOU5F1
HsPOU5F1
LaPOU5F1
Gnathostome POU5F3
OaPOU5F3
MePOU5F3
GgPOU5F3
AlsPOU5F3
SpPOU5F3
CpPOU5F3
XlOCT91
XlOCT25
XlOCT60
AmPOU5F3
LcPOU5F3
EcaPOU5F3
AsPOU5F3
LoPOU5F3
DrPOU5F3
OlPOU5F3
TrPOU5F3
CmPOU5F3
CpuPOU5F3
RhPOU5F3
LePOU5F3
ArPOU5F3
OkPOU5F3
HzPOU5F3
CcPOU5F3
StPOU5F3
ScPOU5F3
0.2

i

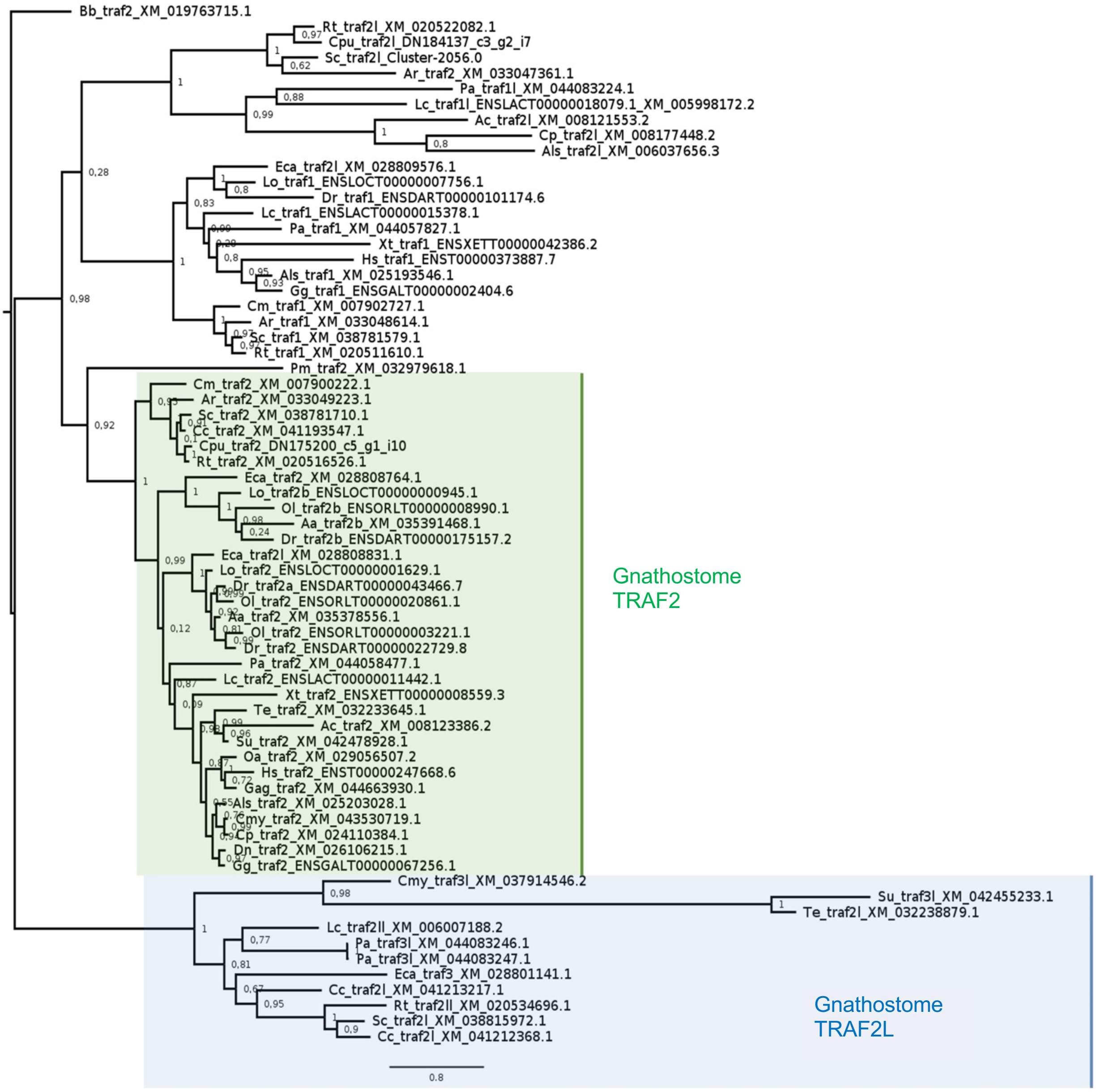

Bb_traf2_XM_019763715.1
Rt_traf2l_XM_020522082.1
Cpu_traf2l_DN184137_c3_g2_i7
Sc_traf2l_Cluster-2056.0
Ar_traf2_XM_033047361.1
Pa_traf1l_XM_044083224.1
Lc_traf1l_ENSLACT00000018079.1_XM_005998172.2
Ac_traf2l_XM_008121553.2
Cp_traf2l_XM_008177448.2
Als_traf2l_XM_006037656.3
Eca_traf2l_XM_028809576.1
Lo_traf1_ENSLOCT00000007756.1
Dr_traf1_ENSDART00000101174.6
Lc_traf1_ENSLACT00000015378.1
Pa_traf1_XM_044057827.1
Xt_traf1_ENSXETT00000042386.2
Hs_traf1_ENST00000373887.7
Als_traf1_XM_025193546.1
Gg_traf1_ENSGALT00000002404.6
Cm_traf1_XM_007902727.1
Ar_traf1_XM_033048614.1
Sc_traf1_XM_038781579.1
Rt_traf1_XM_020511610.1
Pm_traf2_XM_032979618.1
Cm_traf2_XM_007900222.1
Ar_traf2_XM_033049223.1
Sc_traf2_XM_038781710.1
Cc_traf2_XM_041193547.1
Cpu_traf2_DN175200_c5_g1_i10
Rt_traf2_XM_020516526.1
Eca_traf2_XM_028808764.1
Lo_traf2b_ENSLOCT00000000945.1
Ol_traf2b_ENSORLT00000008990.1
Aa_traf2b_XM_035391468.1
Dr_traf2b_ENSDART00000175157.2
Eca_traf2l_XM_028808831.1
Lo_traf2_ENSLOCT00000001629.1
Dr_traf2a_ENSDART00000043466.7
Ol_traf2_ENSORLT00000020861.1
Aa_traf2_XM_035378556.1
Ol_traf2_ENSORLT00000003221.1
Dr_traf2_ENSDART00000022729.8
Pa_traf2_XM_044058477.1
Lc_traf2_ENSLACT00000011442.1
Xt_traf2_ENSXETT00000008559.3
Te_traf2_XM_032233645.1
Ac_traf2_XM_008123386.2
Su_traf2_XM_042478928.1
Oa_traf2_XM_029056507.2
Hs_traf2_ENST00000247668.6
Gag_traf2_XM_044663930.1
Als_traf2_XM_025203028.1
Cmy_traf2_XM_043530719.1
Cp_traf2_XM_024110384.1
Dn_traf2_XM_026106215.1
Gg_traf2_ENSGALT00000067256.1
Cmy_traf3l_XM_037914546.2
Su_traf3l_XM_042455233.1
Te_traf2l_XM_032238879.1
Lc_traf2ll_XM_006007188.2
Pa_traf3l_XM_044083246.1
Pa_traf3l_XM_044083247.1
Eca_traf3_XM_028801141.1
Cc_traf2l_XM_041213217.1
Rt_traf2ll_XM_020534696.1
Sc_traf2l_XM_038815972.1
Cc_traf2l_XM_041212368.1
Gnathostome TRAF2
Gnathostome TRAF2L
0,8

j

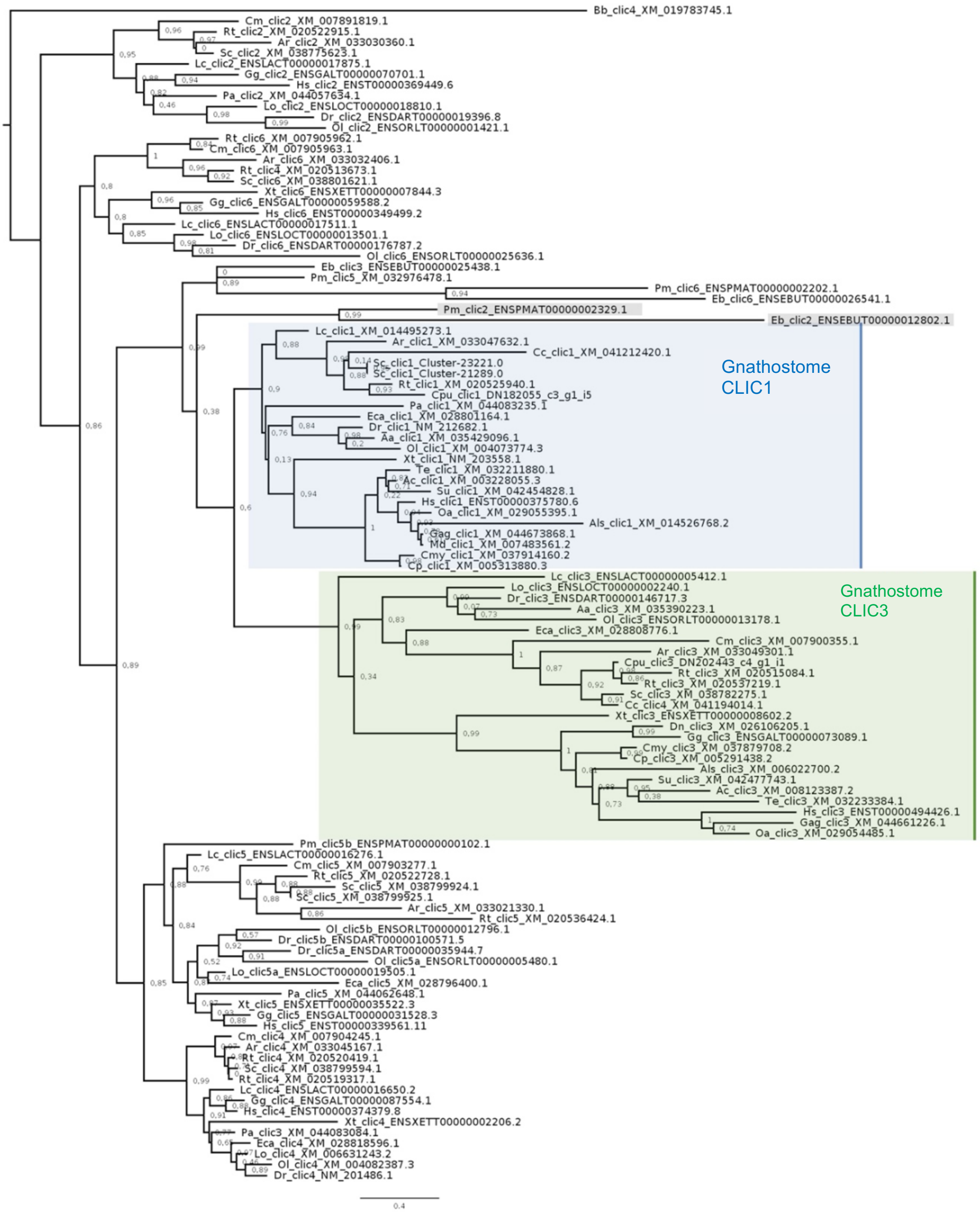

Bb_clic4_XM_019783745.1
Cm_clic2_XM_007891819.1
Rt_clic2_XM_020522915.1
Ar_clic2_XM_033030360.1
Sc_clic2_XM_038775623.1
Lc_clic2_ENSLACT00000017875.1
Gg_clic2_ENSGALT00000070701.1
Hs_clic2_ENST00000369449.6
Pa_clic2_XM_044057634.1
Lo_clic2_ENSLOCT00000018810.1
Dr_clic2_ENSDART00000019396.8
Ol_clic2_ENSORLT00000001421.1
Rt_clic6_XM_007905962.1
Cm_clic6_XM_007905963.1
Ar_clic6_XM_033032406.1
Rt_clic4_XM_020513673.1
Sc_clic6_XM_038801621.1
Xt_clic6_ENSXETT00000007844.3
Gg_clic6_ENSGALT00000059588.2
Hs_clic6_ENST00000349499.2
Lc_clic6_ENSLACT00000017511.1
Lo_clic6_ENSLOCT00000013501.1
Dr_clic6_ENSDART00000176787.2
Ol_clic6_ENSORLT00000025636.1
Eb_clic3_ENSEBUT00000025438.1
Pm_clic5_XM_032976478.1
Pm_clic6_ENSPMAT00000002202.1
Eb_clic6_ENSEBUT00000026541.1
Pm_clic2_ENSPMAT00000002329.1
Eb_clic2_ENSEBUT00000012802.1
Lc_clic1_XM_014495273.1
Ar_clic1_XM_033047632.1
Cc_clic1_XM_041212420.1
Sc_clic1_Cluster-23221.0
Sc_clic1_Cluster-21289.0
Rt_clic1_XM_020525940.1
Cpu_clic1_DN182055_c3_g1_i5
Pa_clic1_XM_044083235.1
Eca_clic1_XM_028801164.1
Dr_clic1_NM_212682.1
Aa_clic1_XM_035429096.1
Ol_clic1_XM_004073774.3
Xt_clic1_NM_203558.1
Te_clic1_XM_032211880.1
Ac_clic1_XM_003228055.3
Su_clic1_XM_042454828.1
Hs_clic1_ENST00000375780.6
Oa_clic1_XM_029055395.1
Als_clic1_XM_014526768.2
Gag_clic1_XM_044673868.1
Md_clic1_XM_007483561.2
Cmy_clic1_XM_037914160.2
Cp_clic1_XM_005313880.3
Gnathostome CLIC1
Lc_clic3_ENSLACT00000005412.1
Lo_clic3_ENSLOCT00000002240.1
Dr_clic3_ENSDART00000146717.3
Aa_clic3_XM_035390223.1
Ol_clic3_ENSORLT00000013178.1
Eca_clic3_XM_028808776.1
Cm_clic3_XM_007900355.1
Ar_clic3_XM_033049301.1
Cpu_clic3_DN202443_c4_g1_i1
Rt_clic3_XM_020515084.1
Rt_clic3_XM_020537219.1
Sc_clic3_XM_038782275.1
Cc_clic4_XM_041194014.1
Xt_clic3_ENSXETT00000008602.2
Dn_clic3_XM_026106205.1
Gg_clic3_ENSGALT00000073089.1
Cmy_clic3_XM_037879708.2
Cp_clic3_XM_005291438.2
Als_clic3_XM_006022700.2
Su_clic3_XM_042477743.1
Ac_clic3_XM_008123387.2
Te_clic3_XM_032233384.1
Hs_clic3_ENST00000494426.1
Gag_clic3_XM_044661226.1
Oa_clic3_XM_029054485.1
Gnathostome CLIC3
Pm_clic5b_ENSPMAT00000000102.1
Lc_clic5_ENSLACT00000016276.1
Cm_clic5_XM_007903277.1
Rt_clic5_XM_020522728.1
Sc_clic5_XM_038799924.1
Sc_clic5_XM_038799925.1
Ar_clic5_XM_033021330.1
Rt_clic5_XM_020536424.1
Ol_clic5b_ENSORLT00000012796.1
Dr_clic5b_ENSDART00000100571.5
Dr_clic5a_ENSDART00000035944.7
Ol_clic5a_ENSORLT00000005480.1
Lo_clic5a_ENSLOCT00000019505.1
Eca_clic5_XM_028796400.1
Pa_clic5_XM_044062648.1
Xt_clic5_ENSXETT00000035522.3
Gg_clic5_ENSGALT00000031528.3
Hs_clic5_ENST00000339561.11
Cm_clic4_XM_007904245.1
Ar_clic4_XM_033045167.1
Rt_clic4_XM_020520419.1
Sc_clic4_XM_038799594.1
Rt_clic4_XM_020519317.1
Lc_clic4_ENSLACT00000016650.2
Gg_clic4_ENSGALT00000087554.1
Hs_clic4_ENST00000374379.8
Xt_clic4_ENSXETT00000002206.2
Pa_clic3_XM_044083084.1
Eca_clic4_XM_028818596.1
Lo_clic4_XM_006631243.2
Ol_clic4_XM_004082387.3
Dr_clic4_NM_201486.1
0.4

**k**

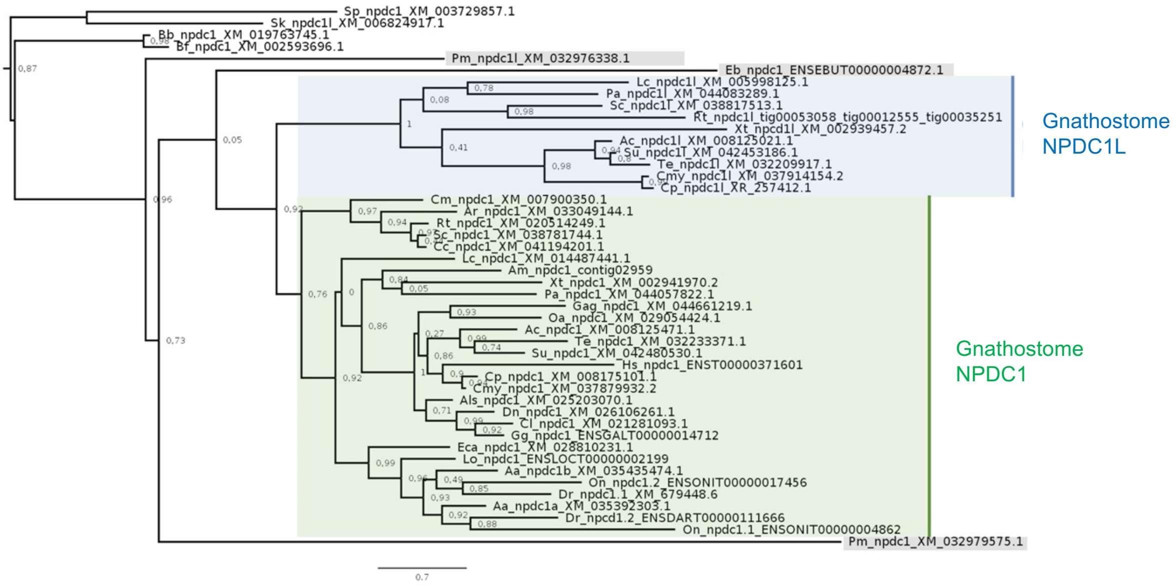
Sp_npdc1_XM_003729857.1
Sk_npdc1l_XM_006824917.1
Bb_npdc1_XM_019763745.1
Bf_npdc1_XM_002593696.1
Pm_npdc1l_XM_032976338.1
Eb_npdc1_ENSEBUT00000004872.1
Lc_npdc1l_XM_005998125.1
Pa_npdc1l_XM_044083289.1
Sc_npdc1l_XM_038817513.1
Rt_npdc1l_tig00053058_tig00012555_tig00035251
Xt_npcd1l_XM_002939457.2
Ac_npdc1l_XM_008125021.1
Su_npdc1l_XM_042453186.1
Te_npdc1l_XM_032209917.1
Cmy_npdc1l_XM_037914154.2
Cp_npdc1l_XR_257412.1
Gnathostome NPDC1L
Cm_npdc1_XM_007900350.1
Ar_npdc1_XM_033049144.1
Rt_npdc1_XM_020514249.1
Sc_npdc1_XM_038781744.1
Cc_npdc1_XM_041194201.1
Lc_npdc1_XM_014487441.1
Am_npdc1_contig02959
Xt_npdc1_XM_002941970.2
Pa_npdc1_XM_044057822.1
Gag_npdc1_XM_044661219.1
Oa_npdc1_XM_029054424.1
Ac_npdc1_XM_008125471.1
Te_npdc1_XM_032233371.1
Su_npdc1_XM_042480530.1
Hs_npdc1_ENST00000371601
Cp_npdc1_XM_008175101.1
Cmy_npdc1_XM_037879932.2
Als_npdc1_XM_025203070.1
Dn_npdc1_XM_026106261.1
Cl_npdc1_XM_021281093.1
Gg_npdc1_ENSGALT00000014712
Eca_npdc1_XM_028810231.1
Lo_npdc1_ENSLOCT00000002199
Aa_npdc1b_XM_035435474.1
On_npdc1.2_ENSONIT00000017456
Dr_npdc1.1_XM_679448.6
Aa_npdc1a_XM_035392303.1
Dr_npcd1.2_ENSDART00000111666
On_npdc1.1_ENSONIT00000004862
Gnathostome NPDC1
Pm_npdc1_XM_032979575.1
0.7

**<u>Related to Fig. 1d.</u> Bayesian analyses of the evolution of mammalian POU5F1 proteins and actinopterygian POU5F3 proteins. (l) Parameters for Bayesian analyses. (m-n) Trees showing evolutionary rates of mammalian POU5F1 proteins (m) and actinopterygian POU5F3 proteins (n).** Analyses were conducted on alignments of POU domain, linker, homeodomain and C-terminal sequences, imposing species phylogenies after Meredith et al. 2011 for mammals and Betancur et al. 2013 for teleosts. Evolutionary rates were calculated using the BEAST software and are represented in black, green to red from low, moderate to high. The turtle *C. picta* was used as outgroup for mammals. The whale shark *R. typus* and coelacanth *L. chalumnae* were used as outgroups for actinopterygians. Asterisks show branches in which accelerations of evolutionary rates have taken place. The scale bar corresponds to the average number of amino acid changes per site. Species name abbreviations: As, *Acipenser sinensis*; Aml, *Ailuropoda melanoleuca*; Amx, *Astyanax mexicanum*; Ao, *Amphiprion ocellaris*; Ba, *Balanoptera acutorostrata*; Bt: *Bos taurus*; Cb, *Clarias batrachus*; Cg, *Cricetulus griseus*; Clf, *Canis lupus familiaris*; Cl, *Chinchilla lanigera*; Cp, *Chrysemys picta*; Cpo, *Cavia porcellus*; Css, *Ceratotherium simum simum*; Dn, *Dasypus novemcinctus*; Dr, *Danio rerio*; Ec, *Equus caballus*; Ee, *Erinaceus europaeus*; Et, *Echinops telfairi*; Fc, *Felis catus*; Gm, *Gadus morhua*; Ga, *Gasterosteus aculeatus*; Hg, *Heterocephalus glaber*; Hs, *Homo sapiens*; It, *Ictidomys tridecemlineatus*; Jj, *Jaculus jaculus*; La, *Loxodonta africana*; Lc, *Latimeria chalumnae*; Lo, *Lepisosteus oculatus*; Me, *Macropus eugenii*; Ml, *Myotis lucifugus*; Mm, *Mus musculus*; Mz, *Maylandia zebra*; Na, *Neophocaena asiaorientalis*; Nb, *Neolamprologus brichardi*; Oa, *Ornithorhynchus anatinus*; Oaa, *Orycteropus afer afer*; Oc, *Oryctolagus cuniculus*; Og, *Otolemur garnettii*; Ol, *Oryzias latipes*; On, *Oreochromis niloticus*; Op, *Ochotona princeps*; Or, *Odobenus rosmarus divergens*; Pa, *Plecoglossus altivelis*; Pb, *Pantodon buchholzi*; Pc, *Phascolarctos cinereus*; Pf, *Perca fluviatilis*; Ph, *Pangasianodon hypophthalmus*; Pmb, *Peromyscus maniculatus bairdii*; Pv, *Pteropus vampyrus*; Rh/Rt, *Rhincodon typus*; Rn, *Rattus norvegicus*; Sa, *Sorex araneus*; Sd, *Seriola dumerili*; Ss, *Sus scrofa*; Ssl, *Salmo salar*; Tb, *Tupaia belangeri*; Tr, *Takifugu rubripes*; Vp, *Vicugna pacos*.

### (l) Parameters used for the estimation POU5 protein evolutionary rates

For the estimate of protein evolutionary rates, XML files were generated from input protein sequences using BEAUTi (v1.10.3) and the parameters indicated below (1-6). Species phylogeny with a monophyly of gnathostome POUF1 and POU5F3 groups was imposed as shown in Figures, except for the position of cyclostomes (Fig. 1d), which was left unconstrained. They were subjected to further process using BEAST. A plateau in the results was observed starting from 1 million iterations and we ran 9 million additional iterations for a total of 10 million iterations to ensure that convergence was reached. We used online platform CIPRES SCIENCE GATEWAY (https://www.phylo.org) for BEAST analysis (Miller et al., 2010). Two outputs from BEAST analysis (.log and .tree file) were analyzed for statistical outcomes of the phylogenetic tree. We constructed and visualized phylogenetic trees using TreeAnnotator v.1.10.3 and FigTree (v.1.4.2). The rate of protein evolution was visualized as color gradient in FigTree.

1) Substitution Model: JTT (Jones et al., 1992)
2) Site heterogeneity Model: Gamma
3) Number of Gamma Categories: 4
4) Clock Model: Lognormal relaxed clock (Uncorrelated)
5) Tree Prior shared by all tree models: Coalescent Model (Fixed)
6) MCMC: Length of chain:10,000,000, Echo state to screen every: 10,000, Log parameters every: 10,000

**m**

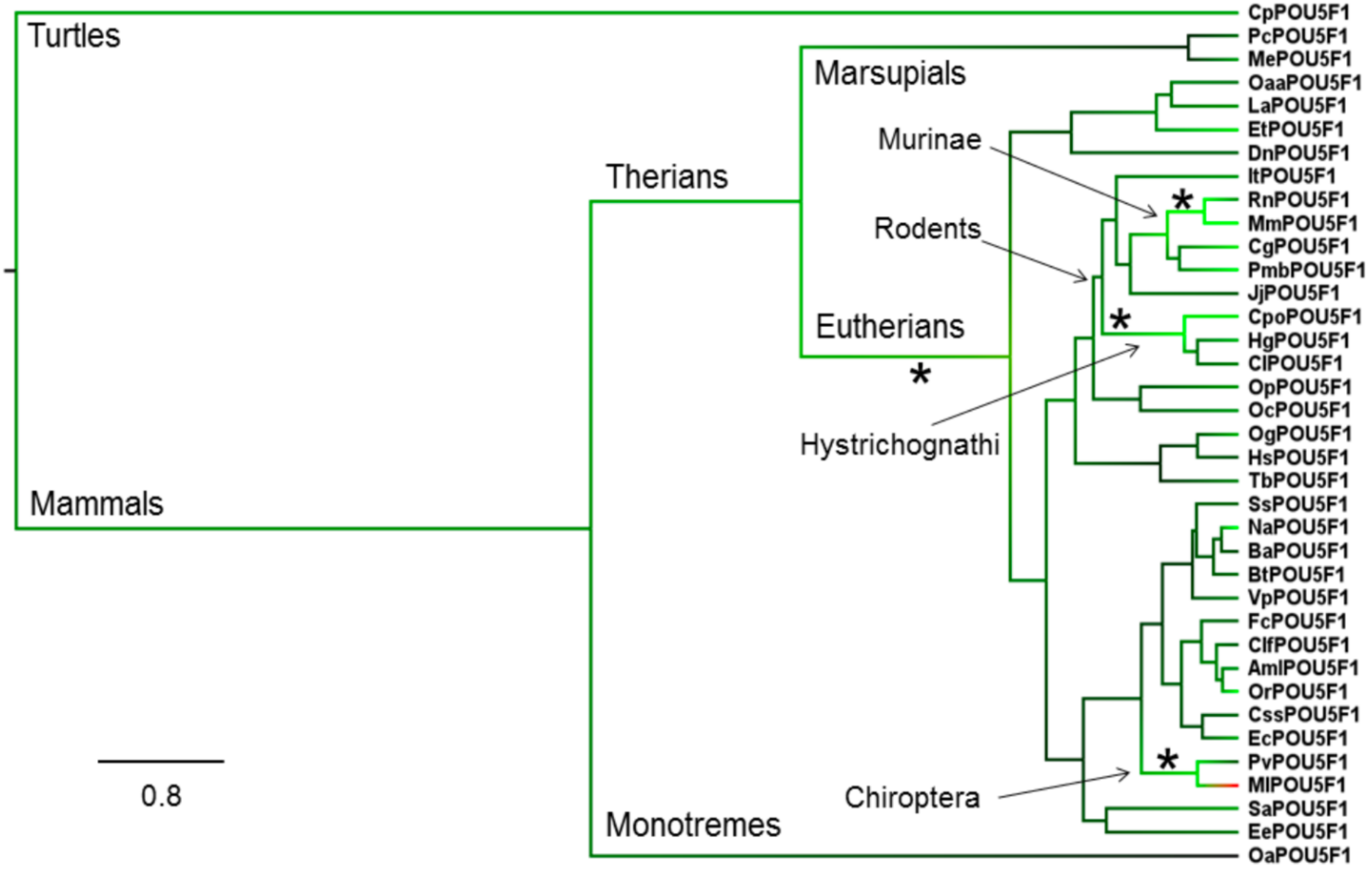
Turtles
Mammals
Therians
Marsupials
Murinae
Rodents
Eutherians
Hystrichognathi
Chiroptera
Monotremes
0.8
*
*
*
*
CpPOU5F1
PcPOU5F1
MePOU5F1
OaaPOU5F1
LaPOU5F1
EtPOU5F1
DnPOU5F1
ItPOU5F1
RnPOU5F1
MmPOU5F1
CgPOU5F1
PmbPOU5F1
JjPOU5F1
CpoPOU5F1
HgPOU5F1
ClPOU5F1
OpPOU5F1
OcPOU5F1
OgPOU5F1
HsPOU5F1
TbPOU5F1
SsPOU5F1
NaPOU5F1
BaPOU5F1
BtPOU5F1
VpPOU5F1
FcPOU5F1
ClfPOU5F1
AmlPOU5F1
OrPOU5F1
CssPOU5F1
EcPOU5F1
PvPOU5F1
MlPOU5F1
SaPOU5F1
EePOU5F1
OaPOU5F1

n

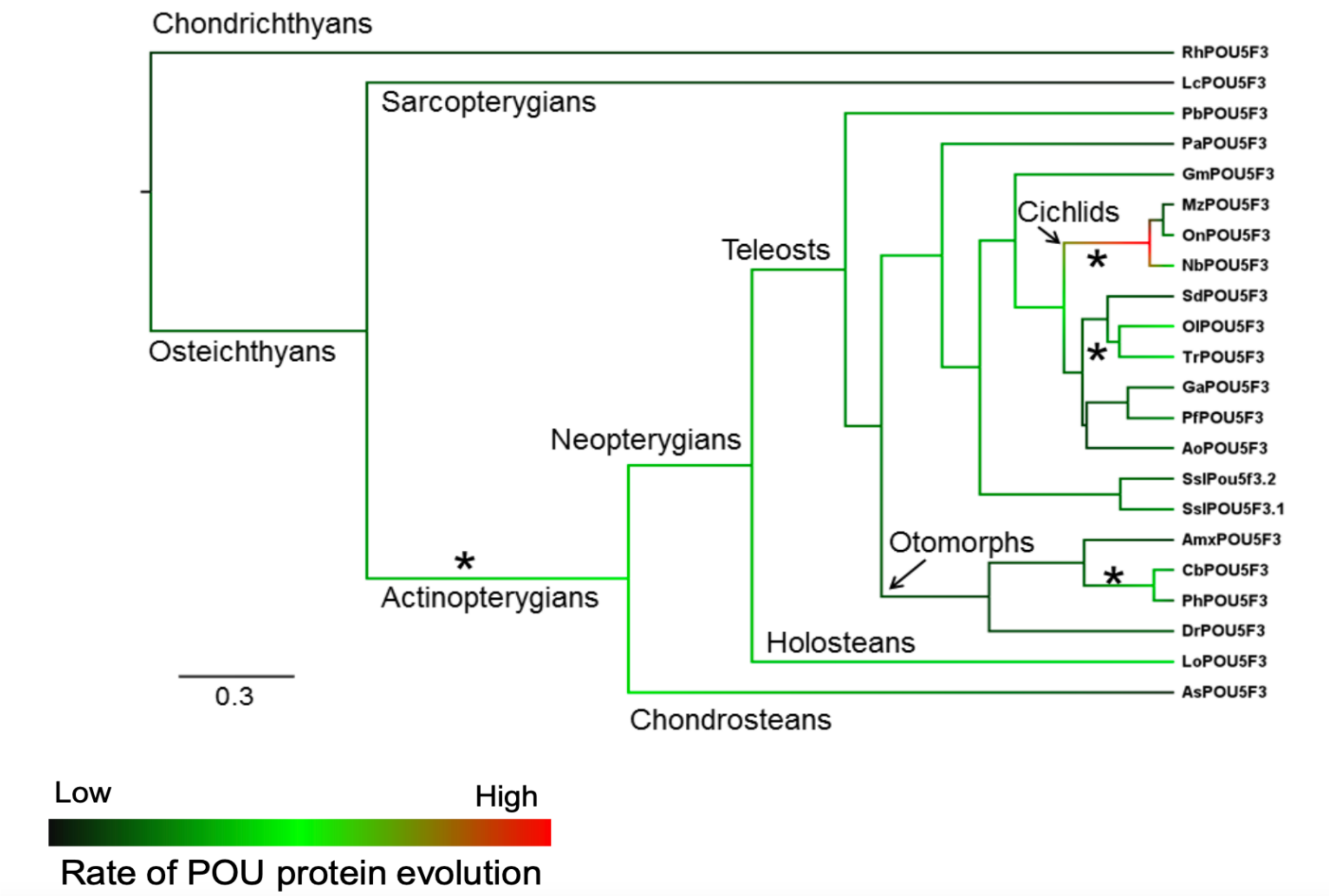

Chondrichthyans
Sarcopterygians
Osteichthyans
Teleosts
Cichlids
Neopterygians
Otomorphs
Actinopterygians
Holosteans
Chondrosteans
0.3
RhPOU5F3
LcPOU5F3
PbPOU5F3
PaPOU5F3
GmPOU5F3
MzPOU5F3
OnPOU5F3
NbPOU5F3
SdPOU5F3
OlPOU5F3
TrPOU5F3
GaPOU5F3
PfPOU5F3
AoPOU5F3
SslPou5f3.2
SslPOU5F3.1
AmxPOU5F3
CbPOU5F3
PhPOU5F3
DrPOU5F3
LoPOU5F3
AsPOU5F3
Low
High
Rate of POU protein evolution
