## Supplementary Information 2 for "Evolutionary Origin of Vertebrate OCT4/POU5 Functions in Supporting Pluripotency"

**Additional information for figures**

### CellProfiler Pipeline for immunostaining quantification (Related to Fig. 3b, c)

#copy this text to .txt or .rtf file#

```
Version:5
DateRevision:413
GitHash:
ModuleCount:7
HasImagePlaneDetails:False

Images:[module_num:1|svn_version:'Unknown'|variable_revision_number:2|show_window:Fals
e|notes:['To begin creating your project, use the Images module to compile a list of files and/or
folders that you want to analyze. You can also specify a set of rules to include only the desired
files in your selected folders.']|batch_state:array([],
dtype=uint8)|enabled:True|wants_pause:False]
    :
    Filter images?:Images only
    Select the rule criteria:and (extension does isimage) (directory doesnot containregexp
"[\\\\/]\\.")

Metadata:[module_num:2|svn_version:'Unknown'|variable_revision_number:6|show_window:Fal
se|notes:['The Metadata module optionally allows you to extract information describing your
images (i.e, metadata) which will be stored along with your measurements. This information can
be contained in the file name and/or location, or in an external file.']|batch_state:array([],
dtype=uint8)|enabled:True|wants_pause:False]
    Extract metadata?:No
    Metadata data type:Text
    Metadata types:{}
    Extraction method count:1
    Metadata extraction method:Extract from file/folder names
    Metadata source:File name
    Regular expression to extract from file name:^(?P<Plate>.*)_(?P<Well>[A-P][0-
9]{2})_s(?P<Site>[0-9])_w(?P<ChannelNumber>[0-9])
    Regular expression to extract from folder name:(?P<Date>[0-9]{4}_[0-9]{2}_[0-9]{2})$
    Extract metadata from:All images
    Select the filtering criteria:and (file does contain "")
    Metadata file location:Elsewhere...|
    Match file and image metadata:[]
    Use case insensitive matching?:No
    Metadata file name:None
    Does cached metadata exist?:No

NamesAndTypes:[module_num:3|svn_version:'Unknown'|variable_revision_number:8|show_win
dow:False|notes:['The NamesAndTypes module allows you to assign a meaningful name to
```

each image by which other modules will refer to it.']|batch_state:array([], dtype=uint8)|enabled:True|wants_pause:False]
    Assign a name to:All images
    Select the image type:Color image
    Name to assign these images:nuclei
    Match metadata:[]
    Image set matching method:Order
    Set intensity range from:Image metadata
    Assignments count:1
    Single images count:0
    Maximum intensity:255.0
    Process as 3D?:No
    Relative pixel spacing in X:1.0
    Relative pixel spacing in Y:1.0
    Relative pixel spacing in Z:1.0
    Select the rule criteria:and (file does contain "")
    Name to assign these images:DNA
    Name to assign these objects:Cell
    Select the image type:Grayscale image
    Set intensity range from:Image metadata
    Maximum intensity:255.0

Groups:[module_num:4|svn_version:'Unknown'|variable_revision_number:2|show_window:False|notes:['The Groups module optionally allows you to split your list of images into image subsets (groups) which will be processed independently of each other. Examples of groupings include screening batches, microtiter plates, time-lapse movies, etc.']|batch_state:array([], dtype=uint8)|enabled:True|wants_pause:False]
    Do you want to group your images?:No
    grouping metadata count:2
    Metadata category:None
    Metadata category:None

ColorToGray:[module_num:5|svn_version:'Unknown'|variable_revision_number:4|show_window:True|notes:[]|batch_state:array([], dtype=uint8)|enabled:True|wants_pause:False]
    Select the input image:nuclei
    Conversion method:Combine
    Image type:RGB
    Name the output image:OrigGray
    Relative weight of the red channel:1.0
    Relative weight of the green channel:1.0
    Relative weight of the blue channel:1.0
    Convert red to gray?:Yes
    Name the output image:OrigRed
    Convert green to gray?:Yes
    Name the output image:OrigGreen
    Convert blue to gray?:Yes
    Name the output image:OrigBlue
    Convert hue to gray?:Yes
    Name the output image:OrigHue

```
    Convert saturation to gray?:Yes
    Name the output image:OrigSaturation
    Convert value to gray?:Yes
    Name the output image:OrigValue
    Channel count:1
    Channel number:1
    Relative weight of the channel:1.0
    Image name:Channel1

IdentifyPrimaryObjects:[module_num:6|svn_version:'Unknown'|variable_revision_number:14|show_window:True|notes:[]|batch_state:array([], dtype=uint8)|enabled:True|wants_pause:False]
    Select the input image:OrigGray
    Name the primary objects to be identified:IdentifyPrimaryObjects
    Typical diameter of objects, in pixel units (Min,Max):15,50
    Discard objects outside the diameter range?:No
    Discard objects touching the border of the image?:No
    Method to distinguish clumped objects:Intensity
    Method to draw dividing lines between clumped objects:Intensity
    Size of smoothing filter:10
    Suppress local maxima that are closer than this minimum allowed distance:7.0
    Speed up by using lower-resolution image to find local maxima?:Yes
    Fill holes in identified objects?:After declumping only
    Automatically calculate size of smoothing filter for declumping?:Yes
    Automatically calculate minimum allowed distance between local maxima?:Yes
    Handling of objects if excessive number of objects identified:Continue
    Maximum number of objects:500
    Display accepted local maxima?:No
    Select maxima color:Blue
    Use advanced settings?:Yes
    Threshold setting version:12
    Threshold strategy:Global
    Thresholding method:Otsu
    Threshold smoothing scale:1.3488
    Threshold correction factor:1.0
    Lower and upper bounds on threshold:0.0,1.0
    Manual threshold:0.0
    Select the measurement to threshold with:None
    Two-class or three-class thresholding?:Three classes
    Log transform before thresholding?:No
    Assign pixels in the middle intensity class to the foreground or the background?:Foreground
    Size of adaptive window:50
    Lower outlier fraction:0.05
    Upper outlier fraction:0.05
    Averaging method:Mean
    Variance method:Standard deviation
    # of deviations:2.0
    Thresholding method:Minimum Cross-Entropy
```

```
MeasureObjectSizeShape:[module_num:7|svn_version:'Unknown'|variable_revision_number:3|show_window:True|notes:[]|batch_state:array([], dtype=uint8)|enabled:True|wants_pause:False]
    Select object sets to measure:IdentifyPrimaryObjects
    Calculate the Zernike features?:Yes
    Calculate the advanced features?:No
```

#copy this text to .txt or .rtf file#

**Gating strategy (Related to Fig. 4d-e, Supplementary Fig. 4b)**

FSC-A/SSC-A gates were set to select cell population of interest and exclude cell debris. FSC-H/FSC-W gates were set to select single cell population. DAPI-A/FSC-A gates were used to select only live cells and exclude dead cells (high DAPI). This DAPI low/negative gates were then used to determine cell populations with/without c-KIT-APC and Nanog-eGFP. Boundary between positive and negative cell populations were set using unstained E14Tg2A ESCs.

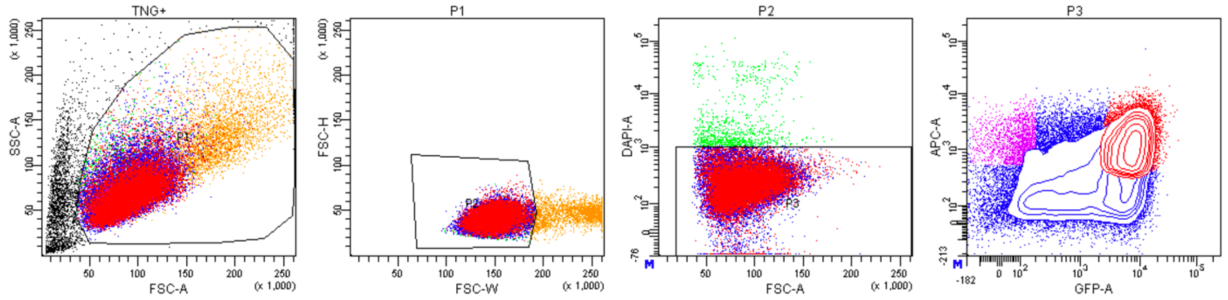


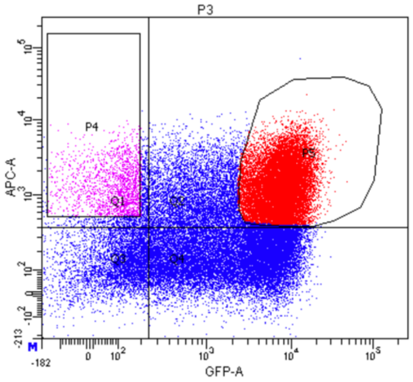


Tube: TNG+

| Population | #Events | %Parent | %Total |
|---|---|---|---|
| All Events | 57,657 | #### | 100.0 |
| P1 | 54,815 | 95.1 | 95.1 |
| P2 | 50,882 | 92.8 | 88.2 |
| P3 | 50,000 | 98.3 | 86.7 |
| Q1 | 2,535 | 5.1 | 4.4 |
| Q2 | 23,808 | 47.6 | 41.3 |
| Q3 | 4,591 | 9.2 | 8.0 |
| Q4 | 19,066 | 38.1 | 33.1 |
| P4 | 1,691 | 3.4 | 2.9 |
| P5 | 17,542 | 35.1 | 30.4 |

**Expression of *ScPou5f1* and *ScPou5f3* during catshark early development (Related to Fig. 6b)**

The catshark develops from a telolecithal egg and exhibits a protracted development, gastrulation movements starting at stage 11, seven to ten days after blastocoel formation, which itself takes place at the time of egg deposition. Prior to gastrulation (blastocoel stage to stage 10), the blastoderm consists of two cell populations, a deep mesenchymal one of endodermal identity, and a superficial, epithelial-like layer, expressing *Sox2* and *Bmp4* in opposite, largely complementary territories[71,89] At these stages, both *ScP1* and *ScP3* show similarly strong, uniform expression in the superficial cell layer, including the blastoderm margin but excluding the deep mesenchyme (Fig. 6b i-ii, ii$^1$, viii-ix). Expression restricted to the superficial layer including the margin is maintained for both paralogues at stage 11, when involution starts posteriorly (Fig. 6b iii, x, x$^1$). When the embryonic axis becomes morphologically visible, at the posterior pole of the blastoderm (stage 12), the signal intensity decreases in a sector located anteriorly to the forming neural plate, while increasing posteriorly, at the level of the forming posterior arms where mesoderm internalisation takes place (Fig. 6b iv, xi). During axis elongation (stage 13-14), the transcripts become confined to the posterior part of the embryonic axis, including the posterior arms but excluding anterior territories, which start to express lineage specific markers (Fig. 6b v-vi, v$^1$, v$^2$, xii-xiii, xiii$^1$; ref[47]). The expression profiles of the two paralogues segregate at the time of neural tube closure (Fig. 6b vii, xiv). From stage 15 to 19, *ScP1* expression is consistently observed in the posterior lateral mesoderm, at the site expected for primordial germ cells (PGCs) (Fig. 6b vii, vii$^1$, vii$^2$), while *ScP3* only exhibits a faint and transient signal at this location at stage 15. *ScP3* expression sites not shared by *ScP1* are also observed at these stages, in the posterior part of the head enlargement at stage 15 (Fig. 6b xiv, xiv$^1$), in the anterior hindbrain, with two bands of expression at stage 16 (Fig. 6b xv, xv$^1$), and at the level of the tail bud at stages 16 to 19 (Fig. 6b xv, xv$^2$, xvi, xvi$^1$).

**Additional Fig. 6b legend**

Whole-mount views of catshark embryos following in situ hybridisations with probes for *ScPou5f1* (*ScP1*) (**i-vii**) or *ScPou5f3* (*ScP3*) (**viii-xvi**). **i-iii** and **viii-x** show dorsal views of the blastoderm at blastula (**i, viii**) and epiboly (**ii, ix**) stages, and at the onset of gastrulation (stage 11;

**iii, x**). **iv-vii** and **xi-xiv** show dorsal views of the elongating embryo at stage 12 (**iv, xi**), stage 13 (**v, xii**), stage 14 (**vi, xiii**), stage 14+ (**xiv**) and stage 15 (**vii**). **xv** and **xvi** show lateral views of stage 19 and 20 embryos respectively. **ii**$^{1}$, **v**$^{1}$, **v**$^{2}$, **x**$^{1}$, **xiii**$^{1}$, **xiv**$^{1}$ and **xvi**$^{1}$ show sections of the embryos presented respectively in **ii**, **v**, **x**, **xiii**, **xiv** and **xvi**, at the plane and level indicated by red dotted lines. **vii**$^{1}$ shows enhanced magnification of the territory boxed in red in **vii**. **vii**$^{2}$ shows a transverse section at the level of presumptive PGCs of a stage 19 embryo, submitted to whole-mount hybridisation with a *ScP1* probe. Red arrows in **iv-vi** and **xi-xiii** point to a signal at the posterior margin lining the posterior arms. Red arrowheads in **vii**, **vii**$^{1}$, **vii**$^{2}$, **xiv** point to signals observed at the expected location for PGCs. **xv**$^{1}$ and **xv**$^{2}$ respectively show magnifications of anterior hindbrain and tailbud territories boxed in red in **xv**, where *ScP3* but not *ScP1*, is expressed. The red asterisk in **xvi**, **xvi**$^{1}$ and the red arrow in **xv**$^{1}$ and **xvi** respectively indicate *ScP3* signals in the tailbud and in the anterior hindbrain, not observed for *ScP1*.
