## Supplementary Information 3 for "Evolutionary Origin of Vertebrate OCT4/POU5 Functions in Supporting Pluripotency"

### Supplementary Information 3
### Structural models of POU5 homologues from AlphaFold2

#### Table 1. Protein sequences used for AlphaFold2 structural modelling

| Oct4 homologs | Protein sequences |
|---|---|
| **mOct4** | UniProt : P20263 https://alphafold.ebi.ac.uk/entry/P20263 |
| **MeP1** | MAGHLAPEYFSPPPGGGGAGSSEPTWNWPGFQGPPSGAGAGPGPDGWGMAPYHPLYDMWGGGVYCEPQPNVGVMAQPAQEIAVPDGEA<br>GPGVESPSEDSSPESRATVRVTKIEPVESGEEQPQETPSPEELEQFAKELKRKRITLGYTQADVGITLGALFGKVFSQTTICRFEAQQ<br>LSFKNMCKLRPLLQKWLEAADDNDHLQELCKAETVLQQARKRKRTSIENGVRGNLETMFLQCPKPTLQQISNIAEELGLEKDVVRVWF<br>CNRRQKGKRSNSNSPREDFEAAGSFPGGPMPFPLAPGPPPFSHPSYGGPHFTALYSPSPFTEGDAFSSLPVTPLGSAMHSS |
| **MeP3** | MPDGSSQYGKGGYIGGSTPRPSQPQPSSQSFFTFPSAVKSEYSLEGPSTPDSSQTKGWYAFSAPPPPAEAGQASASHNLHLMPSKEED<br>SSCSSPSSSSSGAPERDLRETVVPEAKYCAHPGPTYYAPAWTGPFWPGLGSSGSVPTGALPGTPLPPHPLQALGTFPGPRPLYPASVQ<br>QPQPGLGSSGGTSSGSVSSSNVSSSVSSTSGSSGAASEEGIPSSDSGEEDTPTSEELELFAKELKHKRISLGFTQADVGMALGTLYGK<br>MFSQTTICRFEALQLSFKNMCKLKPLLQRWLQAVENTDNPQEMCSMEQVLAQARKRKRRTSIETSVKGTLEGFFRRCGKPTPQQICDL<br>AEELHLDKDVVRVWFCNRRQKGKRLLLPYGEDGEALPYELAPGTALVLPTAAVPQNYAAPPPPEPPALYMSTFPKGEPCIPGMPMPNG<br>GI |
| **CpP1** | MAGHVGQELGRPYPLAGQGLHLDAVPPGPANGAPHEGPGSFLPEAYNGAAPGQGYGYRLDYGPQGGLLETQPGDPPGHQALLRAWCPF<br>PMGGEGWPPPYGPAQPAAAYEGLKPDVKTERECGQQGPMYGHPAQAWGGCFLPQATARPPAALPPPPAGPAGEGGEGSGASSPQSDGS<br>GGSPGAPAAPGEETPSTEEMEQFAKELKHKRITLGFTQADVGLALGVLYGKMFSQTTICRFEALQLSFKNMCKLKPLLQRWLDEADGN<br>ANLQEMCSMESALLQARKRKRTSIETAARGSLESYFLRCPKPSLQEIAHIAHDLHLDKDVVRVWFCNRRQKGKRSGGCSVRDDCEGGA<br>LPFAPPAQLPGPPMGHHPPPPQGYNAAAFATLYVPQFHHGGEPFDPTPPGPPLMGHPMHST |
| **CpP3** | MFSQEALPAASFSLGAGILQDASGQFGKSSYSSSPQPPPFFPFPAVKAEYEPPELQAGEAKPWCPFPGPEPCHQPGMAGGHQGPPLLG<br>SGQAPKEEPGQEKGRRQGSPEAKCALLPAGPYYAHPWSSPFWPALAGHSAAAARGPLPSQTFPGVGLCPGAHHAIYPSDPHQPPSGLS<br>SLGSSGSSSGAASEGGQSSESGDEDATPTSEELEQFAKDLKHKRITLGFTQADVGMALGTLYGKMFSQTTICRFEALQLSFKNMCKLK<br>PLLQRWLNEVENSDSLQELCNAEQVLAQARKRKRRTSIENNVKGTLESFFRKCIKPSPQQISQIAEDLNLDKDVVRVWFCNRRQKGKR<br>LLLPFGDENEGAMYDMNLALAHPALPAAVSSYAPAPLASPPPIYMSAFHKGEICPQALQPAVSMGNSGN |
| **PmP1** | MAGHLRRELGRPYSFPVQGLHLTTLPAVPQHETSGVFLPDAFSGATPGQAFGFKPDYGSQGGIVEPYPGGGEVSHSWYPFPAGGEAWN<br>PPPGVVMGPYAVPQPGEACQGSKPDIKVERDFDQPGPYYGGGQPWAGACLVPPAAARPASCSSTKEAASPSPDPQPSSSPASQVEEAD<br>SADSSPRSDASQSPSEQMASEEVKEEAASGDEETTTSVELEQFAKELKHKRITLGFTQADVGLALGLLYGKMFSQTTICRFEALQLSF<br>KNMCKLKPLLQRWLQEADSNENLQEQLCSMESAMIQARKRKRTSIENNVRGSLESYFLRCPKPNLQEISQIADDLSLEKDVVRVWFCN<br>RRQKGKRNPGPGPRDECDGPVLPQVCPHGPLQAPGLGPAHHQLPPPAQGYNTTAFAAVYLPQFHEGDAFVPPTSSSAVMGHPMHSS |
| **AmP1** | MAGHLGQEIGRAAYGFGAQALHLGAGGLEAGGPGFLSESYGPYAGFKALEYAHGGAEGEGRPGAHGLARAWYPFSEAWGPVYGQSGAG<br>AGFESSRVEVKVERPDKEAGYGQQHQQAWAGYFVPQLAVPARSPASVASGGQVPAAPASPSDDSPHSSTASSSSASPDLGAGGAPRDL<br>DSGDEEGGTSADLEQFAKELKQKRITLGFTQADVGLALGALYGKMFSQTTICRFEALQLSFKNMCKLRPLLQRWLVEADTNENLQELC<br>NLENALQQARKRKRTSIENSVKDNLEAFFLKCPKPTHQEIAHISEDLNLEKDVVRVWFCNRRQKGKRSICREEYDGFQQYPGMQPGPP<br>ALSHLPTSYIAQGYNGAAAAFAAVYMQPFHDSEMYSQTVSRHLHSN |
| **AmP3** | MLGRDTVTRDPSPTFAICRTGGAMYSQDPRAPIPMNIQEGSCQATFPGLPRPRPPAQSNTRPLGHPQPFLPFPGVKTPYGTCERAGGG<br>VEPEQARPWHPFQVPEALGPPGISVGHQVDRLGEVRREFKEEPQAEDGCRQGSPEGTYSPPVPTYGGPYYPQPWNGSFWPALGGTGST<br>ANCSPGSAIPVPPGLYPSPLNQSSSGVSSLGSSSSEATSEGGLSSDSGDEDTPTNEELEQFAKALKHKRITLGFTQADVGLALGSLYG<br>RMFSQTTICRFEALQLSFKNMCKLKPLLQRWLNEAENTDNMEELCNMEQMLAQARKRKRRTSIENNVRGTLESFFLKCSKPGPQEISQ<br>IAEDLSLDKDVVRVWFCNRRQKGKRLLLPFVEEMEGGGMYETNQAMAHPGGAPFTLPTMISSQGYPVSSLNSQTLYMTAFHKTEMFPQ<br>ALHPGVPLGNSIS |
| **X91** | MYNQQTYPSFTHNPALMPDGSGQYNLGTYTGMARHPHQAQAFFPFSGVKSDYGDLGGQTTSVGDTSAWNPLTSLDSANQLGISGQGNP<br>FKNLKREREDDEEKSESPEPKCSPPSLPPAYYTHAWNPTTTFWSQVSSSGTTVVSKPLPTPLQPGDKCDPVEANKIFTSSPDKSGESG<br>ISSLDNSRCSSATSSSSGGTNVGTPRSLSRGASDGLSSDSEEEAPNSGEMEQFAKDLKHKRITMGYTQADVGYALGVLFGKTFSQTTI<br>CRFESLQLSFKNMCKLKPLLRSWLHEVENNENLQEIISRGQIIPQVQKRKHRTSIENNVRCTLENYFMRCSKPSAQEIAQIARELNME<br>KDVVRVWFCNRRQKGKRQVYPYIRENGGEPYDTPQTLTPPSQGPFPLPQVMPSQVFPTVPLGANPTIYAPTYHKNDMFPQAMHHGIGM<br>GNQGN |
| **X25** | MYSQQPFPAFAFNAGLMQDPANCHFGGYTGLGHPQPFSFAFSTLKSENGESGVQGMGDCTTPVMPWNSLASFDHQVQMENNQQGNPPR<br>APSPTLSDSRIKVKEEVVHETDSGEESPEPKYPSPPNPSLYYPNAWTGAPFWQVNPTPGNNINPMPNQTLVKNTSLPGNTTYPTPANQ<br>SPNTPVDCVASSMESSRCSSTNSPNGAINERATTIPNGEMLDGGQSSDNEEEVPSESEMEQFAKDLKHKRVSLGYTQADVGYALGVLY<br>GKMFSQTTICRFESLQLSFKNMCQLKPFLERWVVEAENNDNLQELINREQVIAQTRKRKRRTNIENIVKGTLESYFMKCPKPGAQEMV<br>QIAKELNMDKDVVRVWFCNRRQKGKRQGMPTVEENDGEGYDVAQTMGSPPVGHHALQQVVTPQGYMAAPQIYASAFHKNDLFPQTVPH<br>GMAMGGHIG |
| **LcP1** | MAGHLGQDVGRSYGMCPEVFQVSGGTRQEASSVLGADGASLFSPPGFAFKEEYGHAAVDTQFGDPATMAGAHALSRPWYPFPAAEAWA<br>HGAIMTGQHSTQAPHSSHDIHKVHIKTEEEGRSEAKTSTPVSQGVLHGHPLWSPRFLPQVPSGGLVANSLVSARTLGGHGYWASQQAT<br>QSRPPTSPVSGKSSPSDQPTSPENAEENHRDALTDSGTEDTPTTDDLEQFAKELKHKRISLGFTQADVGLALGALYGKMFSQTTICRF<br>EALQLSFKNMCKLKPLLQRWLDEADTNENLQELCNLEQVLSQARKRKRTSLETTAKGTLESFFLKCSKPSLQEIAQIAEELSLDKDVV<br>RVWFCNRRQKGKRSLYQSEEEYETPPQYGVHPFQPPVPTMHLPNSVVQQAYNGTPFPTTLYVPQFYEGEAFPQPRAGPCTRNEFWDST<br>PFSTSF |
| **LcP3** | MSDRSTTPNQDGSRVCRPIYPQEGLSSLNLGNGMLQDSSSQFSKSNYNGISPQPFFPFPPMKADYGQIMELQVGDSGHSRHWYAFPAP<br>DLSSQLALGGQQAGGHSSQLGEIRDPAKVEIKQEKESPEPKYVSPPAAAAIAAGAYYAHPWNNSFWPALTSSSATTGSSNPIPGQAFP<br>GIGVFPGTSAQLYPTALNHTSSGVSSLASSSSSSSGAGSEEGQSSDSADEDNPTTEELEQFAKELKHKRITLGFTQADVGLALGTLYG<br>KMFSQTTICRFEALQLSFKNMCKLKPLLQRWLNEAENNENLQEMCNIEQVLAQARKRKRRTSIENNVKGTLENYFLKCPKPTSQEISQ |

| | |
|---|---|
| | IADDLNLEKDVVRVWFCNRRQKGKRLAFPFGEEAEGISYDVNPALHAPSTSSMTLHGPVVTQGYPGTTLTAPPPVYMPAFHKSEVFPQ<br>ALHPGVAMTNHTT |
| **RtP1** | MNTAVCAEPLGHRAVPEPSSQATLPAPPAFPLPGDYRLRQFREPSSRMATPWYPESWAPGPPSSSEEEPRRRAQPQVPEVKSPSPALH<br>AGWRSPLPAPSCCFLPPAGYPLPGQPALCPQQEHPARDPPGEQPRSRSRSPGRPQEEEEEAAAGAGSPPDTEGEETPTSEDLKHFAKK<br>LKKKRIIMGFTQAEVGLALGALYGKMFSQTTICRFEALQLSYKNMCKLRPLLQRWLEEAKDNENFQELCSMEQRLATARKRKRRTSID<br>SNVKGLLEAAFIKCPKPSAQEIIQIADDLNLEKDVVRVWFCNRRQKGKRALFHGGEDSEGVPGYGPLPMQHAPLGLPSPLAPHQGYVG<br>EAIGTVYTPHFHKGSLYRQQPAPAPGTMHSS |
| **RtP3** | MSISPGQGANGAKSIQSPERAALVPFGNGVVQDVGAQFYKPGYNGLSAQYLFPFPPLKGEYGHSESPLGDCAAVSPSGYWYPFPGAEP<br>VPTHGAGHNSQVAVAGGYINRPEIKTEKESRGYPQEVRYSSPSGPGPSYNSRWSSPFWQPALAASSSNSSPGSSSSSGPLPSQSYGPF<br>PSPPQLYPSPSQHSDSSQPLQSNTGSTGTASEEGQSSDSEEEYPTKEKMEQFAKELKHKRITLGFTQADVGLALGNLYGKMFSQTTIC<br>RFEALQLSFKNMCKLKPILQRWLNDAENNGSLHEICNVEQVLDQSRKRKRRTSIENGVKRNLETYFMKCPKPTSEEISQIAEDLCLDK<br>EVIRVWFCNRRQKGKRMTLPCMEENDVQIPEGSPLHMSPGALMLPEPTVTQGYSAPMVPPPMYMSPFPQALHPAVSMGNHPS |
| **LeP1** | MTSRIQHEIHRSFPPLFHDPRQLQSQEMSSQLAPERPSIVQGLHCGPAFRGDCQGHMAPEYRLGEPGTAGVSPHLPRPWYPFPAPDHW<br>PHSSVVARYPNAPPAPGREEDEKGTEAKFLPSLYNSPWSSCYLPQLPAAPSPPTARTAAGLQNQGHSPSSDHRSQPASPQMQPNNPSP<br>SPSADHDTKWVAPDGGNEEYPTKEKMEQFAKELKHKRITLGFTQADVGLALGNLYGKMFSQTTICRFEALQLSYKNMCKLKPLLQRWL<br>EEAKDSDNFQELCSIEQTLASSRKRKRRTSIDNNVKEGLENFFTVCPKPSTQEITKIADDLNLEKDVVRVWFCNRRQKGKRAAFQYGD<br>EFVELAGYGMPPLQPPLPGGFGAQGYNAAAISAAALYVPQADSYHCSEAAQAAPVARTMHSS |
| **LeP3** | MSGTLSPVHGPPVTHRVNQSPESAALPAFSSGVVQDMGSQMYKPGYSSMGGQYLFPLGPRKSEYGGCDSAALPHPGYWHQPPVAEPPS<br>TAHWPQGHPHSGHSAHIKVEKDGFSSPPHPSYPHRWCGPGFWPPVPGSSSPASGSGQPCPGYSSFPSPPHMFPSPSQLSDSNPATQSI<br>AGSTGNTSEEGQSSDSEEEYPTKEKMEQFAKELKHKRITLGFTQADVGLALGNLYGKMFSQTTICRFEALQLSFKNMCKLKPILQRWL<br>NDAESDGGIHEICNVEQVLDQSRKRKRRTSIENGVKNNLETYFMKCPKPTSEEISQIAEDLRLDKEVIRVWFCNRRQKGKRMTLPCME<br>ENELQINEESPIRLSPGSLMPDPCVPQGYVMPPHMYMSPFPQPLHPAVTMGNHSS |
| **ScP3** | MSARPGQGTSVSRSVPSPERPPVLSFGNGVVPDVSPQFYKPGYNAISAQYLFPFPGLKGEYGHSETQLGDCAAVSHTGYWYPFTDPAA<br>AHGSGHSSGHPTQLSAPGGLLRPEIKTEKECKGYPQEGRYISPTAPAPSYNHRWSTTFWQPALSSAANSTSSSSSSSSSAPLPSQTYS<br>AFGSFPSPPQMYPTNPSPSQQSDSSQPAHNTGSTGTTSEEGQSSDSEEEYPTKEKMEQFAKELKHKRITLGFTQADVGLALGNLYGKM<br>FSQTTICRFEALQLSFKNMCKLKPILQRWLNDAENNGGLHEICNVEQVLDQSRKRKRRTSIENGVKRNLETYFMKCPKPTSEEISQIA<br>EDLQLDKEVIRVWFCNRRQKGKRMTLPCMEENDVQIHEGSPLHMSPNALMLPDPIVTQGYSAAMVPPPMYMSPFPQALHPAVSMGNHT<br>S |
| **EbP5** | MSSLSIKVDAANELQLMYESYSEASAAQSSVVRPYMEQAAQQRGACCVPGQGSGHGHVHLVPISPLSLGYSGELYEPSLQCPSRAHGA<br>VPGPAWYAYPGPSAAGSEAAQAAASWAAARGVEIKCEEGVGVGEHVESEVKRIRYQDFKYGAVYGSGYVLPNAAMHGGHGQQNPHSLS<br>HLNLHPHLQLNPSPHLQPNPNLHLQLNPNSHLQPSSHLQPNPSSHLQPNPNSHLQLSSHLQQNSHLQPSSHLQPNSHLQPNPNSHLQP<br>NPNSHLQPNTNSHLQPNPNSHLQPNPNSHLQPNPNSHLQLSSHSQQNPLSLLNVHPQAYPHSQSPAPTPLAAALASTQSPGASSGTED<br>GGCARERSRNSEHDLGSGDEGTISSETLAQFARDLKHKRITLGFTQADVGVALGSLYGRIFSQTTICRFEALQLSYRNMCKLQPLLER<br>WMIEAENADNVKEVCHLFLFNSSMDQSLANVNKLRRRRTTIENGVRDTLEAWYLVCSKPSAKEIARIAGELNLDKEVVRVWFCNRRQK<br>LKQLSPPLPKEEEPEMSSQHQLMPHIQEFGTLPTSSSSPAYDIHTYNMQASMNAPILPS |
| **EbP5$^{S4L}$** | MSSLSIKVDAANELQLMYESYSEASAAQSSVVRPYMEQAAQQRGACCVPGQGSGHGHVHLVPISPLSLGYSGELYEPSLQCPSRAHGA<br>VPGPAWYAYPGPSAAGSEAAQAAASWAAARGVEIKCEEGVGVGEHVESEVKRIRYQDFKYGAVYGSGYVLPNAAMHGGHGQQNPHSLS<br>HLNLHPHLQLNPSPHLQPNPNLHLQLNPNSHLQPSSHLQPNPSSHLQPNPNSHLQLSSHLQQNSHLQPSSHLQPNSHLQPNPNSHLQP<br>NPNSHLQPNTNSHLQPNPNSHLQPNPNSHLQPNPNSHLQLSSHSQQNPLSLLNVHPQAYPHSQSPAPTPLAAALASTQSPGASSGTED<br>GGCARERSRNSEHDLGSGDEGTISSETLAQFARDLKHKRITLGFTQADVGVALGSLYGRIFSQTTICRFEALQLSFKNMCKLKPLLQR<br>WLDEADTNENLQELCNLEQVLSQARRRRTTIENGVRDTLEAWYLVCSKPSAKEIARIAGELNLDKEVVRVWFCNRRQKLKQLSPPLPK<br>EEEPEMSSQHQLMPHIQEFGTLPTSSSSPAYDIHTYNMQASMNAPILPS |
| **EbP5$^{H1H2}$** | MSSLSIKVDAANELQLMYESYSEASAAQSSVVRPYMEQAAQQRGACCVPGQGSGHGHVHLVPISPLSLGYSGELYEPSLQCPSRAHGA<br>VPGPAWYAYPGPSAAGSEAAQAAASWAAARGVEIKCEEGVGVGEHVESEVKRIRYQDFKYGAVYGSGYVLPNAAMHGGHGQQNPHSLS<br>HLNLHPHLQLNPSPHLQPNPNLHLQLNPNSHLQPSSHLQPNPSSHLQPNPNSHLQLSSHLQQNSHLQPSSHLQPNSHLQPNPNSHLQP<br>NPNSHLQPNTNSHLQPNPNSHLQPNPNSHLQPNPNSHLQLSSHSQQNPLSLLNVHPQAYPHSQSPAPTPLAAALASTQSPGASSGTED<br>GGCARERSRNSEHDLGSGDEGTISSETLAQFARDLKHKRITLGFTQADVGVALGSLYGRIFSQTTICRFEALQLSYRNMCKLQPLLER<br>WMIEAENADNVKEVCHLFLFNSSMDQSLANVNKLRKRKRTSLETTAKGTLESFFLKCSKPSLQEIAQIAEELNLDKEVVRVWFCNRRQ<br>KLKQLSPPLPKEEEPEMSSQHQLMPHIQEFGTLPTSSSSPAYDIHTYNMQASMNAPILPS |
| **EbP5$^{LH2}$** | MSSLSIKVDAANELQLMYESYSEASAAQSSVVRPYMEQAAQQRGACCVPGQGSGHGHVHLVPISPLSLGYSGELYEPSLQCPSRAHGA<br>VPGPAWYAYPGPSAAGSEAAQAAASWAAARGVEIKCEEGVGVGEHVESEVKRIRYQDFKYGAVYGSGYVLPNAAMHGGHGQQNPHSLS<br>HLNLHPHLQLNPSPHLQPNPNLHLQLNPNSHLQPSSHLQPNPSSHLQPNPNSHLQLSSHLQQNSHLQPSSHLQPNSHLQPNPNSHLQP<br>NPNSHLQPNTNSHLQPNPNSHLQPNPNSHLQPNPNSHLQLSSHSQQNPLSLLNVHPQAYPHSQSPAPTPLAAALASTQSPGASSGTED<br>GGCARERSRNSEHDLGSGDEGTISSETLAQFARDLKHKRITLGFTQADVGVALGSLYGRIFSQTTICRFEALQLSYRNMCKLQPLLER<br>WMIEAETNENLQELCNLEQVLSQARKRKRTSLETTAKGTLESFFLKCSKPSLQEIAQIAEELNLDKEVVRVWFCNRRQKLKQLSPPLP<br>KEEEPEMSSQHQLMPHIQEFGTLPTSSSSPAYDIHTYNMQASMNAPILPS |
| **EbP5$^{S4LH2}$** | MSSLSIKVDAANELQLMYESYSEASAAQSSVVRPYMEQAAQQRGACCVPGQGSGHGHVHLVPISPLSLGYSGELYEPSLQCPSRAHGA<br>VPGPAWYAYPGPSAAGSEAAQAAASWAAARGVEIKCEEGVGVGEHVESEVKRIRYQDFKYGAVYGSGYVLPNAAMHGGHGQQNPHSLS<br>HLNLHPHLQLNPSPHLQPNPNLHLQLNPNSHLQPSSHLQPNPSSHLQPNPNSHLQLSSHLQQNSHLQPSSHLQPNSHLQPNPNSHLQP<br>NPNSHLQPNTNSHLQPNPNSHLQPNPNSHLQPNPNSHLQLSSHSQQNPLSLLNVHPQAYPHSQSPAPTPLAAALASTQSPGASSGTED<br>GGCARERSRNSEHDLGSGDEGTISSETLAQFARDLKHKRITLGFTQADVGVALGSLYGRIFSQTTICRFEALQLSFKNMCKLKPLLQR<br>WLDEADTNENLQELCNLEQVLSQARKRKRTSLETTAKGTLESFFLKCSKPSLQEIAQIAEELNLDKEVVRVWFCNRRQKLKQLSPPLP<br>KEEEPEMSSQHQLMPHIQEFGTLPTSSSSPAYDIHTYNMQASMNAPILPS |

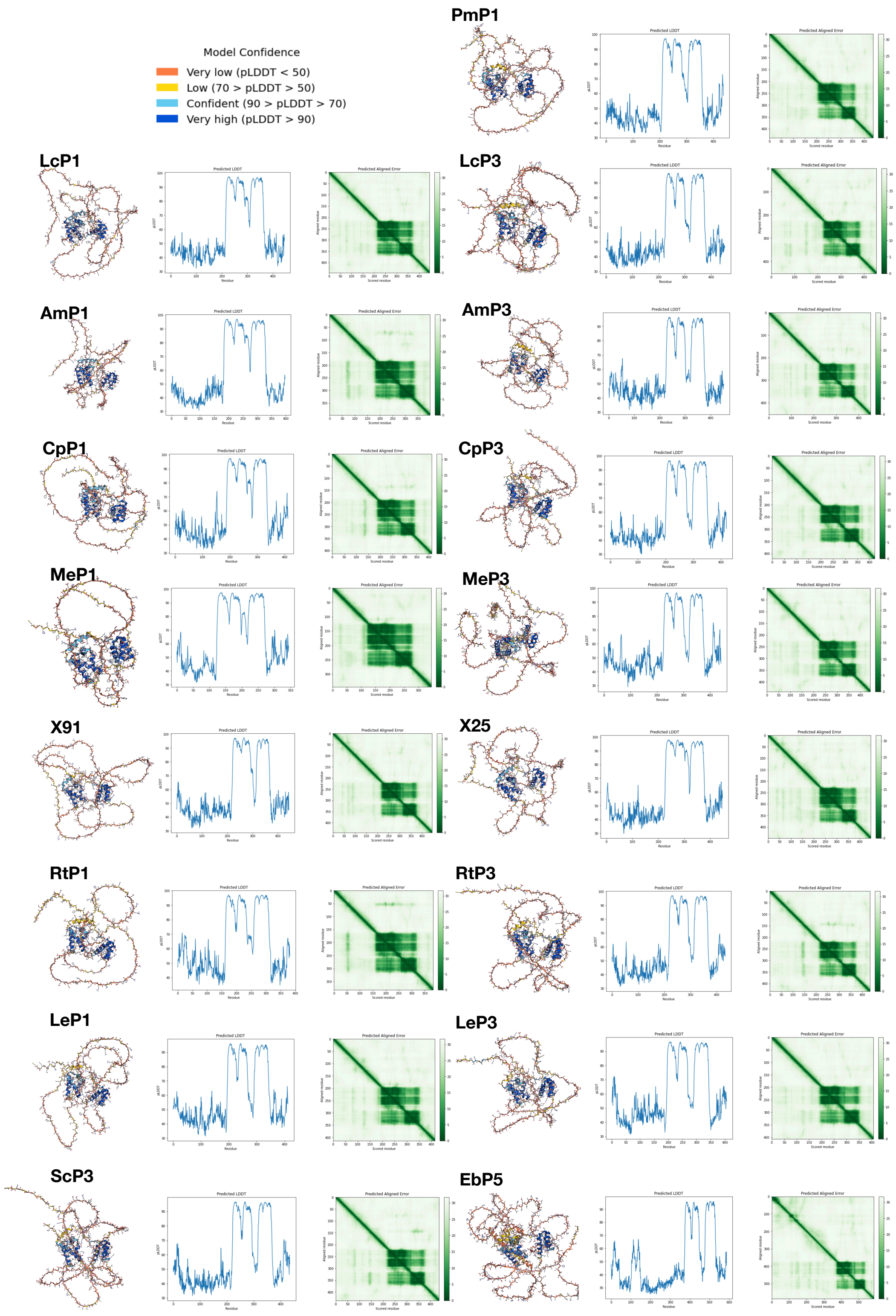


**Fig. 1 Outputs from AlphaFold2-structural modelling of POU5 proteins** including 3D coordinates (left), per-residue confidence metric called pLDDT (middle) and Predicted Aligned Error (right)

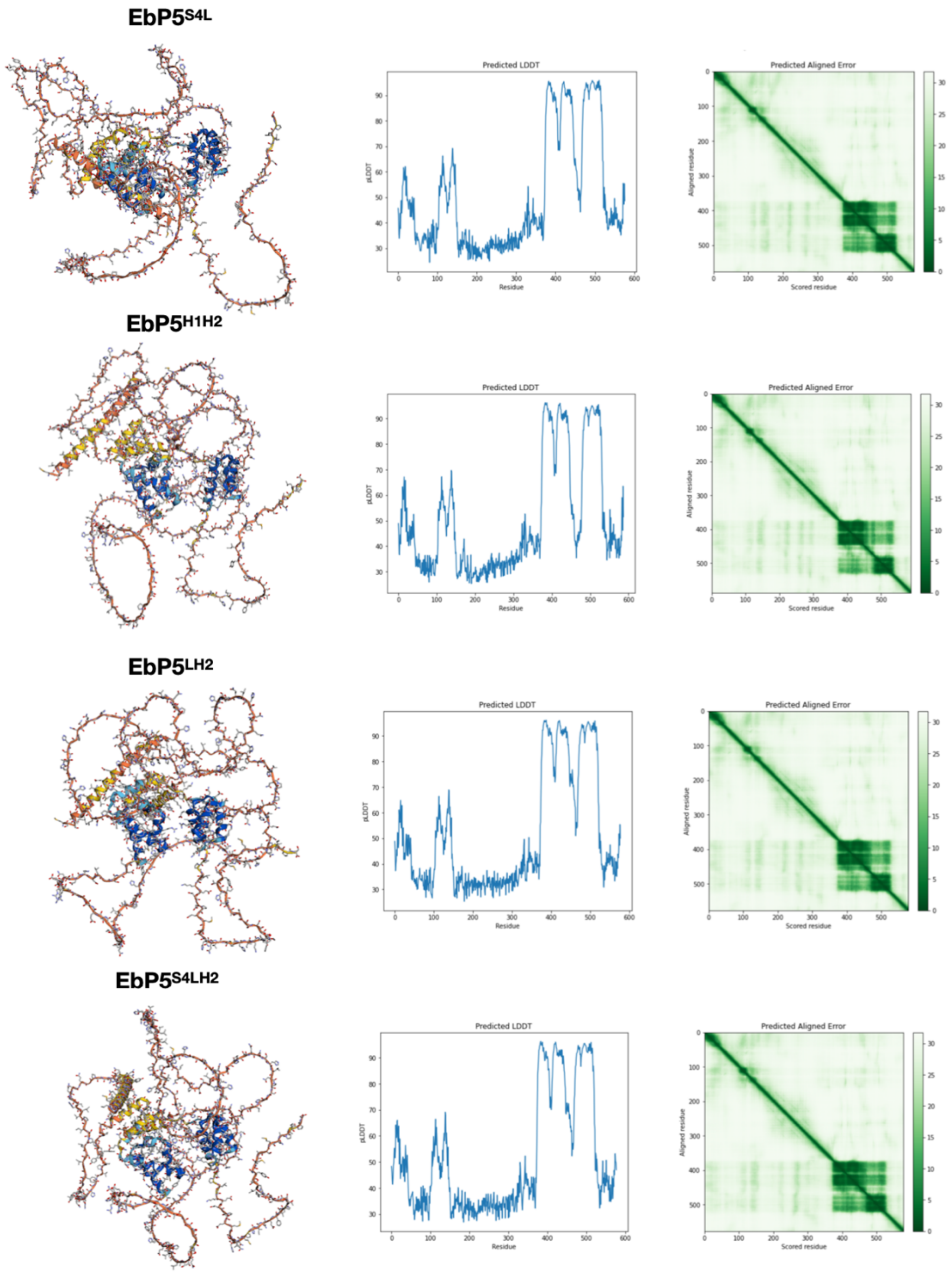


**Fig. 2 Outputs from AlphaFold2-structural modelling of hagfish-coelacanth chimeric POU5 proteins** including 3D coordinates (left), per-residue confidence metric called pLDDT (middle) and Predicted Aligned Error (right)

#### Table 2. Structural assessment of isolated POU domains

| Oct4 homologues/ chimeric proteins | | Structural assessment using MolProbity on Phenix | | | |
|---|---|---|---|---|---|
| | | Isolated POU-S-L | | Isolated POU-HD | |
| | | Ramachandran favoured | Clash score | Ramachandran favoured | Clash score |
| **x-ray diffraction PDB: 3L1P** (Esch et al., 2013) | **mOct4** | 89.02 | 4.95 | 93.75% | 0.00 |
| **AlphaFold2** (in this study) | **mOct4** | 98.90% | 1.96 | 100.00% | 0.00 |
| | **MeP1** | 100.00% | 2.63 | 100.00% | 0.00 |
| | **MeP3** | 98.91% | 2.66 | 100.00% | 1.18 |
| | **CpP1** | 98.90% | 1.36 | 100.00% | 1.20 |
| | **CpP3** | 90.22% | 0.00 | 95.92% | 0.00 |
| | **PmP1** | 96.81% | 0.00 | 97.96% | 0.00 |
| | **AmP1** | 97.83% | 1.34 | 100.00% | 0.00 |
| | **AmP3** | 90.22% | 0.00 | 95.92% | 0.00 |
| | **X91** | 89.53% | 0.00 | 97.96% | 0.00 |
| | **X25** | 92.22% | 0.00 | 95.92% | 0.00 |
| | **LcP1** | 89.13% | 0.00 | 89.80% | 1.19 |
| | **LcP3** | 92.22% | 0.00 | 93.88% | 0.00 |
| | **RtP1** | 100.00% | 0.66 | 100.00% | 0.00 |
| | **RtP3** | 98.90% | 1.33 | 100.00% | 1.17 |
| | **LeP1** | 96.70% | 1.32 | 100.00% | 0.00 |
| | **LeP3** | 95.12% | 0.74 | 100.00% | 0.00 |
| | **ScP3** | 95.45% | 0.00 | 100.00% | 0.00 |
| | **EbP5** | 91.09% | 0.00 | 97.96% | 0.00 |
| | **$EbP5^{S4L}$** | 96.67% | 0.00 | 100.00% | 0.00 |
| | **$EbP5^{H1H2}$** | 97.00% | 3.70 | 100.00% | 0.00 |
| | **$EbP5^{LH2}$** | 97.83% | 0.00 | 100.00% | 0.00 |
| | **$EbP5^{S4LH2}$** | 96.74% | 2.00 | 100.00% | 0.00 |

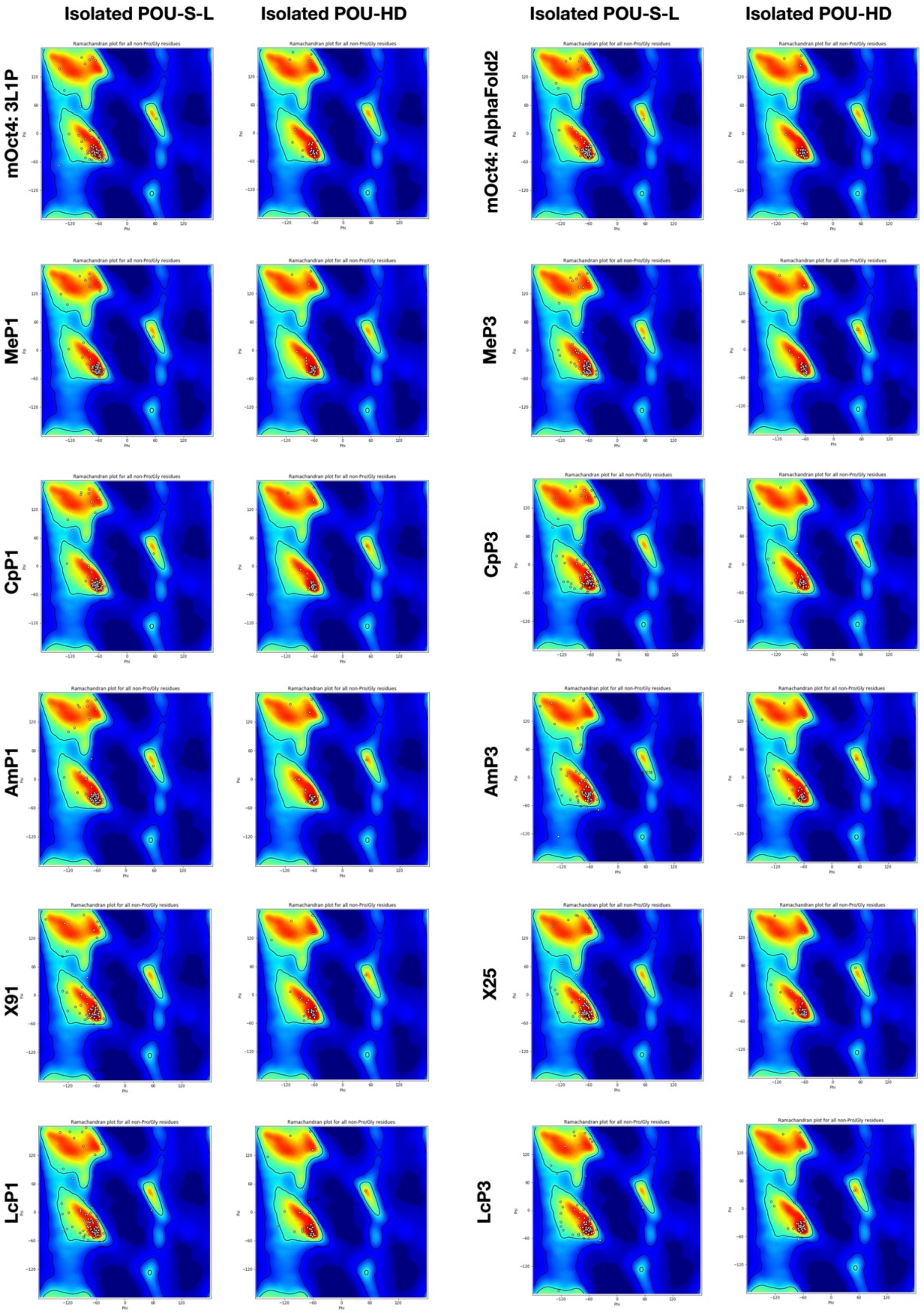


**Fig. 3 Ramachandran plots of POU5 structural models** Isolated POU specific domain including structural linker (POU-S-L) and isolated POU homeodomain (POU-HD) from each Alphafold2-predicted POU5 protein structures were analysed by Phenix.

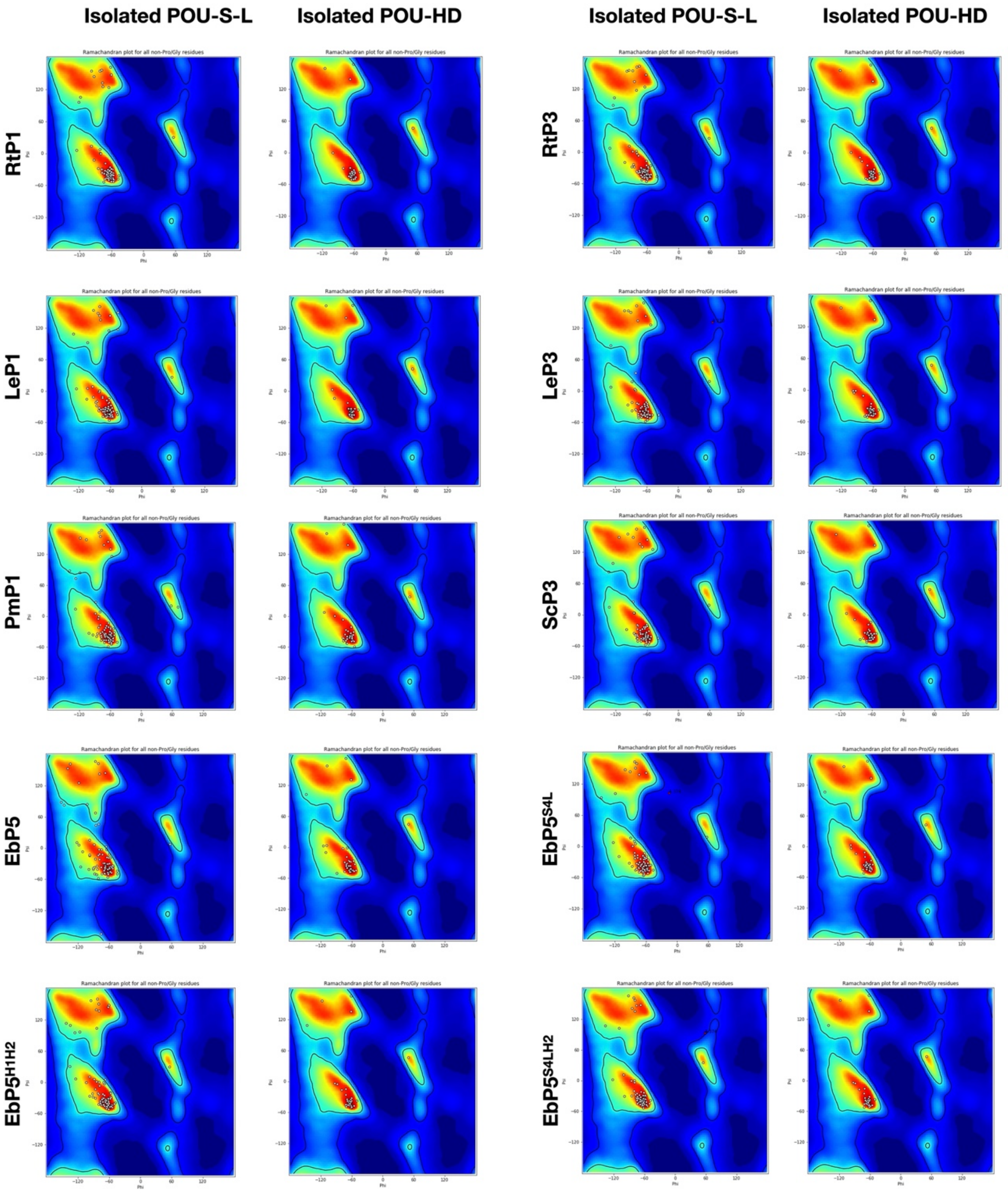


**Fig. 3 (Continued) Ramachandran plots of POU5 structural models** Isolated POU specific domain including structural linker (POU-S-L) and isolated POU homeodomain (POU-HD) from each Alphafold2-predicted POU5 protein structures were analysed by Phenix.

#### Table 3. Structural assessment of isolated POU domains on *PORE* sequence

| **Oct4 homologues/chimeric proteins on *PORE*** (AlphaFold2-based structures) | **Structural assessment using MolProbity on Phenix** | |
|---|---|---|
| | Ramachandran favoured (%) | Clash score |
| **mOct4-*PORE*** | 99.29 | 7.51 |
| **MeP1-*PORE*** | 100.00 | 8.39 |
| **MeP3-*PORE*** | 99.29 | 8.65 |
| **CpP1-*PORE*** | 99.29 | 7.71 |
| **CpP3-*PORE*** | 92.20 | 4.98 |
| **PmP1-*PORE*** | 97.20 | 4.15 |
| **AmP1-*PORE*** | 98.60 | 5.21 |
| **AmP3-*PORE*** | 92.20 | 5.78 |
| **X91-*PORE*** | 92.59 | 7.76 |
| **X25-*PORE*** | 93.53 | 5.81 |
| **LcP1-*PORE*** | 89.36 | 5.79 |
| **LcP3-*PORE*** | 92.81 | 5.81 |
| **RtP1-*PORE*** | 100.00 | 5.99 |
| **RtP3-*PORE*** | 99.29 | 8.64 |
| **LeP1-*PORE*** | 97.89 | 5.20 |
| **LeP3-*PORE*** | 96.95 | 6.26 |
| **ScP3-*PORE*** | 97.06 | 6.89 |
| **EbP5-*PORE*** | 93.33 | 5.07 |
