## Supplementary Table 3 for "Evolutionary Origin of Vertebrate OCT4/POU5 Functions in Supporting Pluripotency"

**Supplementary Table 3. Resources used in and generated by this study**

| Resources | Sources | Catalogue number |
|---|---|---|
| **Chemicals for cell culture** | | |
| Glasgow Minimum Essential Medium (GMEM) | Sigma-Aldrich, Merck | G5154-6X500 ML |
| Dulbecco's Modified Eagle Medium (DMEM)-High Glucose | Invitrogen, ThermoFisher Scientific | 41965-039 |
| Fetal Bovine Serum (FBS) | Invitrogen, ThermoFisher Scientific | 10270-106 |
| KnockOut Serum Replacement | Invitrogen, ThermoFisher Scientific | 10828-028 |
| Non-essential amino acids (NEAA) | Sigma-Aldrich, Merck | M7145 |
| L-glutamine | Invitrogen, ThermoFisher Scientific | 25030024 |
| Sodium pyruvate | Invitrogen, ThermoFisher Scientific | 11360039 |
| β –mercaptoethanol | Sigma-Aldrich, Merck | M6250 |
| Leukemia Inhibitory Factor (LIF) | Homemade (Brickman Lab) | N/A |
| L-ascorbic acid | Sigma-Aldrich, Merck | A4403-100MG |
| Alk5 inhibitor | Tocris | A83-01 |
| Gelatin | Homemade (Brickman Lab) | N/A |
| Tetracycline hydrochloride (Tc) | Sigma-Aldrich, Merck | T-7660 |
| Puromycin dihydrochloride | Sigma-Aldrich, Merck | P8833 |
| N-2 Supplement | homemade | |
| B-27 Supplement | Gibco | 17504001 |
| Neurobasal$^{TM}$ medium | Gibco | 21103049 |
| DMEM/F12 | Gibco | 21331020 |
| PD0325901 | Sigma-Aldrich | PZ0162-5MG |
| CHIR99021 | Axon Med Chem | CT 99021 |
| IWP-2 | Calbiochem | 681671 |
| Activin A | Peprotech | 120-14E |
| bFGF | Peprotech | 450-33 |
| Fibronectin | Millipore Sigma | FC010-1MG |
| **Transgenic mice** | | |
| Nanog-eGFP mice | Ian Chamber Laboratory, University of Edinburgh | N/A |
| 129S2/ScPasCrl | Charles Reiver | N/A |
| **Cell Line** | | |
| *Mus musculus* (house mouse) Embryonic Stem Cell ZHBTc4 containing tetracycline-suppressible Oct4 transgene | Niwa et al., 2000 (ref[27]) | N/A |
| Nanog-eGFP mouse embryonic fibroblasts (MEF) | Homemade | N/A |
| **Antibodies for immunofluorescence, flow cytometry and Western Blots** | | |
| Goat polyclonal anti-mouse Klf4 (Dilution 1:200) | R&D Systems | AF3158 |
| Goat polyclonal anti-human Gata6 (Dilution 1:200) | R&D Systems | AF1700 |

| | | |
|---|---|---|
| Goat polyclonal anti-human/mouse E-cadherin (1:200) | R&D Systems | AF748 |
| Rabbit polyclonal anti-mouse/human Oct4 (1:1000) | Abcam | Ab19857 |
| Mouse monoclonal anti-mouse/rat/human Oct-3/4 (1:200) | Santa Cruz | Sc-5279 |
| Mouse monoclonal anti-Flag (Dilution 1:1000) | Sigma-Aldrich, Merck | F3165 |
| Mouse monoclonal anti-mouse Cdx2 (Dilution 1:100) | BioGenex | MU392A-UC |
| Mouse monoclonal anti-mouse p120 catenin (Dilution 1:250) | BD Transduction Laboratories, BD Biosciences | 610134 |
| Mouse monoclonal anti-mouse Histone H3 (Dilution 1:2000) | Abcam | Ab10799 |
| Rat monoclonal anti-mouse CD31 (PECAM-1) conjugated with APC (Dilution 1:100) | BD Pharmingen, BD Biosciences | 551262 |
| Mouse c-KIT (CD117) conjugated with APC (Dilution 1:500) | BD Pharmingen, BD Biosciences | 561074 |
| Mouse SSEA1 conjugated with Alexa Fluor® 647 (Dilution 1:50) | BD Pharmingen, BD Biosciences | 560120 |
| DAPI | Sigma-Aldrich, Merck | D9542 |
| Donkey anti goat AlexaFluor 488 (IgG) (Dilution 1:800) | Molecular Probes | A11055 |
| Donkey anti goat AlexaFluor 568 (IgG) (Dilution 1:800) | Molecular Probes | A11057 |
| Donkey anti goat AlexaFluor 647 (IgG) (Dilution 1:800) | Molecular Probes | A21447 |
| Donkey anti mouse AlexaFluor 488 (IgG) (Dilution 1:800) | Molecular Probes | A21202 |
| Donkey anti mouse AlexaFluor 568 (IgG) (Dilution 1:800) | Molecular Probes | A10037 |
| Secondary antibody for Western Blots (Dilution 1:2000) | Molecular Probes | N/A |
| **Commercial Assays** | | |
| Alkaline Phosphatase Staining Kit | Sigma-Aldrich, Merck | 86R-1KT |
| Agilent's microarray kit (SurePrint G3 Mouse GE 8X60K Kit; LowInput QuickAmp Labeling Kit One-Color; RNA Spike In Kit-One Color; Gene Expression Hybridization Kit; Pack 5 Backings 8 arrays per slide; Gene Expression Wash Pack) | Agilent Technologies | G4852A; 5190-2305; 5188-5282; 5188-5242; G2534-60014; 5188-532) |
| RNeasy™ Mini Kit | Qiagen | 74106 |
| SuperScript™ III Reverse Transcriptase | Invitrogen, ThermoFisher Scientific | 18080044 |
| LightCycler® 480 Probes Master Mix | Roche Diagnostics | 04902343001 |
| Retro-X™ Concentrator | Clontech | 631453 |
| Retro-X™ qRT-PCR Titration Kit | Clontech | 631456 |
| **Oligonucleotides** | | |
| qPCR primers: *Esrrb* UPL Probe#93<br>Forward: AACTGGGCCAAGCACATC<br>Reverse: ATCTCCATCCAGGCACTCTG | This paper | N/A |
| qPCR primers: *Prdm14* UPL Probe#73<br>Forward: GGCCATACCAGTGCGTGTA<br>Reverse: TGCTGTCTGATGTGTGTTCG | This paper | N/A |
| qPCR primers: *Cdx2* UPL Probe#34<br>Forward: CACCATCAGGAGGAAAAGTGA<br>Reverse: CTGCGGTTCTGAAACCAAAT | This paper | N/A |
| qPCR primers: *Nanog* UPL Probe#25<br>Forward: CCTCCAGCAGATGCAAGAA<br>Reverse: GCTTGCACTTCATCCTTTGG | This paper | N/A |
| qPCR primers: *Fgf5* UPL Probe#95<br>Forward: GCGAAACTTCAGTCTGTACTTCACT<br>Reverse: ACCGGTGAAACCAAAGGTG | This paper | N/A |

| | | |
|---|---|---|
| qPCR primers: *Gata6* UPL Probe#40<br>Forward: GGTCTCTACAGCAAGATGAATGG<br>Reverse: TGGCACAGGACAGTCCAAG | This paper | N/A |
| qPCR primers: *Tbp* UPL Probe#97<br>Forward: GGGGAGCTGTGATGTGAAGT<br>Reverse: CCAGGAAATAATTCTGGCTCA | This paper | N/A |
| qPCR primers: *IRES-PAC* UPL Probe#41<br>Forward: TGGCTCTCCTCAAGCGTATT<br>Reverse: CCCCAGATCAGATCCCATAC | This paper | N/A |
| **Plasmids** | | |
| pCAGIP-3xflag mOct4 (*Mus musculus* POU5F1) | Livigni et al., 2013 (ref[35]) | N/A |
| pCAGIP-empty | Livigni et al., 2013; Morrison and Brickman 2006 (ref[35,38]) | N/A |
| pCAGIP-3xflag XlPOU91 (*Xenopus laevis* POU91)/XlPOU5F3.1, X91 | Livigni et al., 2013 (ref[35]) | N/A |
| pCAGIP-3xflag XlPOU25 (*Xenopus laevis* POU25)/XlPOU5F3.2, X25 | Livigni et al., 2013 (ref[35]) | N/A |
| pCAGIP-3xflag LcPOU5F1 (*Latimeria chalumnae* POU5F1), LcP1 | This paper | Synthetic gene |
| pCAGIP-3xflag LcPOU5F3 (*Latimeria chalumnae* POU5F3), LcP3 | This paper | Synthetic gene |
| pCAGIP-3xflag AmPOU5F1 (*Ambystoma mexicanum* POU5F1), AmP1 | This paper | cDNA from Elly Tanaka |
| pCAGIP-3xflag AmPOU5F3 (*Ambystoma mexicanum* POU5F3), AmP3 | This paper | cDNA from Elly Tanaka |
| pCAGIP-3xflag CpPOU5F1 (*Chrysemys picta* POU5F1), CpP1 | This paper | Synthetic gene |
| pCAGIP-3xflag CpPOU5F3 (*Chrysemys picta* POU5F3), CpP3 | This paper | Synthetic gene |
| pCAGIP-3xflag MePOU5F1 (*Macropus eugenii* POU5F1), MeP1 | This paper | Synthetic gene |
| pCAGIP-3xflag MePOU5F3 (*Macropus eugenii* POU5F3), MeP3 | This paper | Synthetic gene |
| pCAGIP-3xflag RtPOU5F1 (*Rhincodon typus* POU5F1), RtP1 | This paper | Synthetic gene |
| pCAGIP-3xflag RtPOU5F3 (*Rhincodon typus* POU5F3), RtP3 | This paper | Synthetic gene |
| pCAGIP-3xflag LePOU5F1 (*Leucoraja erinacea* POU5F1), LeP1 | This paper | Synthetic gene |
| pCAGIP-3xflag LePOU5F3 (*Leucoraja erinacea* POU5F3), LeP3 | This paper | Synthetic gene |
| pCAGIP-3xflag ScPOU5F1 (*Scyliorhinus canicula* POU5F1), ScP1 | This paper | Synthetic gene |
| pCAGIP-3xflag ScPOU5F3 (*Scyliorhinus canicula* POU5F3), ScP3 | This paper | Synthetic gene |
| pCAGIP-3xflag EbPOU5 (*Eptatretus burgeri* POU5), EbP5 | This paper | Synthetic gene |
| pCAGIP-3xflag EbP5LH2 (Hagfish EbP5-coelacanth LcP1 chimeric protein) | This paper | Synthetic gene |
| pCAGIP-3xflag EbP5S4LH2 (Hagfish EbP5-coelacanth LcP1 chimeric protein) | This paper | Synthetic gene |
| pCAGIP-3xflag S313 (*Scyliorhinus canicula* POU5F1-POU5F3 chimeric protein) | This paper | Synthetic gene |
| pCAGIP-N25P91C25 (*Xenopus laevis* POU5F3.2/POU25- POU5F3.1/POU91 chimeric protein) | This paper | Synthetic gene |

| | | |
|---|---|---|
| pCAGIP-N91P25C91 (*Xenopus laevis* POU5F3.2/POU25- POU5F3.1/POU91 chimeric protein) | This paper | Synthetic gene |
| pMXs-mOct4 (*Mus musculus*) | Takahashi and Yamanaka, 2006 (ref[85]) | Addgene |
| pMXs-3xflag Xlpou91 (*Xenopus laevis* POU91)/XlPOU5F3.1, X91 | This paper | cDNA from Morrison and Brickman 2006 |
| pMXs-3xflag Xlpou25 (*Xenopus laevis* POU25)/XlPOU5F3.2, X25 | This paper | cDNA from Morrison and Brickman 2006 |
| **Deposited Data** | | |
| DNA microarray data of LcPOU5F1, LcPOU5F3 and mOct4-rescued ESCs | This paper | GSE148167 |
| DNA microarray data of X91 SKM iPSCs, X25 SKM iPSCs and mOct4 SKM iPSCs | This paper | GSE183049 |
| **Software** | | |
| BEAST | Suchard et al., 2018 (ref[86]) | https://beast.community |
| MrBayes | Huelsenbeck and Ronquist, 2001 (ref[87]) | https://nbisweden.github.io/MrBayes/index.html |
| CellProfiler | Stirling et al., 2021 (ref[69]) | https://cellprofiler.org |
| NIA Array Analysis Tool | Sharov et al., 2005 (ref[70]) | https://lgsun.grc.nia.nih.gov/ANOVA/ |
| Morpheus | Morpheus, https://software.broadinstitute.org/morpheus (ref[80]) | https://software.broadinstitute.org/morpheus |
| PANTHER Classification System | | http://pantherdb.org |
| ShinyGO v0.61 | Ge et al., 2019 (ref[71]) | http://bioinformatics.sdstate.edu/go/ |
| Fiji ImageJ | Schindelin et al., 2012 (ref[68]) | https://imagej.net/Fiji |
| AlphaFold2 | Jumper et al., 2021 (ref[48]) | https://colab.research.google.com/github/deepmind/alphafold/blob/main/notebooks/AlphaFold.ipynb#scrollTo=rowN0bVYLe9n |
| PyMol | The PyMOL Molecular Graphics System, Version 2.0 Schrödinger, LLC. (ref[73]) | https://pymol.org/2/ (version 2.5.1) |
| Phenix (Python-based Hierarchical ENvironment for Integrated Xtallography) | Liebschner et al., 2019 (ref[74]) | https://phenix-online.org/download/ |
| UCSF ChimeraX version: 1.2 | Pettersen et al., 2021 (ref[76]) | https://www.rbvi.ucsf.edu/chimerax/ |
| **Other resources** | | |
| Resource website for *POU5* genes | NCBI genome database; Ensembl | https://www.ncbi.nlm.nih.gov/genome; https://m.ensembl.org |
| Resource website for Chondrichthyes *POU5* genes | Squalomix database | https://transcriptome.riken.jp/squalomix/blast/ |
| Resource website for little skate *POU5* genes | SkateBase | http://skatebase.org/ |

| | | |
|---|---|---|
| Resource website for thorny skate *POU5* genes | GenomeArk | https://vgp.github.io/genomeark/Amblyraja_radiata/ |
| Resource website for lamprey *POU5* genes | Stowers Institute | https://genomes.stowers.org/organism/Petromyzon/marinus |
| Resource website for mouse Oct4 structure on *PORE* DNA | PDB: 3L1P (ref[49]) | Esch et al., 2013 DOI: 10.2210/pdb3L1P/pdb |
| Resource website for mouse Sox2 structure | PDB: 6HT5<br>PDB: 1GT0 (ref[52]) | Remenyi et al., 2003 DOI: 10.2210/pdb6HT5/pdb<br>DOI: 10.2210/pdb1GT0/pdb |
